# PanVasc Research for AI assisted evidence analysis in panvascular intervention

**DOI:** 10.64898/2026.09.22.752857

**Authors:** Lingsen You, Yujun Guo, Wentong Wang, Zisu Peng, Xinyu Zhong, Li Shen, Junbo Ge

**Affiliations:** Department of Cardiology, Zhongshan Hospital, Fudan University, Shanghai Institute of Cardiovascular Diseases, Shanghai 200032, China; National Clinical Research Center for Interventional Medicine, Shanghai 200032, China; State Key Laboratory of Cardiovascular Diseases, Zhongshan Hospital, Fudan University, Shanghai 200032, China; University of Shanghai for Science and Technology (USST), Oriental Pan-Vascular Devices Innovation College, Shanghai 200093, China; College of Biomedical Engineering, Fudan University, Shanghai 200433, China

**Author notes:** Correspondence to Li Shen and Junbo Ge. Lingsen You, Yujun Guo, Wentong Wang, Zisu Peng, and Xinyu Zhong contributed equally to this work.

**Keywords:** panvascular intervention, artificial intelligence, evidence provenance, clinical trial registry, natural language inference, research software

## Abstract

Panvascular intervention research requires evidence workflows that preserve source identity, outcome definitions and observation windows. We developed PanVasc Research, an executable research framework, and evaluated a fixed local Qwen3-4B model using two complementary tasks. Fifty ClinicalTrials.gov records from five vascular query strata generated 200 source-fidelity tests with intact evidence or controlled removal of the requested source, primary outcome or timeframe, plus 50 clean controls. A separate 100-statement sample from the official NLI4CT test set assessed clinical-trial entailment and evidence selection against existing expert labels. Generic and checklist prompts used identical evidence and maximum generation budgets; each output was also evaluated with an input-only deterministic contract. Registry exact accuracy was 51/200 (25.5%) with the generic prompt and 77/200 (38.5%) with the checklist; intact-case agreement was 50/50 and 48/50. The rule baseline recovered 200/200 tuples. NLI label accuracy was 48/100 (48.0%) and 50/100 (50.0%), respectively. The source contract retained 35 and 31 incorrect NLI labels in the two arms.

Registry references establish fidelity to a registration snapshot, while NLI4CT concerns breast-cancer trials and does not validate vascular expertise. The framework also retains provenance-recorded literature retrieval, structured research planning and local numerical analysis. These experiments support a bounded assessment of source handling and semantic failure, rather than a new foundation model or autonomous scientific discovery. A proposed endpoint representation identifies the additional domain annotation and independent validation required for a panvascular research model.

## Introduction

Panvascular intervention connects vascular territories through shared devices and research methods while retaining differences in disease, anatomy and outcome assessment. A device-vessel research perspective motivates shared infrastructure without making coronary, limb, carotid, aortic and venous outcomes interchangeable [1]. Research on panvascular experimental systems and imaging similarly connects measurements across scales while preserving their context [7,8]. A useful research assistant must retain where an observation came from, how it was defined and which conclusions it can support.

Endpoint interpretation requires more than recognizing a familiar name. Reporting standards distinguish endpoint components, procedural context and observation rules across vascular specialties [2-6]. A patient-level risk cannot be substituted for a lesion-level proportion, and an event within a follow-up interval is not equivalent to a measurement at a scheduled visit. An assistant that reproduces the endpoint name while dropping these restrictions can generate a plausible but misleading evidence table. Correct source binding is a necessary first step, although it cannot by itself resolve clinical interpretation.

Scientific agents already connect literature, tools and researcher interaction. ScienceBuddy combines optimization of an agent’s operating procedures with model reinforcement learning [9]. TrialMind and LEADS address medical literature search and extraction, while C-TrO-based research represents relations between trial arms, treatments, endpoints and outcomes [17-19]. MESHAgents provides a cardiovascular research-agent precedent involving cardiac and aortic imaging phenotypes [20]. These studies preclude a broad novelty claim based merely on a vascular interface, structured output or agent orchestration.

We developed PanVasc Research to examine a narrower question: how should a small-model evidence workflow separate source fidelity from semantic support? We compare two fixed prompts, retain a deterministic lookup baseline, and measure the consequences of applying an input-only output contract. A vascular registry task tests exact field extraction and controlled missingness. An independently defined NLI4CT task tests clinical-trial entailment and evidence selection in a different disease domain [21]. The surrounding software records provenance, supports structured research questions and performs local numerical analyses. We also propose a richer endpoint representation, while separating this future design from the capabilities actually evaluated.

## Methods

### Framework and evaluation boundaries

The framework records a task’s vascular domain, question, source material, analysis inputs, outputs and reviewer decisions. Five configurations cover coronary, peripheral arterial, carotid, aortic and venous research; these are workflow configurations rather than validated specialty expertise. The carotid configuration does not cover all neurointervention, and the venous configuration does not cover all venous procedures. The architecture connects the original deterministic tools to a separate local-model evaluation workflow (Figure 1). The benchmark runner, rather than the desktop interface, executes the reported model experiments.

**Figure 1.**
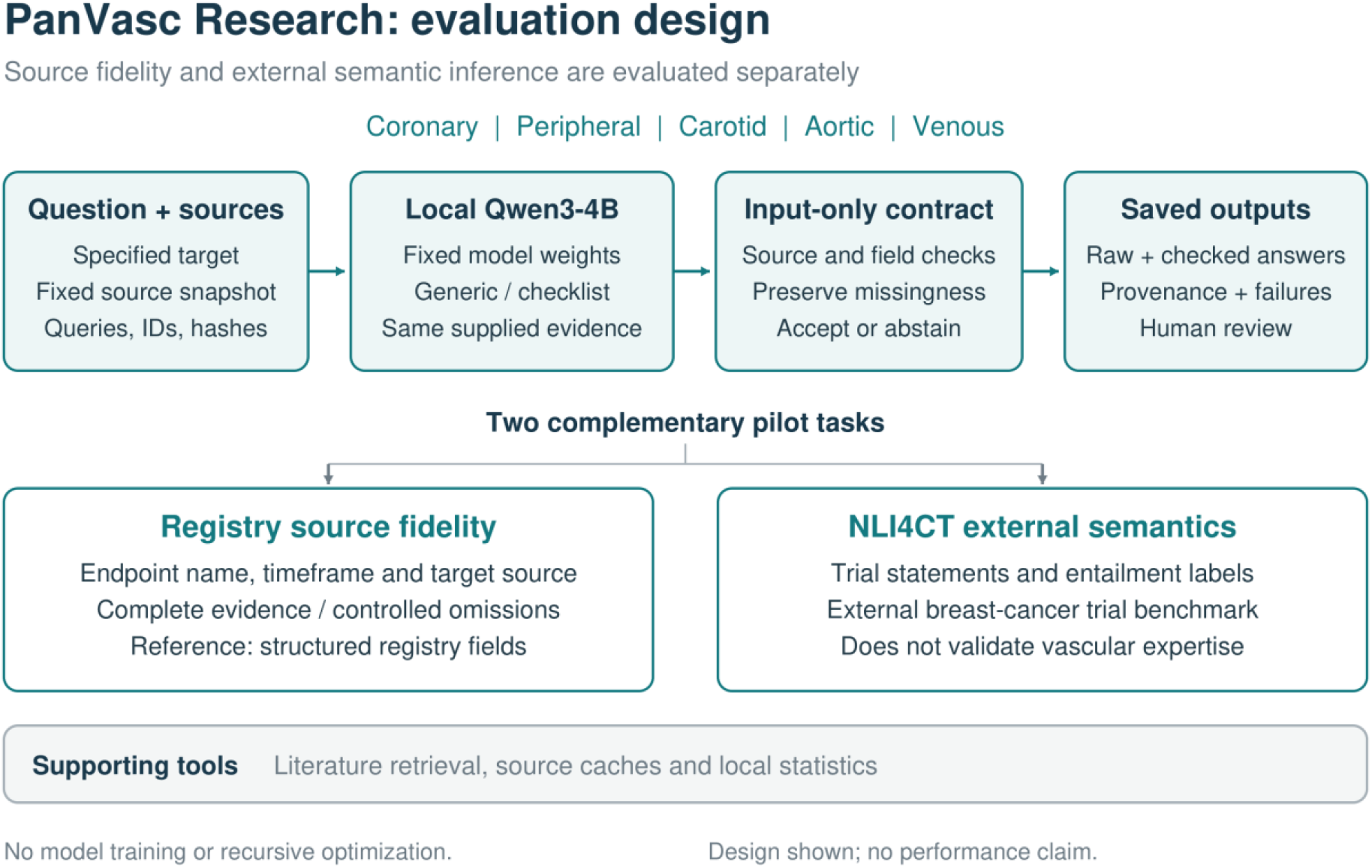
PanVasc Research evaluation architecture. The fixed local model receives a question and source snapshot, followed by input-only contract checks and saved outputs. Registry source fidelity and external clinical-trial semantics are evaluated separately. Human review is an intended workflow step, not a recruited panel in this study. No model training or recursive optimization was performed.

The implemented source contract checks whether an output meets explicit conditions recoverable from the supplied input. For registry extraction these conditions include the target record, original outcome index, exact measure and timeframe, and missing-field state. For NLI4CT they include output shape, allowed label, referenced trial identities and evidence-index validity. The latter contract cannot decide whether a referenced line supports the conclusion. Rejected outputs are retained in the audit record and counted as incorrect in unconditional accuracy; the guard does not replace them with reference answers.

Figure 2 describes a proposed four-layer endpoint representation: endpoint definition, observation rule, analysis context and estimand evidence. Components and thresholds belong to the first layer; time anchors and windows to the second; units, denominators and event handling to the third; and explicitly reported treatment-effect attributes to the fourth. This proposal is not an implemented clinical ontology or a validated compatibility classifier. An endpoint is only one attribute of an estimand, and an intention-to-treat label does not fully specify all intercurrent-event strategies [24]. The current experiments test source fields and externally defined inference tasks, not these four clinical layers.

**Figure 2.**
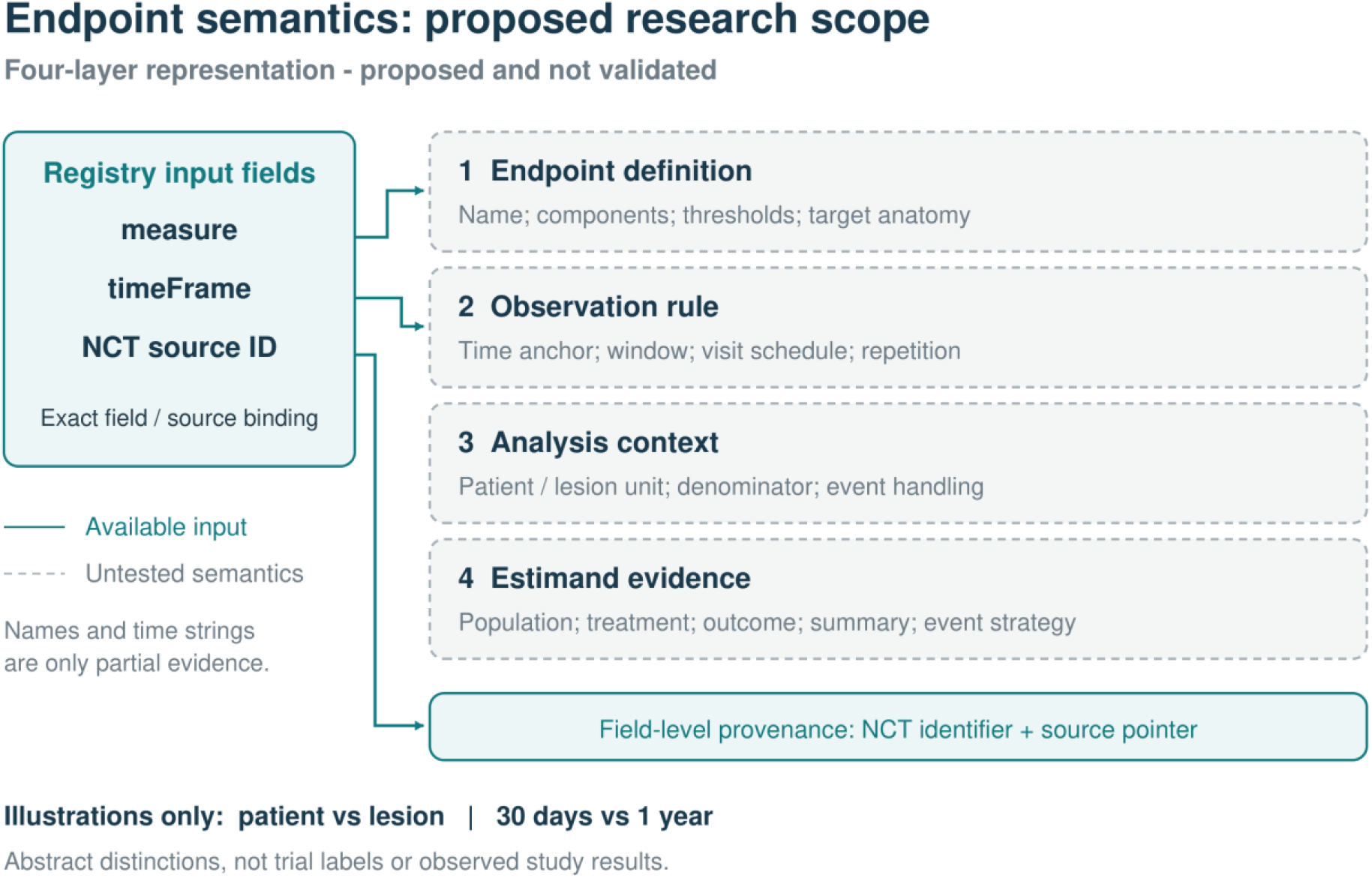
Proposed endpoint representation and the scope of current source fields. Exact registry measure and timeframe strings provide partial evidence for endpoint definitions and observation rules. The four clinical layers remain proposed and unvalidated. Patient versus lesion and 30 days versus one year are abstract illustrations, not observed trial labels. Source-field agreement does not establish clinical comparability or suitability for pooling treatment effects.

### Vascular registry material

We queried the ClinicalTrials.gov version 2 API on 19 September 2026 using five predefined vascular search strata [23]. Each query requested at most 100 records from its first response page. After NCT deduplication, 498 candidates remained. Eligibility required an interventional study, at least one DEVICE or PROCEDURE intervention, and at least one primary outcome with nonempty measure and timeFrame. This automatic rule does not establish expert-adjudicated vascular relevance. After applying these field-based criteria and excluding previously viewed pilot records, 246 records remained eligible. Seeded sampling selected two development and ten test records per stratum. An exposure addendum documented that the previously reviewed ATTRACT identifier was absent from the candidate pool. The 10 development and 50 test NCTs do not overlap.

The retained snapshot contains trial identifiers, titles, conditions, interventions and ordered outcome fields, without contacts or locations. A separate extraction directly from each API response supplies reference field values. Response and field hashes, queries, timestamps, selection logic and identifiers are recorded. The reference establishes what the registry snapshot reported, not medical truth, published trial results or endpoint appropriateness.

For each test record we selected its first primary outcome with both required fields. The query specifies only its NCT and original zero-based outcome index. The evidence packet contains that record and two same-stratum test distractors, with seeded ordering. Development packets have one available same-stratum distractor. Packets contain titles and numbered primary outcome measures and timeframes; descriptions and secondary outcomes are omitted. No NCT appears across development and test, including as a distractor.

Four paired conditions retain the intact packet or delete the requested source, requested outcome, or requested timeframe. Deleted values are not placed in the query, and outcome indices are not renumbered. The output schema requires an availability state, source identifier, outcome index, measure and timeframe, preserving unavailable fields as null. These controlled deletions create 200 tests from 50 target records. They do not estimate naturally occurring missingness. An additional 50 intact single-source controls distinguish source copying from distractor handling. A deterministic packet lookup is an explicit baseline because the requested fields are structurally specified (Figure 3).

**Figure 3.**
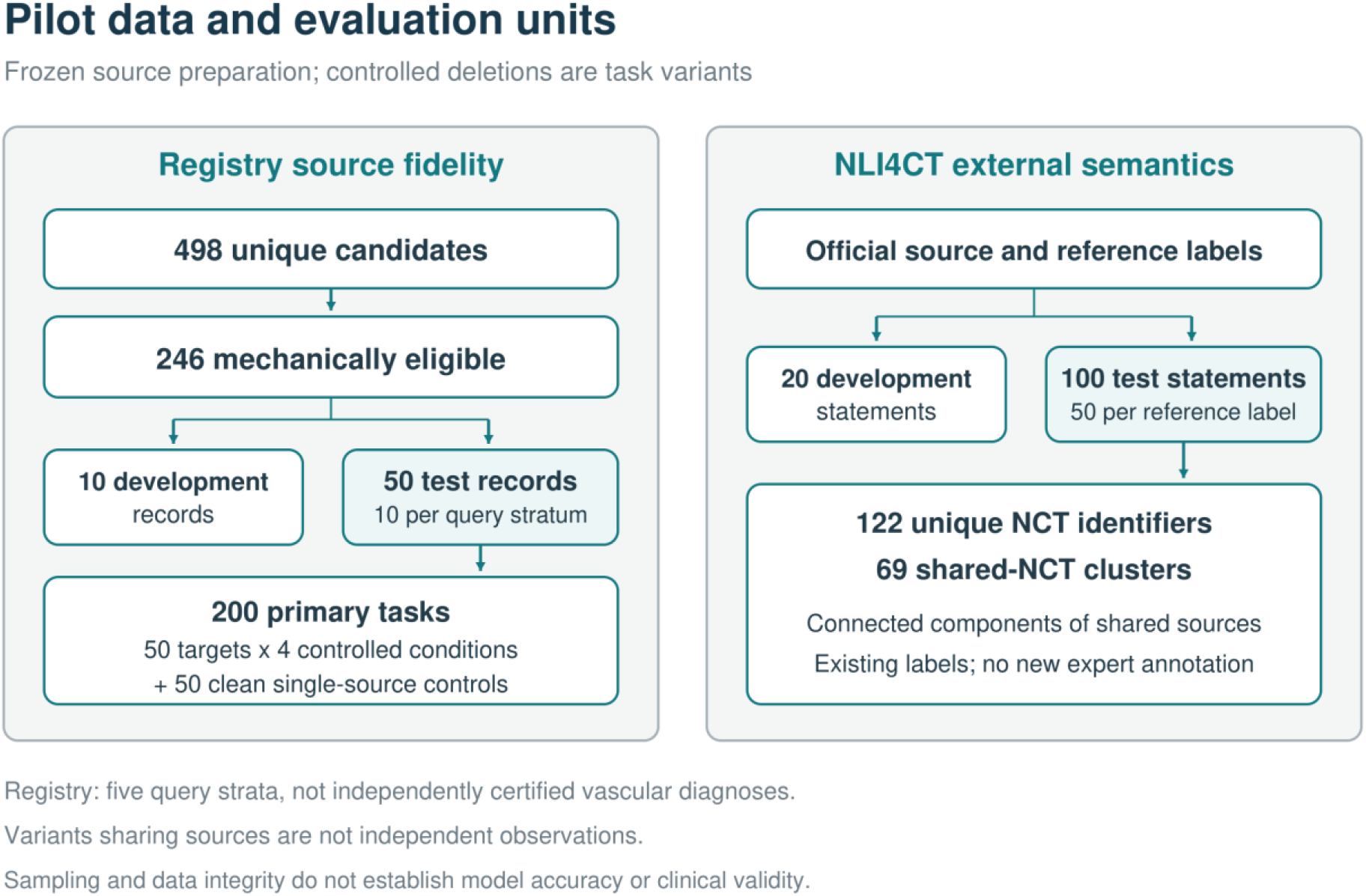
Source material, partitions and task construction. The registry cohort is sampled from first-page query results after automatic eligibility and development-exposure exclusions. Query strata are not expert labels. The 200 controlled tasks comprise four conditions per target; 50 clean controls are additional. NLI4CT labels and evidence originate from its official release. Neither dataset supplies newly adjudicated panvascular semantic labels.

### External semantic task

NLI4CT provides clinical-trial statements, relevant trial sections, entailment or contradiction labels, and evidence-line annotations [21]. We used the official repository at revision 8e8e5fc506b916984011873403b9ea5ed6dd96c2. Fixed-seed, label-stratified sampling selected 20 statements from the official development set and 100 from the official test set, with the latter’s original gold labels stored separately. The test sample contains 50 statements of each label, 39 single-trial statements and 61 comparisons, involving 122 unique NCTs. No development NCT occurs in the test sample. Shared primary or secondary NCTs connect the 100 statements into 69 components, with at most four statements per component.

The model receives the statement and all indexed lines of the specified section for each associated trial. It returns a label and an evidence-index list for each supplied trial. The reference labels and evidence annotations are inherited from the original expert-annotated dataset; this study did not recruit new annotators. The dataset concerns breast-cancer trials and supplies an external semantic task rather than external vascular validation. Because it is public and predates the model, pretraining contamination cannot be excluded.

### Model and comparison conditions

We used the official Qwen3-4B-GGUF Q8_0 checkpoint at revision bc640142c66e1fdd12af0bd68f40445458f3869b [22]. Local inference used llama.cpp build b10964, commit b29c606e28a01b1bc8c1351026a0fa6e616bf6c4, with the Vulkan backend on an NVIDIA GeForce RTX 5070 Ti with 16 GB graphics memory. The server was bound to the local loopback interface, used an 8192-token context and one concurrent slot, and received no participant-level records. The pilot runner and scoring used Python 3.12.14 and NumPy 2.3.5, separately from the earlier software-test environment. Model and runtime hashes are included in the reproducibility material.

Both prompts share the task definition, exact output schema, evidence packet and instruction to treat source text as data. The generic arm asks for the requested output directly. The checklist arm additionally directs the model to verify record identity, field or section location, quantities or negations, and the treatment of missing evidence before returning JSON. No demonstrations, external search, self-revision or additional model calls are used. Thinking mode is disabled, temperature is zero, top_p is one, the seed is 20260919, and the maximum generated length is 512 tokens. This deterministic pilot setting differs from the model card’s general sampling recommendation. Both arms have equal output limits; actual token use may differ.

Prompts, scoring code, input files and reference files were hashed before model evaluation. This was a local analysis freeze, not a public preregistration. A technical development check executed 10 calls on reserved development inputs before the 700-call test run. No development accuracy claim is made; its purpose was to verify request and output handling. The frozen prompts and scoring rules were not changed after these calls. Test calls were interleaved in a seeded shuffled order. Each task and arm received one call; raw outputs, errors, token counts, finish reasons and elapsed times were recorded. There was no retry for incorrect or malformed content and no prompt change in response to test results.

Each original model output is evaluated with and without the deterministic contract, yielding a two-by-two comparison without extra inference. JSON parsing accepts an optional outer code fence but not an arbitrary recoverable substring. Both raw and guarded analyses require the prespecified keys and types; boolean or floating-point values do not satisfy an integer-index field. The NLI contract additionally requires exactly the supplied trial identities, unique in-range integer indices and nonempty evidence overall. It does not inspect reference labels or annotated evidence. A valid source or index therefore remains compatible with a wrong semantic judgment.

### Outcomes and statistics

The primary registry measure is exact agreement of the complete output tuple with the source-field reference and logged deletion. We average the four conditions within each target NCT, then give NCTs equal weight. Intact, missing-source, missing-outcome and missing-time results are also reported separately, because the artificial three-to-one missingness mixture must not obscure intact performance. Clean controls are excluded from the primary denominator. Source and field errors, retained coverage and accuracy among retained outputs are secondary measures. Coverage denotes a retained schema-valid response, including a valid unavailable-field state; it does not mean that a complete endpoint value was available.

The primary NLI measure is unconditional label accuracy. Secondary measures are macro label F1, evidence precision, recall and F1, exact evidence agreement, malformed-output frequency, contract rejection and cost. Evidence is scored as sets of trial-index pairs. True positives, false positives and false negatives are aggregated across all 100 statements for micro evidence F1. Missing outputs contribute false negatives, repeated indices cannot add true positives, and a zero denominator gives zero. Evidence agreement measures correspondence to the dataset annotations rather than completeness of every possible supporting rationale. The balanced majority-label baseline has 50% label accuracy and no evidence output.

Paired checklist-minus-generic differences and accuracy intervals use 2000 bootstrap resamples with seed 20260919. Registry resampling retains all four conditions within each target NCT. NLI resampling retains statements within shared-NCT connected components and computes example-weighted accuracy. These 95% percentile intervals characterize the convenience samples. Registry distractor reuse and potentially related registrations leave dependencies beyond target-NCT grouping; small stratum samples and unknown pretraining exposure further limit inference. No prospective power calculation, multiplicity-adjusted superiority test or cross-paper score comparison was performed.

### Supporting software demonstrations

The original workflow retrieves metadata through NCBI E-utilities or an explicitly recorded Europe PMC route [10,11]. Query semantics differ between these services, and requests are capped at 25 records for exploratory use. Identifiers, actual queries, response hashes, missing fields and transport failures are preserved. Identifier presence and available retraction flags are checked separately from semantic support. These functions do not implement a complete systematic review.

Structured research planning requires population, intervention, comparator, outcome and follow-up fields and returns a template with unresolved investigator decisions. Local CSV tools check missingness, repeated identifiers, binary outcomes, selected leakage patterns and supplied split overlap. They calculate descriptive summaries, crude binary risks with Wilson intervals and a two-sided Fisher exact test using SciPy [12]. They do not model censoring, confounding or repeated observations. The existing remote API adapter [13] was not exercised in this evaluation; it is distinct from the new local-model runner.

The supporting development tests and earlier retrieval demonstrations used Python 3.11.9, NumPy 2.4.6, pandas 2.3.3 and SciPy 1.17.1. They comprise 28 development-time checks, five-domain live metadata retrieval and a fully synthetic 120-row numerical example. They are engineering demonstrations, separate from the frozen model comparisons. Task-linked accept, revise and reject feedback is stored with content hashes; no real investigator panel, automatic procedure search or weight update was performed.

## Results

### Material and execution

All 50 registry targets and 100 NLI statements entered the test evaluation. The two prompt arms generated 500 registry calls, including 100 clean-control calls, and 200 NLI calls, for 700 calls in total. There were 0 transport failures and 0 responses that reached the generation limit; all remained in the denominators. Recorded generation comprised 33,695 completion tokens, with 6.0 minutes of summed request time. This is an observed local deployment cost, not a standardized hardware benchmark. The registry acquisition and NLI component structure are shown in Figure 3. All 50 selected registry targets happened to use original primary outcome index zero; arbitrary-index extraction was therefore not assessed. Requests used an already loaded server with prefix caching; elapsed times do not describe cold-start or standardized hardware performance.

### Registry source fidelity

Exact tuple agreement across the 200 controlled tests was 51/200 (25.5%) for the generic prompt and 77/200 (38.5%) for the checklist (Figure 4). The paired difference was +13.0 percentage points (95% cluster bootstrap CI +9.5 to +16.5). By condition, the generic arm achieved intact 50/50, source removed 0/50, outcome removed 1/50, timeframe removed 0/50; the checklist achieved intact 48/50, source removed 29/50, outcome removed 0/50, timeframe removed 0/50. In the separate single-source controls, the arms achieved 50/50 and 50/50, respectively. The deterministic lookup recovered 200/200 tuples. Its result demonstrates that this structurally specified task can be solved without a language model; it is not an estimate of clinical endpoint understanding.

**Figure 4.**
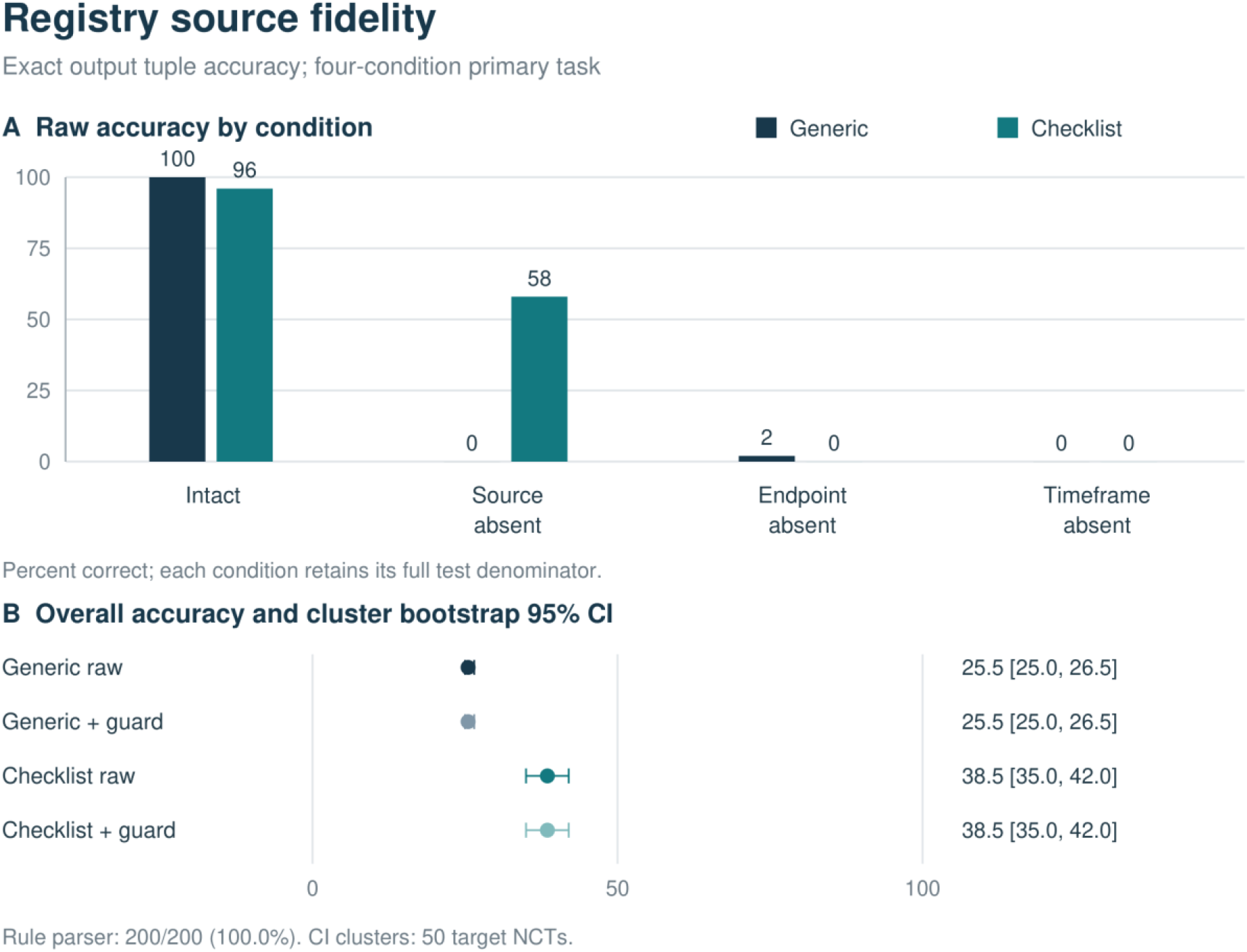
Registry source-field extraction. Generic and checklist prompts are evaluated on the same 50 target records under four controlled conditions. Exact accuracy requires the complete source-bound tuple and missingness state. The primary result weights the four conditions equally within each NCT; intervals use 2000 target-NCT bootstrap resamples. Distractor reuse creates additional dependencies. The rule baseline accesses the same explicit fields directly, and the 50 clean controls are separate from the main denominator.

Exploratory inspection after the primary scoring showed that strict tuple errors did not all represent fabricated clinical content. With the timeframe removed, 30/50 generic and 35/50 checklist outputs correctly left the time value null, despite zero fully correct tuples. In 30 and 31 cases, respectively, availability was the only incorrect field. The other 20 and 15 non-null time outputs included inferred intervals and missing-value text; these violate the source-field contract but are not all proven fabrications of clinical facts. Missingness-state consistency and unsupported content completion should therefore be distinguished.

### Clinical trial inference and evidence selection

Unconditional NLI label accuracy was 48/100 (48.0%) with the generic prompt and 50/100 (50.0%) with the checklist (Figure 5). The paired difference was +2.0 percentage points (95% cluster bootstrap CI -5.1 to +8.4), using the 69 shared-NCT components. Macro label F1 was 0.505 and 0.534. Evidence micro F1 was 0.253 and 0.286; exact evidence-set agreement occurred in 6/100 and 5/100 statements. The balanced majority-label comparator achieved 50/100 label accuracy. These results concern the sampled breast-cancer statements and do not establish vascular-domain performance. There were 10 malformed JSON responses in the generic arm and 13 in the checklist arm; all occurred on comparison statements. They remained incorrect in the primary analysis and also account for the schema failures, rather than forming an additional failure category. None reached the 512-token generation limit.

**Figure 5.**
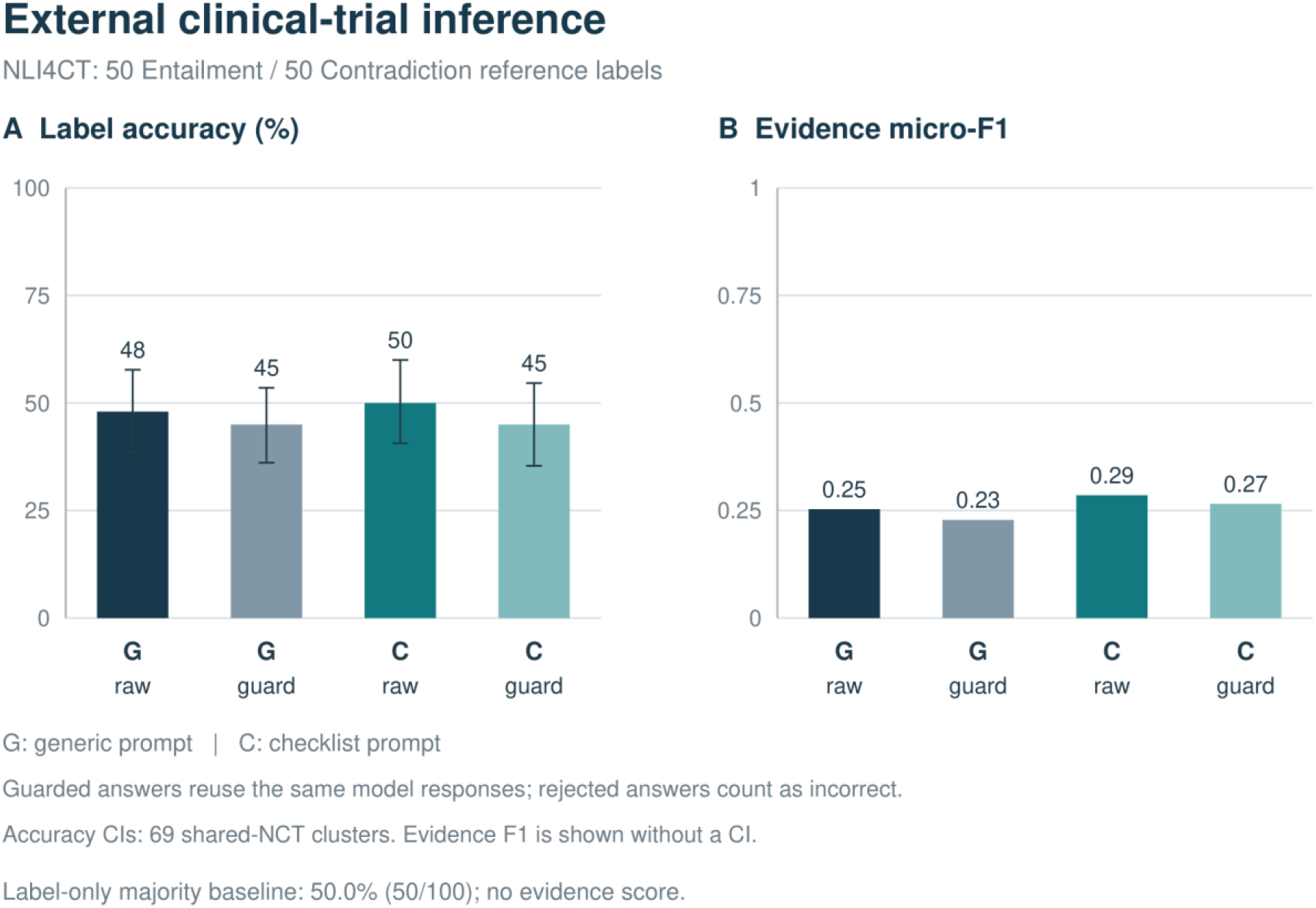
NLI4CT semantic inference and evidence selection. Label accuracy uses all 100 statements, including malformed or rejected outputs; accuracy intervals resample 69 components connected by shared trial identities. Evidence F1 aggregates trial-index agreement across all statements. Raw and guarded results reuse identical generated answers. This breast-cancer benchmark is an external semantic task and does not validate vascular expertise.

### Contract coverage and failure boundaries

For registry with the generic prompt, the contract retained 51 outputs, including 0 incorrect answers, and rejected 149 outputs, including 0 answers that were correct on the raw primary metric. For registry with the checklist prompt, the contract retained 77 outputs, including 0 incorrect answers, and rejected 123 outputs, including 0 answers that were correct on the raw primary metric. For nli4ct with the generic prompt, the contract retained 80 outputs, including 35 incorrect answers, and rejected 20 outputs, including 3 answers that were correct on the raw primary metric. For nli4ct with the checklist prompt, the contract retained 76 outputs, including 31 incorrect answers, and rejected 24 outputs, including 5 answers that were correct on the raw primary metric. Figure 6 shows these paired outcomes. Guarded accuracy includes rejected outputs as incorrect, whereas retained-answer accuracy uses only accepted outputs. Registry acceptance is checked against explicit packet fields; zero retained tuple errors, if observed, follows from that narrow contract. NLI source and index checks do not verify the semantic label. Table 2 reports primary accuracy and coverage together.

**Table 1.** Evaluation scope and comparison with related work. The distinctions concern reported study design, not rankings or cross-benchmark performance.

| Work or component | Relevant capability | Relationship to the present study |
| --- | --- | --- |
| ScienceBuddy [9] | Recursive procedure optimization and model training | Inspiration for separating procedures from weights; not reproduced here |
| TrialMind and LEADS [17,18] | Medical literature search and extraction | Prior evidence workflows; no matched comparison performed |
| C-TrO and MESHAgents [19,20] | Relational trial extraction and cardiovascular research agents | Prior structured and specialty research systems |
| Registry pilot | Fixed model, controlled source omissions and rule baseline | Tests source-field fidelity, not medical reasoning |
| NLI4CT pilot | Existing expert labels and evidence indices | Tests external breast-cancer trial semantics, not vascular validity |
| Proposed endpoint representation | Four clinical layers and source provenance | Design proposal requiring independent domain annotation |

**Table 2.** Primary correctness and retained coverage. Registry denominators are 200 controlled tasks from 50 target NCTs; NLI denominators are 100 statements in 69 source-sharing components. Coverage counts schema-valid structured responses, including unavailable-field states, and does not mean that a complete endpoint is available. Rejected outputs remain incorrect in unconditional accuracy. A rejection-only guard cannot increase the number of correct outputs and can remove a correct NLI label with invalid evidence. The same model outputs are used for raw and guarded rows.

| Task and prompt | Output | Correct / total | Coverage | Correct among retained |
| --- | --- | --- | --- | --- |
| Registry generic | raw | 51/200 | 100.0% | 25.5% |
| Registry generic | guard | 51/200 | 25.5% | 100.0% |
| Registry checklist | raw | 77/200 | 100.0% | 38.5% |
| Registry checklist | guard | 77/200 | 38.5% | 100.0% |
| NLI4CT generic | raw | 48/100 | 90.0% | 53.3% |
| NLI4CT generic | guard | 45/100 | 80.0% | 56.2% |
| NLI4CT checklist | raw | 50/100 | 87.0% | 57.5% |
| NLI4CT checklist | guard | 45/100 | 76.0% | 59.2% |

**Figure 6.**
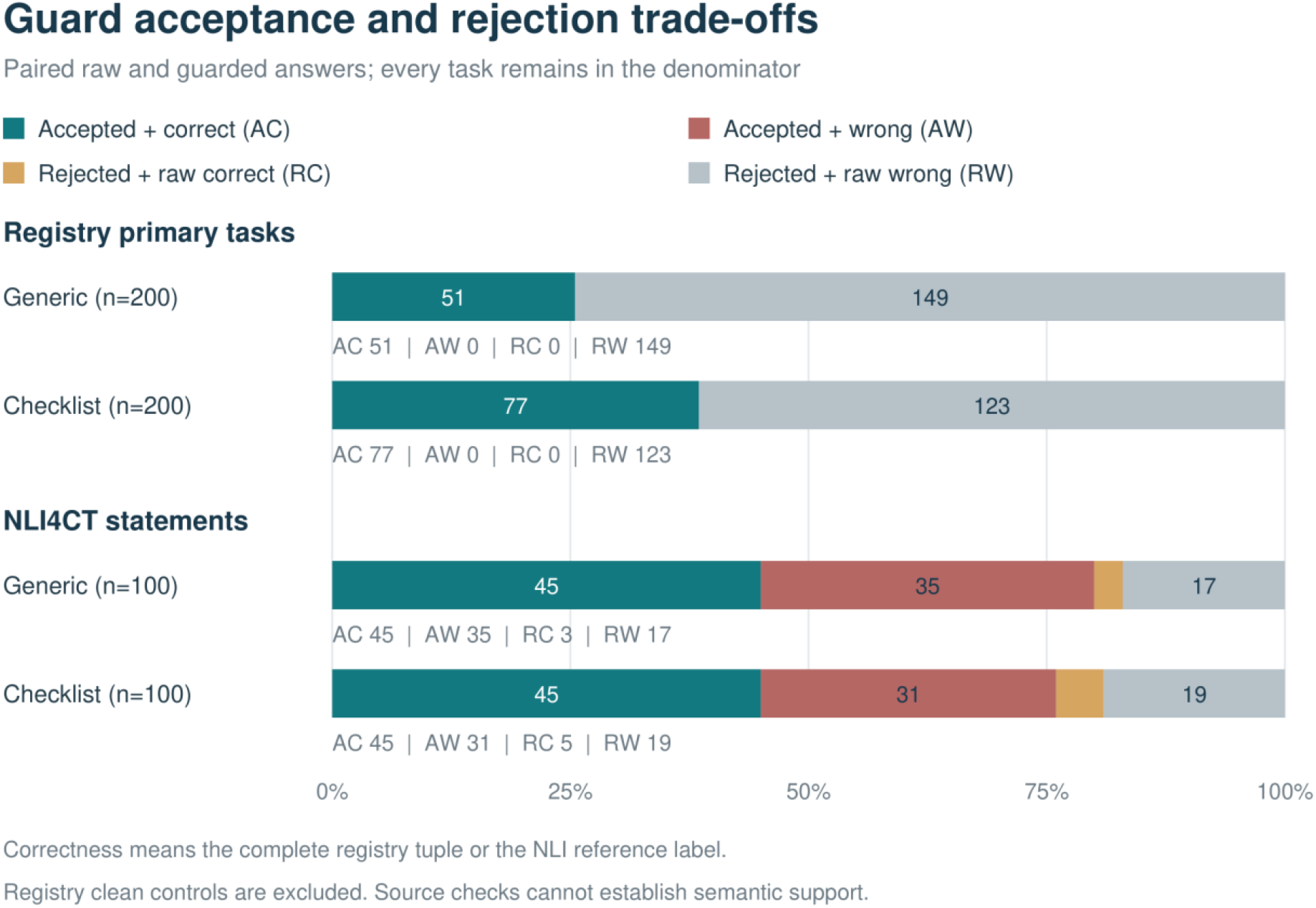
Accepted errors and the cost of rejection. Each task is classified using its original primary score and subsequent contract decision. Registry correctness is exact tuple agreement; NLI correctness is the label only, so a correct raw label may still carry invalid evidence. Source checks can reject outputs while preserving semantic errors among accepted NLI answers. The registry panel excludes clean controls. Counts depict the observed sample, not clinical risk rates.

### Supporting workflow behavior

The original acceptance suite passed 28 of 28 checks after development corrections to blank identifiers, missing outcomes and equivalent numeric identifiers. These are selected regression fixtures, not independent generalization tasks. Initial NCBI requests failed in all five domains because of transport errors; an initial Europe PMC run completed three, and a later run completed all five, returning 15 records with nonempty identifiers and titles. All five final requests succeeded on their first attempt, so the three-to-five change cannot establish a causal benefit of retries. Retrieval relevance and coverage were not adjudicated.

The synthetic dataset contained 60 records per group, with constructed event counts of 8 and 12 and three missing age values. The program reported risks of 13.33% and 20.00%, a crude risk difference of -6.67 percentage points and a two-sided Fisher P value of 0.46316. These calculations validate selected numerical behavior and support no biological or treatment-effect conclusion. The registry and NLI experiments, rather than these software checks, supply the model-specific observations in the revised study.

## Discussion

The paired pilot found a checklist-minus-generic difference of +13.0 percentage points (95% cluster bootstrap CI +9.5 to +16.5) for registry source fidelity and +2.0 percentage points (95% cluster bootstrap CI -5.1 to +8.4) for NLI label accuracy. These are task-specific observations from one small model and one frozen sample. The deterministic registry baseline and the 35 and 31 incorrect NLI labels retained by the source contract show why field access, citation validity and semantic support must be evaluated separately. The results do not justify transferring performance claims from source copying to clinical interpretation. The registry improvement was concentrated in absent-source handling; intact agreement decreased from 50/50 to 48/50, and neither arm produced a fully correct missing-time tuple. NLI raw accuracy did not exceed the 50% majority reference, and its paired interval included zero. These findings support a failure analysis and a limited source-handling effect rather than a useful semantic capability or panvascular advantage.

The registry task has a deliberately narrow meaning. Exact field agreement establishes fidelity to the supplied registration snapshot. It does not show that the endpoint is appropriate, matches a published report, or is clinically comparable with another trial. A deterministic lookup is therefore the relevant simplest baseline. A model should not receive credit for medical reasoning merely because it reproduces a field obtainable through direct indexing. This task instead makes source confusion and behavior under unavailable evidence observable.

The NLI task exposes a different boundary. A well-formed answer can cite genuine, in-range lines and still misunderstand a comparison or endpoint statement. The source contract has no access to the expert label and cannot resolve that semantic mistake. Retained coverage must therefore be interpreted together with errors among accepted outputs. An output guard that improves apparent reliability by withholding answers has a different practical effect from a model that produces more correct answers. Neither should be described as clinical validation.

The work takes inspiration from ScienceBuddy’s distinction between operating procedures and model weights, but does not reproduce its training experiments [9]. We compare two manually specified prompts around one fixed model. We do not optimize candidate procedures, learn from expert trajectories, perform reinforcement learning or test autonomous scientific discovery. ScienceBuddy’s base model, task distribution, compute allocation and outcome definitions differ, so its published scores cannot serve as numerical comparators. The contribution here is a small, inspectable domain-oriented evaluation and a documented failure boundary, not equivalent experimental scale.

The proposed endpoint representation identifies where meaningful panvascular specialization could be tested. Clinical reviewers would need to annotate components, thresholds, time anchors, analysis units and estimand evidence in actual reports, preserving both absent information and unresolved disagreements. Even full endpoint agreement would identify only a candidate for comparison, not establish that treatment effects may be pooled. Population, intervention, study design, follow-up and intercurrent-event handling remain relevant. C-TrO and established reporting standards provide prior foundations rather than evidence that these functions have already been implemented here [2-6,19,24].

The device-vessel, FLOW and imaging work indexed on the authors’ research website motivates retaining biological context across computational and experimental workflows [1,7,8]. AI-assisted PLLA modifier generation similarly illustrates a separate setting where candidate generation and model-based screening require their own experimental confirmation [16]. These connections explain the intended research program; they are not validation data for the present model.

The next decisive evaluation should use independently annotated vascular reports, with entire trial families withheld from development. Same-model, same-material comparisons should isolate domain rules, source verification and model adaptation. Outcomes should include complete endpoint-tuple accuracy, unsupported completion, mistaken compatibility judgments and useful coverage. A subsequent investigator study could evaluate final extraction quality and time saved. Prediction of patient outcomes, if later added, would require distinct external validation, calibration and risk-of-bias assessment [14,15]. None of those clinical or user-benefit outcomes is established by the present pilot.

### Limitations

First-page registry sampling is a convenience design, and query strata were not adjudicated by vascular specialists. Fifty target records do not represent the diversity of panvascular intervention; copied registry fields are not an expert semantic gold standard. Controlled deletions deliberately enrich missing evidence and cannot estimate its prevalence. NCT-level separation prevents identical registrations crossing partitions but does not establish independence of related trial families. Distractor reuse limits the interpretation of registry confidence intervals.

The semantic sample is small, comes from breast-cancer trials, and uses public historical data with possible pretraining exposure. Only one model, one quantization, two manually written prompts and one generation seed were assessed. There was no model-family replication, clinical investigator evaluation, blinded domain adjudication or randomized usability study. Guarded registry fidelity follows partly from the exact fields that the guard verifies, while NLI index validity has no semantic guarantee. Model results therefore do not establish a leading panvascular system, a new general-purpose foundation model, or equivalence to ScienceBuddy.

The initial network demonstrations were adaptive and their historical transport conditions cannot be reproduced exactly. Registry text may change after retrieval. Complete third-party text is excluded from the shareable supplement where redistribution terms were not verified, so source identifiers, hashes and retrieval scripts are necessary but may not fully reconstruct a mutable historical registry snapshot. The package documents these limits instead of implying an independently replicated benchmark.

## Conclusion

PanVasc Research combines provenance-recorded research tools with a reproducible local-model pilot that separates source fidelity from semantic support. The two prompts achieved 25.5% and 38.5% exact registry agreement and 48.0% and 50.0% NLI label accuracy; source checks retained semantic errors. A clinically meaningful panvascular model requires a further layer of independent endpoint annotation, controlled domain comparisons and investigator evaluation. The present work provides an explicit starting point and testable boundaries for that program.

## Supporting information

Supplementary data and reproducibility code

## Declarations

### Data and code availability

The accompanying bundle includes original implementation and evaluation code, prompts, input identifiers, acquisition and model revisions, file hashes, aggregate and per-task numerical scores, acceptance-test results and synthetic data. No public repository accession or DOI has yet been assigned. Source registry text, NLI4CT trial sections, article abstracts, complete model weights and account information are excluded from the shareable package. NLI4CT can be obtained from its official task repository; no explicit corpus redistribution license was located in the inspected archive. Scripts and manifests document its fixed revision and selection. The registry API is publicly accessible, but changes to records can prevent exact reconstruction of the retained local snapshot.

### Ethics and human data

No participants were recruited and no patient-level clinical records, human tissue or animal data were used. The registry and NLI material consists of public trial-level information; the tabular demonstration is synthetic. No institutional approval or exemption is claimed. Future use of real research records requires the applicable permissions and governance arrangements.

### AI assistance

OpenAI Codex assisted literature investigation, software development, test construction, analysis scripting, figure preparation and manuscript drafting. The local Qwen model was the evaluated system, with its outputs and scoring recorded separately from the drafting assistance. The authors are responsible for verifying the scientific content and the final submitted version. AI tools are not authors.

### Author contributions

Lingsen You, Yujun Guo, Wentong Wang, Zisu Peng, and Xinyu Zhong contributed equally to this work. Li Shen and Junbo Ge are joint corresponding authors. Lingsen You: Conceptualization, Project administration, Writing - original draft. Yujun Guo: Methodology, Software. Wentong Wang: Data curation, Validation, Formal analysis. Li Shen: Supervision, Writing - review and editing. Junbo Ge: Supervision, Writing - review and editing. Data and analysis roles concern public research material, software evaluation and synthetic demonstrations; no new patient cohort or expert annotation panel is claimed.

## References

1. You L, Shen L, Ge J. The foundation and development of China’s National Basic Science Center for panvascular interventional complex systems: pioneering device-vessel suitcordance research. European Heart Journal. 2025;46:3400–3403. 10.1093/eurheartj/ehaf418

2. Garcia-Garcia HM, et al. Standardized End Point Definitions for Coronary Intervention Trials: The Academic Research Consortium-2 Consensus Document. European Heart Journal. 2018;39:2192–2207. 10.1093/eurheartj/ehy223

3. Patel MR, et al. Evaluation and Treatment of Patients With Lower Extremity Peripheral Artery Disease: Consensus Definitions From Peripheral Academic Research Consortium (PARC). Journal of the American College of Cardiology. 2015;65:931–941. 10.1016/j.jacc.2014.12.036

4. Nedeltchev K, et al. Standardized definitions and clinical endpoints in carotid artery and supra-aortic trunk revascularization trials. Catheterization and Cardiovascular Interventions. 2010;76:333–344. 10.1002/ccd.22560

5. Oderich GS, et al. Reporting standards for endovascular aortic repair of aneurysms involving the renal-mesenteric arteries. Journal of Vascular Surgery. 2021;73:4S–52S. 10.1016/j.jvs.2020.06.011

6. Vedantham S, et al. Delphi Consensus on Reporting Standards in Clinical Studies for Endovascular Treatment of Acute Iliofemoral Venous Thrombosis and Chronic Iliofemoral Venous Obstruction. Circulation Cardiovascular Interventions. 2023;16:e012894. 10.1161/CIRCINTERVENTIONS.123.012894

7. You L, Chen Y, Zhang Z, Wang Y, Gu Z, Shen L, Ge J. The FLOW Framework: a panvascular-on-a-chip platform to model systemic disease and guide panvascular interventional device suitcordance. Science Bulletin. 2026;71(3):654–670. 10.1016/j.scib.2025.12.051

8. You L, Yao J, Qiu Y, Wang Y, Sun Y, Zhang R, Shen L, Ge J. From intravascular imaging to adaptive vascular care: intelligent photonics and digital twins in panvascular disease. Light: Science & Applications. 2026;15:335. 10.1038/s41377-026-02410-6

9. Xue S, Zhong J, Nan Z, et al. ScienceBuddy: Recursive-in-Recursive Self-Improvement for Interactive Scientific Agents. arXiv. 2026;2609.17523. https://arxiv.org/abs/2609.17523

10. National Center for Biotechnology Information. A General Introduction to the E-utilities. Entrez Programming Utilities Help. https://www.ncbi.nlm.nih.gov/books/NBK25497/ Accessed 19 September 2026.

11. Europe PMC. RESTful Web Service. https://europepmc.org/RestfulWebService Accessed 19 September 2026.

12. Virtanen P, et al. SciPy 1.0: fundamental algorithms for scientific computing in Python. Nature Methods. 2020;17:261–272. 10.1038/s41592-019-0686-2

13. OpenAI. Text generation. OpenAI API documentation. https://developers.openai.com/api/docs/guides/text Accessed 19 September 2026.

14. Collins GS, et al. TRIPOD+AI statement: updated guidance for reporting clinical prediction models that use regression or machine learning methods. BMJ. 2024;385:e078378. 10.1136/bmj-2023-078378

15. Moons KGM, et al. PROBAST+AI: an updated quality, risk of bias, and applicability assessment tool for prediction models using regression or artificial intelligence methods. BMJ. 2025;388:e082505. 10.1136/bmj-2024-082505

16. You L, Guo Y, Peng Z, Wang W, Shen L, Ge J. AI-assisted generation and screening of PLLA modifiers for bioresorbable vascular scaffolds (in Chinese). Chinese Science Bulletin. 2026;71(19):4653–4662. 10.1360/CSB-2026-0332

17. Wang Z, Cao L, Danek B, Jin Q, Lu Z, Sun J. Accelerating clinical evidence synthesis with large language models. npj Digital Medicine. 2025;8:509. 10.1038/s41746-025-01840-7

18. Wang Z, Cao L, Jin Q, et al. A foundation model for human-AI collaboration in medical literature mining. Nature Communications. 2025;16:8361. 10.1038/s41467-025-62058-5

19. Witte C, Schmidt DM, Cimiano P. Comparing generative and extractive approaches to information extraction from abstracts describing randomized clinical trials. Journal of Biomedical Semantics. 2024;15:3. 10.1186/s13326-024-00305-2

20. Zhang W, Qiao M, Zang C, Niederer S, Matthews PM, Bai W, Kainz B. Multi-Agent Reasoning for Cardiovascular Imaging Phenotype Analysis. arXiv. 2025;2507.03460v2. https://arxiv.org/abs/2507.03460v2

21. Jullien M, Valentino M, Frost H, O’Regan P, Landers D, Freitas A. SemEval-2023 Task 7: Multi-Evidence Natural Language Inference for Clinical Trial Data. Proceedings of the 17th International Workshop on Semantic Evaluation (SemEval-2023). Toronto, Canada: Association for Computational Linguistics; 2023:2216–2226. 10.18653/v1/2023.semeval-1.307

22. Qwen Team. Qwen3-4B-GGUF. Model card and model weights, Q8_0 quantization, revision bc640142c66e1fdd12af0bd68f40445458f3869b. Hugging Face. https://huggingface.co/Qwen/Qwen3-4B-GGUF/tree/bc640142c66e1fdd12af0bd68f40445458f3869b Accessed 19 September 2026.

23. National Library of Medicine. ClinicalTrials.gov API, version 2: About the API. https://clinicaltrials.gov/data-api/about-api Accessed 19 September 2026.

24. International Council for Harmonisation of Technical Requirements for Pharmaceuticals for Human Use. Addendum on Estimands and Sensitivity Analysis in Clinical Trials to the Guideline on Statistical Principles for Clinical Trials, E9(R1). Final version, adopted 20 November 2019. https://database.ich.org/sites/default/files/E9-R1_Step4_Guideline_2019_1203.pdf

