## Supplementary data and reproducibility code for "PanVasc Research for AI assisted evidence analysis in panvascular intervention": app.py

```
"""Local desktop research workspace. No web server and no automatic model calls."""
from __future__ import annotations

import csv
import json
import queue
import threading
import uuid
from datetime import datetime
from pathlib import Path
import tkinter as tk
from tkinter import filedialog, messagebox, ttk
from tkinter.scrolledtext import ScrolledText

import pandas as pd

from panvasc import core, llm


BASE = Path(__file__).resolve().parent
OUTPUT = BASE / "local_workspace"


def json_text(value):
    return json.dumps(value, ensure_ascii=False, indent=2, allow_nan=False)


class ResearchApp(tk.Tk):
    def __init__(self):
        super().__init__()
        self.title("泛血管介入科研助手 · 本地研究原型")
        self.geometry("1160x840")
        self.minsize(920, 710)
        self.configure(background="#eef3f7")
        self.messages = queue.Queue()
        self.busy = False
        self.tasks = {}
        self.records = []
        self.search_task_id = None
        self.search_provenance = ""
        self.dataframe = None
        self.data_path = None
        self.domain_keys = {v["label"]: k for k, v in core.DOMAINS.items()}
        self.domain = tk.StringVar(value=next(iter(self.domain_keys)))
        self.status = tk.StringVar(value="就绪。模板和统计计算在本机执行；文献检索连接公开数据库。")
        self.remote_enabled = tk.BooleanVar(value=False)
        self._style()
        self._build()
        self._load_tasks()
        self.after(100, self._poll)

    def _style(self):
        style = ttk.Style(self)
        if "clam" in style.theme_names():
            style.theme_use("clam")
        style.configure("TFrame", background="#eef3f7")
        style.configure("TLabel", background="#eef3f7", font=("Microsoft YaHei UI", 10))
        style.configure("Title.TLabel", font=("Microsoft YaHei UI", 20, "bold"), foreground="#163849")
        style.configure("Note.TLabel", foreground="#4e6473", font=("Microsoft YaHei UI", 9))
        style.configure("TButton", font=("Microsoft YaHei UI", 10), padding=(12, 7))
        style.configure("TNotebook.Tab", font=("Microsoft YaHei UI", 10), padding=(18, 9))
        style.configure("Treeview", font=("Microsoft YaHei UI", 9), rowheight=28)
        style.configure("Treeview.Heading", font=("Microsoft YaHei UI", 9, "bold"))
        self.option_add("*Font", ("Microsoft YaHei UI", 10))

    def _build(self):
        top = ttk.Frame(self, padding=(24, 18, 24, 10))
        top.pack(fill="x")
        ttk.Label(top, text="泛血管介入科研助手", style="Title.TLabel").pack(anchor="w")
        ttk.Label(top, text="文献证据 · 研究设计 · 本地数据分析 · 研究者审阅", style="Note.TLabel").pack(anchor="w", pady=(3, 10))
        line = ttk.Frame(top)
        line.pack(fill="x")
        ttk.Label(line, text="研究领域").pack(side="left")
        ttk.Combobox(line, textvariable=self.domain, values=list(self.domain_keys), state="readonly", width=20).pack(side="left", padx=10)
        mode = "可选模型接口已配置；远程生成默认关闭" if llm.configured() else "模板模式 · 模型接口未配置"
        ttk.Label(line, text=mode, foreground="#176b69").pack(side="left", padx=10)
        ttk.Label(top, text="研究原型，输出需研究者复核。反馈仅保存审阅记录，当前不执行模型训练或自动改进。", style="Note.TLabel").pack(anchor="w", pady=(10, 0))
        self.tabs = ttk.Notebook(self)
        self.tabs.pack(fill="both", expand=True, padx=20, pady=8)
        self.literature_tab = ttk.Frame(self.tabs, padding=16)
        self.protocol_tab = ttk.Frame(self.tabs, padding=16)
        self.data_tab = ttk.Frame(self.tabs, padding=16)
        self.history_tab = ttk.Frame(self.tabs, padding=16)
        for frame, title in [(self.literature_tab, "文献检索"), (self.protocol_tab, "PICO 研究草案"), (self.data_tab, "本地 CSV 分析"), (self.history_tab, "任务、反馈与导出")]:
            self.tabs.add(frame, text=title)
        self._build_literature()
        self._build_protocol()
        self._build_data()
        self._build_history()
        bottom = ttk.Frame(self, padding=(22, 0, 22, 12))
        bottom.pack(fill="x")
        self.progress = ttk.Progressbar(bottom, mode="indeterminate", length=130)
        self.progress.pack(side="right", padx=(12, 0))
        ttk.Label(bottom, textvariable=self.status, style="Note.TLabel", wraplength=870).pack(side="left")

    def _text(self, parent, height=10):
        widget = ScrolledText(parent, height=height, wrap="word", font=("Microsoft YaHei UI", 10), relief="flat", padx=12, pady=10, background="#ffffff", foreground="#203844")
        widget.configure(state="disabled")
        return widget

    @staticmethod
    def _put(widget, text):
        widget.configure(state="normal")
        widget.delete("1.0", "end")
        widget.insert("1.0", text)
        widget.configure(state="disabled")

    def _build_literature(self):
        form = ttk.Frame(self.literature_tab)
        form.pack(fill="x")
        ttk.Label(form, text="英文关键词").grid(row=0, column=0, sticky="w")
        self.keywords = ttk.Entry(form)
        self.keywords.grid(row=0, column=1, columnspan=6, sticky="ew", padx=8)
        form.columnconfigure(1, weight=1)
        ttk.Button(form, text="检索文献", command=self._search).grid(row=0, column=7, padx=(6, 0))
        ttk.Label(form, text="短语用分号分隔，按 AND 组合。先访问 PubMed，连接失败时尝试 Europe PMC；请勿输入个人信息。", style="Note.TLabel").grid(row=1, column=0, columnspan=8, sticky="w", pady=6)
        options = ttk.Frame(self.literature_tab)
        options.pack(fill="x", pady=(0, 8))
        self.start_year = tk.StringVar(value="2015")
        self.end_year = tk.StringVar(value=str(datetime.now().year))
        self.limit = tk.StringVar(value="10")
        for label, variable, width in [("起始年份", self.start_year, 7), ("结束年份", self.end_year, 7), ("返回条数（1–25）", self.limit, 5)]:
            ttk.Label(options, text=label).pack(side="left", padx=(0, 6))
            ttk.Entry(options, textvariable=variable, width=width).pack(side="left", padx=(0, 18))
        self.search_note = tk.StringVar(value="尚未检索。结果展示来源元数据与摘要原文，不自动判断研究质量或结论。")
        ttk.Label(self.literature_tab, textvariable=self.search_note, style="Note.TLabel", wraplength=1000).pack(anchor="w")
        self.paper_list = ttk.Treeview(self.literature_tab, columns=("pmid", "year", "title"), show="headings", height=5, selectmode="browse")
        for key, title, width in [("pmid", "PMID", 95), ("year", "年份", 90), ("title", "原文标题", 720)]:
            self.paper_list.heading(key, text=title)
            self.paper_list.column(key, width=width, stretch=key == "title")
        self.paper_list.pack(fill="x", pady=8)
        self.paper_list.bind("<<TreeviewSelect>>", self._show_paper)
        self.paper_text = self._text(self.literature_tab, 7)
        self.paper_text.pack(fill="both", expand=True)
        remote = ttk.Frame(self.literature_tab)
        remote.pack(fill="x", pady=(12, 0))
        ttk.Label(remote, text="可选模型草案：公开课题问题").pack(anchor="w")
        self.model_question = ttk.Entry(remote)
        self.model_question.pack(fill="x", pady=5)
        remote_actions = ttk.Frame(remote)
        remote_actions.pack(fill="x")
        ttk.Checkbutton(remote_actions, text="我明确启用远程生成：仅发送上方公开问题及本次公开文献", variable=self.remote_enabled, command=self._refresh_model_button).pack(side="left")
        self.model_button = ttk.Button(remote_actions, text="生成研究讨论草案", command=self._generate_model, state="disabled")
        self.model_button.pack(side="right")
        ttk.Label(remote, text="需预先配置接口。CSV、数据分析和反馈不进入模型请求；请勿在问题框粘贴患者资料或未授权内容。", style="Note.TLabel").pack(anchor="w", pady=(4, 0))

    def _build_protocol(self):
        ttk.Label(self.protocol_tab, text="填写 PICO 与随访，生成固定结构草案。此操作不调用模型，也不检索或补造证据。", style="Note.TLabel").pack(anchor="w", pady=(0, 12))
        form = ttk.Frame(self.protocol_tab)
        form.pack(fill="x")
        form.columnconfigure(1, weight=1)
        self.pico = {}
        fields = [("population", "P · 研究人群"), ("intervention", "I · 干预或暴露"), ("comparator", "C · 对照"), ("outcome", "O · 主要结局"), ("follow_up", "随访时间 / 窗口")]
        for row, (key, label) in enumerate(fields):
            ttk.Label(form, text=label).grid(row=row, column=0, sticky="w", padx=(0, 14), pady=6)
            entry = ttk.Entry(form)
            entry.grid(row=row, column=1, sticky="ew", pady=6)
            self.pico[key] = entry
        ttk.Button(self.protocol_tab, text="生成并保存 PICO 模板", command=self._protocol).pack(anchor="w", pady=12)
        self.protocol_text = self._text(self.protocol_tab, 12)
        self.protocol_text.pack(fill="both", expand=True)

    def _build_data(self):
        ttk.Label(self.data_tab, text="CSV 在本机读取和计算；不会发送给模型。仅支持去标识研究表格，最多 100,000 行、300 列、50 MB。", style="Note.TLabel", wraplength=1000).pack(anchor="w", pady=(0, 10))
        files = ttk.Frame(self.data_tab)
        files.pack(fill="x")
        ttk.Button(files, text="选择本地 CSV", command=self._select_csv).pack(side="left")
        ttk.Label(files, text="编码").pack(side="left", padx=(12, 5))
        self.encoding = tk.StringVar(value="utf-8-sig")
        ttk.Combobox(files, textvariable=self.encoding, values=["utf-8-sig", "gb18030"], state="readonly", width=11).pack(side="left")
        self.file_note = tk.StringVar(value="尚未载入数据")
        ttk.Label(self.data_tab, textvariable=self.file_note, style="Note.TLabel", wraplength=1000).pack(anchor="w", pady=10)
        controls = ttk.Frame(self.data_tab)
        controls.pack(fill="x")
        self.columns = {}
        for index, (key, label) in enumerate([("id", "分析单位 ID"), ("group", "分组列"), ("outcome", "二元结局列（0/1）"), ("time", "随访时间列（可选）")]):
            ttk.Label(controls, text=label).grid(row=0, column=index, sticky="w", padx=(0, 15))
            box = ttk.Combobox(controls, state="readonly", width=21)
            box.grid(row=1, column=index, sticky="ew", padx=(0, 15), pady=(5, 10))
            controls.columnconfigure(index, weight=1)
            self.columns[key] = box
        actions = ttk.Frame(self.data_tab)
        actions.pack(fill="x", pady=(0, 10))
        ttk.Button(actions, text="数据质量 + 描述统计", command=self._audit).pack(side="left")
        ttk.Button(actions, text="两组二元结局比较", command=self._compare).pack(side="left", padx=10)
        ttk.Label(self.data_tab, text="二元比较：完整记录、未经调整的事件比例、Wilson 区间及双侧 Fisher 检验；不是因果估计，不处理删失。", style="Note.TLabel", wraplength=1000).pack(anchor="w", pady=(0, 8))
        self.data_text = self._text(self.data_tab, 15)
        self.data_text.pack(fill="both", expand=True)

    def _build_history(self):
        ttk.Label(self.history_tab, text="选择一个真实完成的任务，再审阅、保存反馈或导出。每个任务保留独立 ID；所有记录存于 local_workspace。", style="Note.TLabel", wraplength=1000).pack(anchor="w", pady=(0, 10))
        self.task_list = ttk.Treeview(self.history_tab, columns=("kind", "time", "task_id"), show="headings", height=5, selectmode="browse")
        for key, title, width in [("kind", "任务类型", 170), ("time", "完成时间（UTC）", 235), ("task_id", "真实任务 ID", 480)]:
            self.task_list.heading(key, text=title)
            self.task_list.column(key, width=width, stretch=key == "task_id")
        self.task_list.pack(fill="x")
        self.task_list.bind("<<TreeviewSelect>>", self._show_task)
        self.task_text = self._text(self.history_tab, 7)
        self.task_text.pack(fill="both", expand=True, pady=10)
        feedback = ttk.Frame(self.history_tab)
        feedback.pack(fill="x")
        ttk.Label(feedback, text="审阅者").pack(side="left")
        self.reviewer = ttk.Entry(feedback, width=20)
        self.reviewer.pack(side="left", padx=(7, 15))
        ttk.Label(feedback, text="审阅决定").pack(side="left")
        self.decision = ttk.Combobox(feedback, values=["accept · 接受", "revise · 修改", "reject · 拒绝"], state="readonly", width=16)
        self.decision.set("revise · 修改")
        self.decision.pack(side="left", padx=7)
        ttk.Button(feedback, text="导出选中任务 JSON", command=self._export).pack(side="right")
        ttk.Label(self.history_tab, text="修改内容或理由（修改 / 拒绝时必填）").pack(anchor="w", pady=(8, 4))
        self.correction = ScrolledText(self.history_tab, height=3, wrap="word")
        self.correction.pack(fill="x")
        ttk.Button(self.history_tab, text="保存选中任务的反馈", command=self._feedback).pack(anchor="w", pady=(8, 0))

    def _domain(self):
        return self.domain_keys[self.domain.get()]

    def _run(self, description, work, done):
        if self.busy:
            messagebox.showinfo("任务进行中", "请等当前操作完成；窗口仍可切换页面和查看已有记录。", parent=self)
            return
        self.busy = True
        self._refresh_model_button()
        self.status.set(description)
        self.progress.start(12)

        def worker():
            try:
                self.messages.put((True, work(), done))
            except Exception as exc:
                self.messages.put((False, f"{type(exc).__name__}: {exc}", None))
        threading.Thread(target=worker, daemon=True).start()

    def _poll(self):
        try:
            success, value, callback = self.messages.get_nowait()
        except queue.Empty:
            pass
        else:
            self.busy = False
            self.progress.stop()
            try:
                if not success:
                    raise RuntimeError(value)
                callback(value)
            except Exception as exc:
                self._error(exc)
            self._refresh_model_button()
        self.after(100, self._poll)

    def _error(self, exc):
        self.status.set("操作失败，未生成替代结果。请检查输入或网络后重试。")
        messagebox.showerror("操作未完成", str(exc), parent=self)

    def _save_task(self, kind, result, *, task_id=None, source_ids=None, inputs=None):
        task_id = task_id or str(uuid.uuid4())
        record = {"schema_version": "1.0", "task_id": task_id, "created_utc": core.utc_now(), "kind": kind,
                  "inputs": inputs or {}, "source_ids": source_ids or [], "result": result}
        folder = OUTPUT / "tasks"
        folder.mkdir(parents=True, exist_ok=True)
        # UUID is internally created, never a user-provided path.
        uuid.UUID(task_id)
        path = folder / f"{task_id}.json"
        serialized = json_text(record)
        with path.open("x", encoding="utf-8") as stream:
            stream.write(serialized)
        self.tasks[task_id] = record
        self._insert_task(record)
        self.task_list.selection_set(task_id)
        self.task_list.see(task_id)
        self._show_task()
        self.status.set(f"已完成并保存：{kind} · {task_id}")
        return task_id

    def _insert_task(self, record):
        self.task_list.insert("", 0, iid=record["task_id"], values=(record["kind"], record["created_utc"], record["task_id"]))

    def _load_tasks(self):
        skipped = 0
        for path in sorted((OUTPUT / "tasks").glob("*.json")):
            try:
                record = json.loads(path.read_text(encoding="utf-8"))
                task_id = record["task_id"]
                uuid.UUID(task_id)
                if path.stem != task_id or task_id in self.tasks:
                    raise ValueError("Task identity mismatch")
                self._insert_task(record)
                self.tasks[task_id] = record
            except (ValueError, OSError, KeyError, TypeError, tk.TclError):
                skipped += 1
        if skipped:
            self.status.set(f"已载入 {len(self.tasks)} 个任务；有 {skipped} 个记录无法读取，原文件已保留。")

    def _search(self):
        if self.busy:
            messagebox.showinfo("任务进行中", "请等当前操作完成后再开始新检索。", parent=self)
            return
        try:
            domain, keywords = self._domain(), self.keywords.get().strip()
            start, end, limit = int(self.start_year.get()), int(self.end_year.get()), int(self.limit.get())
            core.build_query(domain, keywords, start, end)
            if not 1 <= limit <= 25:
                raise ValueError("返回条数应为 1–25。")
        except Exception as exc:
            self._error(exc)
            return

        # Do not let a failed new search silently reuse the previous search's evidence.
        self.records = []
        self.search_task_id = None
        self.search_provenance = ""
        self.paper_list.delete(*self.paper_list.get_children())
        self.search_note.set("本次检索尚无成功结果；当前没有可发送给模型的文献。此前结果仍保留在历史任务中。")
        self._put(self.paper_text, "等待本次检索完成。若检索失败，模型生成将保持禁用。")
        self._refresh_model_button()

        def done(result):
            result.setdefault("fallback_used", False)
            source = result.get("api_source", "来源未标记")
            task_id = self._save_task("文献检索 · " + source, result, task_id=result["run_id"], source_ids=[r["pmid"] for r in result["records"]])
            self.records, self.search_task_id = result["records"], task_id
            self.search_provenance = f"实际接口：{source}\n备用来源 fallback_used：{result['fallback_used']}\n实际检索式：{result['query']}"
            if result.get("primary_failure"):
                failure = result["primary_failure"]
                self.search_provenance += f"\n主来源失败 primary_failure：{failure.get('service')} / {failure.get('exception')}\n{failure.get('detail', '')[:240]}\n完整错误保存在任务 JSON 中。"
            if result.get("query_semantics"):
                self.search_provenance += f"\n检索范围说明：{result['query_semantics']}"
            self.paper_list.delete(*self.paper_list.get_children())
            for record in self.records:
                self.paper_list.insert("", "end", iid=record["pmid"], values=(record["pmid"], record["year"], record["title"]))
            fallback_note = " · 主来源失败，已使用备用检索；完整原因见下方" if result["fallback_used"] else " · 未使用备用检索"
            self.search_note.set(f"{source}{fallback_note}\n{core.DOMAINS[result['domain']]['label']} · 命中 {result['count']} 条，取得 {len(self.records)} 条。")
            self._put(self.paper_text, self.search_provenance + "\n\n" + ("没有返回匹配记录。可调整关键词或年份。" if not self.records else "请选择文献查看原始摘要。"))
            if self.records:
                self.paper_list.selection_set(self.records[0]["pmid"])
                self._show_paper()
            self._refresh_model_button()
        self._run("正在检索文献；先访问 PubMed，失败时尝试 Europe PMC 并明确记录来源……", lambda: core.search_literature(domain, keywords, limit, start, end, OUTPUT / "literature"), done)

    def _show_paper(self, _event=None):
        selection = self.paper_list.selection()
        if not selection:
            return
        record = next((r for r in self.records if r["pmid"] == selection[0]), None)
        if not record:
            return
        warning = "\n撤稿相关标记：有，需核对来源通知。" if record["retraction_flag"] else ""
        if record.get("notice_coverage"):
            warning += "\n通知检索范围：" + record["notice_coverage"]
        text = (f"{self.search_provenance}\n\n{record['title']}\n\nPMID: {record['pmid']}  |  年份: {record['year']}\n"
                f"期刊: {record['journal']}\nDOI: {record.get('doi') or '来源未提供'}\n{record['url']}{warning}\n\n"
                f"摘要原文\n{record.get('abstract') or '来源未提供摘要。'}")
        self._put(self.paper_text, text)

    def _refresh_model_button(self):
        enabled = not self.busy and llm.configured() and self.remote_enabled.get() and bool(self.records) and bool(self.search_task_id)
        self.model_button.configure(state="normal" if enabled else "disabled")

    def _generate_model(self):
        question = self.model_question.get().strip()
        if not llm.configured() or not self.remote_enabled.get() or not self.records or not self.search_task_id:
            self._error(ValueError("需配置模型接口、明确勾选远程生成，并先取得真实公开文献。"))
            return
        if not question or len(question) > 6000:
            self._error(ValueError("请填写 1–6000 字的公开课题问题。"))
            return
        records = json.loads(json_text(self.records))
        parent_id = self.search_task_id

        def done(result):
            self._save_task("模型研究草案 · 未验证", result, source_ids=[r["pmid"] for r in records], inputs={"public_question": question, "literature_task_id": parent_id, "remote_explicitly_enabled": True, "csv_transmitted": False})
            self.tabs.select(self.history_tab)
        self._run("正在调用已配置模型；仅发送公开课题问题与本次文献，草案需复核……", lambda: llm.generate(question, records, allow_remote=True), done)

    def _protocol(self):
        try:
            domain = self._domain()
            fields = {key: entry.get().strip() for key, entry in self.pico.items()}
            result = core.make_protocol(domain, **fields)
            self._save_task("PICO 固定模板", result)
            self._put(self.protocol_text, "固定模板草案 · 未执行模型推理\n\n" + json_text(result))
        except Exception as exc:
            self._error(exc)

    def _select_csv(self):
        if self.busy:
            messagebox.showinfo("任务进行中", "请等待当前操作完成后选择 CSV。", parent=self)
            return
        selected = filedialog.askopenfilename(parent=self, title="选择本地研究 CSV", filetypes=[("CSV 文件", "*.csv"), ("所有文件", "*.*")])
        if not selected:
            return
        path, encoding = Path(selected), self.encoding.get()

        def load():
            if path.stat().st_size > 50 * 1024 * 1024:
                raise ValueError("文件超过原型 50 MB 限制。")
            with path.open(encoding=encoding, newline="") as stream:
                header = next(csv.reader(stream), [])
            if not header or len(header) != len(set(header)) or any(not h.strip() for h in header):
                raise ValueError("CSV 列名须非空且唯一；请先修正列名。")
            if len(header) > 300:
                raise ValueError("CSV 超过 300 列限制。")
            frame = pd.read_csv(path, encoding=encoding, nrows=100001)
            if frame.empty or len(frame) > 100000:
                raise ValueError("CSV 应包含 1–100,000 行数据。")
            return frame

        def done(frame):
            self.dataframe, self.data_path = frame, path
            values = [str(c) for c in frame.columns]
            defaults = {"id": "patient_id", "group": "group", "outcome": "outcome", "time": "follow_up"}
            for key, box in self.columns.items():
                box.configure(values=[""] + values)
                box.set(defaults[key] if defaults[key] in values else "")
            self.file_note.set(f"{path.name} · {len(frame):,} 行 × {len(frame.columns)} 列 · 本地内存")
            self._put(self.data_text, "文件已载入。请选择列，然后运行数据质量检查或二元比较。\n\n列名：\n" + "\n".join(values))
            self.status.set("CSV 已在本机载入，未向任何模型服务发送。")
        self._run("正在本机读取 CSV……", load, done)

    def _data_inputs(self):
        if self.dataframe is None:
            raise ValueError("请先选择本地 CSV。")
        return self.dataframe, {key: box.get().strip() for key, box in self.columns.items()}, {"csv_name": self.data_path.name, "csv_path_local": str(self.data_path), "remote_transmission": False}

    def _audit(self):
        try:
            frame, columns, inputs = self._data_inputs()
        except Exception as exc:
            self._error(exc)
            return

        def work():
            audit = core.audit_dataframe(frame, id_column=columns["id"] or "patient_id", outcome=columns["outcome"] or None, time_column=columns["time"] or None)
            try:
                description = core.describe_dataframe(frame)
                description_error = None
            except ValueError as exc:
                description, description_error = None, str(exc)
            return {"quality_audit": audit, "descriptive_statistics": description, "descriptive_error": description_error}

        def done(result):
            self._save_task("本地数据质量与描述", result, inputs={**inputs, "selected_columns": columns})
            self._put(self.data_text, "本地计算结果\n注意：自动检查不证明数据已去标识、设计有效或完全不存在泄漏。\n\n" + json_text(result))
        self._run("正在本机检查数据质量并计算描述统计……", work, done)

    def _compare(self):
        try:
            frame, columns, inputs = self._data_inputs()
            if not columns["group"] or not columns["outcome"]:
                raise ValueError("请选择分组列和二元结局列。")
        except Exception as exc:
            self._error(exc)
            return

        def done(result):
            self._save_task("本地两组二元比较", result, inputs={**inputs, "selected_columns": columns})
            self._put(self.data_text, "未经调整的两组二元比较\n组别顺序及缺失记录剔除数量见结果。\n\n" + json_text(result))
        self._run("正在本机计算事件比例、区间与 Fisher 精确检验……", lambda: core.compare_binary(frame, columns["group"], columns["outcome"]), done)

    def _selected_task(self):
        selection = self.task_list.selection()
        if not selection or selection[0] not in self.tasks:
            raise ValueError("请先选择一个已完成并保存的任务。")
        return self.tasks[selection[0]]

    def _show_task(self, _event=None):
        try:
            self._put(self.task_text, json_text(self._selected_task()))
        except ValueError:
            pass

    def _feedback(self):
        try:
            task = self._selected_task()
            feedback = core.save_feedback(OUTPUT / "feedback.jsonl", task["task_id"], self.decision.get().split(" ", 1)[0], self.correction.get("1.0", "end").strip(), self.reviewer.get().strip(), task["source_ids"])
            self.status.set(f"反馈已保存并绑定任务 {task['task_id']}；未触发训练。")
            messagebox.showinfo("反馈已保存", f"任务 ID：{task['task_id']}\n反馈 ID：{feedback['feedback_id']}\n保存位置：local_workspace/feedback.jsonl\n未更新模型权重。", parent=self)
        except Exception as exc:
            self._error(exc)

    def _export(self):
        try:
            task = self._selected_task()
            selected = filedialog.asksaveasfilename(parent=self, title="导出选中任务", defaultextension=".json", initialfile=f"panvasc-{task['task_id']}.json", filetypes=[("JSON", "*.json")])
            if selected:
                Path(selected).write_text(json_text(task), encoding="utf-8")
                self.status.set(f"已导出：{selected}")
        except Exception as exc:
            self._error(exc)


if __name__ == "__main__":
    ResearchApp().mainloop()
```
