## Supplementary data and reproducibility code for "PanVasc Research for AI assisted evidence analysis in panvascular intervention": MODEL_CARD.md

# PanVasc Research system and evaluated model

## Identity and intended use

The original desktop prototype supports exploratory literature retrieval, structured PICO templates, local CSV checks and basic summaries, and task-linked reviewer feedback. The revised study adds a separate command-line local-model pilot. Its evaluated model is the fixed official Qwen3-4B Q8_0 checkpoint, not newly trained vascular weights. The desktop's optional remote API adapter remains untested and is distinct from this runner.

## Observed evidence

The local pilot made 700 test calls, comparing generic and checklist prompts on the same evidence. Registry exact agreement was 51/200 (25.5%) and 77/200 (38.5%) across 200 controlled tasks; another 50 clean tasks per arm were controls. NLI label accuracy was 48/100 (48.0%) and 50/100 (50.0%) on a 100-statement external breast-cancer sample. Contracts inspect source fields or index validity and do not validate clinical reasoning. No patient-level prediction, researcher utility, model training or recursive optimization was evaluated.

Earlier software evidence consists of 28/28 development checks, final five-domain retrieval returning 15 records, and a synthetic numerical example. Earlier transport failures remain documented. These are separate from model accuracy.

## Material and limitations

Registry strata are query-derived and the data are convenience-sampled. Deletion conditions are experimentally constructed. NLI labels are original public expert annotations, not new vascular labels. Pretraining contamination is unknown; related trial families and reused distractors limit independence. One model, quantization, generation seed and small test sample cannot establish panvascular superiority.

## Data movement and reproduction

Public API requests acquire literature and registry material. The reported inference uses a loopback server at 127.0.0.1:18123 and sends no participant records to external inference services. Optional desktop remote generation remains separately configured and explicitly activated by its operator. No automatic feedback-to-weight update occurs.

The protocol, code, hashes, prompt settings, all numerical scores and exact model/runtime versions are in the pilot supplement. Third-party source text, full model responses, weights and credentials are not redistributed there. Retrieve NLI4CT at its pinned official revision; mutable live registry records may not reconstruct the historical snapshot. Start the recorded local model server before executing the frozen pilot runner. A new reproduction must keep its own freeze and outputs rather than mixing runs.
