## Supplementary data and reproducibility code for "PanVasc Research for AI assisted evidence analysis in panvascular intervention": core.py

```
"""Deterministic, inspectable components for a panvascular research assistant.

No LLM output is needed for these operations. Biomedical text is data, never code.
"""
from __future__ import annotations
import hashlib
import json
import math
import numbers
import re
import time
import uuid
import xml.etree.ElementTree as ET
from datetime import datetime, timezone
from pathlib import Path
from typing import Any

import numpy as np
import pandas as pd
import requests
from scipy.stats import fisher_exact

DOMAINS = {
    "coronary": {"label": "冠状动脉", "term": '("Percutaneous Coronary Intervention"[MeSH Terms] OR "coronary intervention"[Title/Abstract])', "endpoint": "Target lesion failure at a prespecified follow-up; define each component and adjudication rule."},
    "peripheral": {"label": "外周动脉", "term": '("Peripheral Arterial Disease"[MeSH Terms] AND (endovascular[Title/Abstract] OR angioplasty[Title/Abstract]))', "endpoint": "Major adverse limb events or target lesion revascularization; do not treat patency as an interchangeable endpoint."},
    "carotid": {"label": "颈动脉与脑血管", "term": '("Carotid Stenosis"[MeSH Terms] AND (stenting[Title/Abstract] OR endovascular[Title/Abstract]))', "endpoint": "Prespecified stroke/death endpoint with laterality, severity, and time window recorded separately."},
    "aortic": {"label": "主动脉", "term": '("Aortic Aneurysm"[MeSH Terms] AND (endovascular[Title/Abstract] OR EVAR[Title/Abstract]))', "endpoint": "Aortic reintervention or aneurysm-related mortality; specify anatomical segment, rupture status, and device generation."},
    "venous": {"label": "静脉", "term": '("Venous Thrombosis"[MeSH Terms] AND (thrombectomy[Title/Abstract] OR endovascular[Title/Abstract]))', "endpoint": "Recurrent venous thromboembolism or post-thrombotic syndrome with a named definition and assessment time."},
}

def utc_now():
    return datetime.now(timezone.utc).isoformat()

def digest(value: Any):
    return hashlib.sha256(json.dumps(value, ensure_ascii=False, sort_keys=True, allow_nan=False).encode("utf-8")).hexdigest()

def normalize_identifier(value):
    """Normalize scalar identifiers; preserve meaningful leading zeroes in strings."""
    if pd.isna(value) or (isinstance(value,str) and not value.strip()):
        raise ValueError("Patient identifiers must be nonmissing and nonblank")
    if isinstance(value,numbers.Real):
        if not math.isfinite(value):
            raise ValueError("Patient identifiers must be finite")
        if float(value).is_integer():
            return str(int(value))
    return str(value).strip()

def build_query(domain: str, keywords: str = "", start_year: int = 2015, end_year: int = 2026):
    if domain not in DOMAINS:
        raise ValueError("Unknown vascular domain")
    if not 1900 <= int(start_year) <= int(end_year) <= datetime.now().year:
        raise ValueError("Invalid publication-year interval")
    # Inputs are literal title/abstract phrases. Advanced PubMed syntax is intentionally not accepted here.
    phrases = [p.strip() for p in re.split(r"[;；\n]", str(keywords)) if p.strip()]
    if len(phrases) > 8 or any(len(p) > 120 for p in phrases):
        raise ValueError("Use at most eight short phrases separated by semicolons")
    clean = [re.sub(r'[^\w\s\-]', ' ', p, flags=re.UNICODE).strip() for p in phrases]
    if any(not p for p in clean):
        raise ValueError("Empty keyword after normalization")
    extras = "".join(f' AND "{p}"[Title/Abstract]' for p in clean)
    return f'{DOMAINS[domain]["term"]}{extras} AND ("{int(start_year)}"[Date - Publication] : "{int(end_year)}"[Date - Publication])'

def _text(element):
    return "".join(element.itertext()).strip() if element is not None else ""

def parse_pubmed(xml: str):
    root = ET.fromstring(xml)
    records, seen = [], set()
    for item in root.findall(".//PubmedArticle"):
        pmid = _text(item.find(".//MedlineCitation/PMID"))
        if not pmid.isdigit() or pmid in seen:
            continue
        seen.add(pmid)
        title = _text(item.find(".//ArticleTitle"))
        doi = next((_text(e) for e in item.findall(".//ArticleId") if e.get("IdType") == "doi"), None)
        abstracts = item.findall(".//Abstract/AbstractText")
        abstract = "\n".join(((e.get("Label") + ": ") if e.get("Label") else "") + _text(e) for e in abstracts)
        types = [_text(e) for e in item.findall(".//PublicationType")]
        notices = [{"type": e.get("RefType"), "pmid": _text(e.find("PMID"))} for e in item.findall(".//CommentsCorrections")]
        flagged = any("retract" in t.lower() for t in types) or any("retract" in str(n["type"]).lower() for n in notices)
        authors = [" ".join(filter(None, [_text(a.find("ForeName")), _text(a.find("LastName"))])) or _text(a.find("CollectiveName")) for a in item.findall(".//AuthorList/Author")]
        year = _text(item.find(".//JournalIssue/PubDate/Year")) or _text(item.find(".//JournalIssue/PubDate/MedlineDate"))
        records.append({"pmid": pmid, "title": title, "doi": doi, "authors": authors,
                        "journal": _text(item.find(".//Journal/Title")), "year": year,
                        "abstract": abstract or None, "publication_types": types,
                        "retraction_flag": flagged, "notices": notices,
                        "url": f"https://pubmed.ncbi.nlm.nih.gov/{pmid}/", "source": "PubMed"})
    return records

def search_pubmed(domain: str, keywords: str = "", limit: int = 5, start_year: int = 2015, end_year: int = 2026, output_dir: str | Path | None = None):
    query = build_query(domain, keywords, start_year, end_year)
    if not 1 <= int(limit) <= 25:
        raise ValueError("limit must be 1-25")
    started, timestamp = time.perf_counter(), utc_now()
    base = "https://eutils.ncbi.nlm.nih.gov/entrez/eutils/"
    headers = {"User-Agent": "PanVascResearch/0.1 research-prototype"}
    def fetch(name, params):
        for attempt in range(3):
            response = requests.get(base + name, params={**params, "tool": "panvasc_research"}, headers=headers, timeout=35)
            if response.status_code not in (429, 500, 502, 503):
                response.raise_for_status()
                return response
            time.sleep(1 + attempt)
        response.raise_for_status()
    raw_search = fetch("esearch.fcgi", {"db": "pubmed", "term": query, "retmode": "json", "retmax": int(limit), "sort": "relevance"})
    search = raw_search.json()["esearchresult"]
    ids = search["idlist"]
    time.sleep(0.4)  # remain below NCBI's unauthenticated three-requests/second ceiling
    raw_xml = "<PubmedArticleSet/>" if not ids else fetch("efetch.fcgi", {"db": "pubmed", "id": ",".join(ids), "retmode": "xml"}).text
    result = {"run_id": str(uuid.uuid4()), "timestamp_utc": timestamp, "domain": domain, "query": query, "api_source": "NCBI E-utilities",
              "query_translation": search.get("querytranslation"), "count": int(search["count"]),
              "requested_pmids": ids, "records": parse_pubmed(raw_xml), "elapsed_seconds": round(time.perf_counter()-started, 3),
              "status": "success", "scope": "Metadata retrieval; relevance and claim entailment not adjudicated."}
    result["source_sha256"] = hashlib.sha256(raw_xml.encode("utf-8")).hexdigest()
    if output_dir:
        folder = Path(output_dir) / result["run_id"]
        folder.mkdir(parents=True, exist_ok=False)
        (folder / "esearch.json").write_text(raw_search.text, encoding="utf-8")
        (folder / "efetch.xml").write_text(raw_xml, encoding="utf-8")
        (folder / "result.json").write_text(json.dumps(result, ensure_ascii=False, indent=2), encoding="utf-8")
    return result

def make_protocol(domain: str, population: str, intervention: str, comparator: str, outcome: str, follow_up: str):
    if domain not in DOMAINS:
        raise ValueError("Unknown vascular domain")
    fields = dict(population=population, intervention=intervention, comparator=comparator, outcome=outcome, follow_up=follow_up)
    if any(not isinstance(v, str) or not v.strip() or len(v) > 3000 for v in fields.values()):
        raise ValueError("Every PICO field and follow-up must contain 1-3000 characters")
    return {"schema_version": "1.0", "mode": "deterministic_template", "domain": domain, "pico": fields,
            "research_question": f"Among {population}, how does {intervention}, compared with {comparator}, relate to {outcome} over {follow_up}?",
            "endpoint_design_note": DOMAINS[domain]["endpoint"],
            "design": "Select randomized trial, observational cohort, or evidence synthesis before analysis; this template does not choose a design from the question.",
            "required_decisions": ["Define index date, analysis unit, inclusion/exclusion, and target estimand.",
                "Specify primary endpoint, components, adjudication, and follow-up before accessing outcome associations.",
                "For observational comparisons, prespecify confounders and acknowledge residual confounding.",
                "For time-to-event outcomes, prespecify censoring and competing risks; crude risks are insufficient with unequal follow-up.",
                "Specify missingness, clustering, multiplicity, and sensitivity analyses.",
                "For prediction, split by patient and consider time/site validation; fit preprocessing on training data only.",
                "Obtain investigator review and any required research approvals before real-data use."],
            "status": "Draft for researcher review; no clinical recommendation or trained-model inference."}

def search_europepmc(domain: str, keywords: str = "", limit: int = 5, start_year: int = 2015, end_year: int = 2026, output_dir: str | Path | None = None):
    """Explicit alternative MEDLINE route. Query semantics differ from PubMed."""
    build_query(domain, keywords, start_year, end_year)  # common input validation
    if not 1 <= int(limit) <= 25:
        raise ValueError("limit must be 1-25")
    terms = {
        "coronary": '(TITLE_ABS:"coronary" AND (TITLE_ABS:"intervention" OR TITLE_ABS:"stent"))',
        "peripheral": '((TITLE_ABS:"peripheral arterial" OR TITLE_ABS:"femoropopliteal") AND (TITLE_ABS:"endovascular" OR TITLE_ABS:"angioplasty"))',
        "carotid": '(TITLE_ABS:"carotid" AND (TITLE_ABS:"stenting" OR TITLE_ABS:"endovascular"))',
        "aortic": '(TITLE_ABS:"aortic" AND (TITLE_ABS:"endovascular" OR TITLE_ABS:"EVAR"))',
        "venous": '(TITLE_ABS:"venous" AND (TITLE_ABS:"thrombectomy" OR TITLE_ABS:"endovascular"))',
    }
    extras = [re.sub(r'[^\w\s\-]', ' ', p).strip() for p in re.split(r"[;；\n]", keywords) if p.strip()]
    query = f'SRC:MED AND {terms[domain]} AND FIRST_PDATE:[{start_year}-01-01 TO {end_year}-12-31]' + ''.join(f' AND TITLE_ABS:"{p}"' for p in extras)
    started, timestamp = time.perf_counter(), utc_now()
    attempts = []
    for attempt in range(3):
        try:
            response = requests.get("https://www.ebi.ac.uk/europepmc/webservices/rest/search", params={"query":query,"format":"json","resultType":"core","pageSize":int(limit)}, timeout=20)
            attempts.append({"attempt":attempt+1,"http_status":response.status_code})
            response.raise_for_status()
            break
        except requests.RequestException as exc:
            if not attempts or attempts[-1].get("attempt") != attempt+1:
                attempts.append({"attempt":attempt+1,"exception":type(exc).__name__})
            status = getattr(getattr(exc,"response",None),"status_code",None)
            if attempt == 2 or (status is not None and status not in (429,500,502,503,504)):
                raise
            time.sleep(0.5*(attempt+1))
    payload = response.json()
    records, seen = [], set()
    for item in payload.get("resultList",{}).get("result",[]):
        pmid = str(item.get("pmid", ""))
        if not pmid.isdigit() or pmid in seen:
            continue
        seen.add(pmid)
        types = item.get("pubTypeList",{}).get("pubType",[])
        records.append({"pmid":pmid,"title":item.get("title", ""),"doi":item.get("doi"),
           "authors":[a.get("fullName","") for a in item.get("authorList",{}).get("author",[])],
           "journal":item.get("journalInfo",{}).get("journal",{}).get("title", ""),"year":item.get("pubYear"),
           "abstract":item.get("abstractText"),"publication_types":types,
           "retraction_flag":any("retract" in str(t).lower() for t in types), "notices":[],
           "notice_coverage":"Publication types only; linked notices not checked by this adapter.",
           "url":f"https://pubmed.ncbi.nlm.nih.gov/{pmid}/", "source":"Europe PMC MEDLINE"})
    result={"run_id":str(uuid.uuid4()),"timestamp_utc":timestamp,"domain":domain,"query":query,
        "api_source":"Europe PMC REST","query_semantics":"Europe PMC title/abstract; not equivalent to PubMed MeSH query",
        "count":int(payload.get("hitCount",0)),"records":records,"elapsed_seconds":round(time.perf_counter()-started,3),
        "source_sha256":hashlib.sha256(response.content).hexdigest(),"status":"success","transport_attempts":attempts,
        "scope":"Metadata retrieval only; relevance and claim entailment not adjudicated."}
    if output_dir:
        folder=Path(output_dir)/result["run_id"]
        folder.mkdir(parents=True,exist_ok=False)
        (folder/"europepmc.json").write_text(response.text,encoding="utf-8")
        (folder/"result.json").write_text(json.dumps(result,ensure_ascii=False,indent=2),encoding="utf-8")
    return result

def search_literature(domain: str, keywords: str = "", limit: int = 5, start_year: int = 2015, end_year: int = 2026, output_dir: str | Path | None = None):
    try:
        result=search_pubmed(domain,keywords,limit,start_year,end_year,output_dir)
        result["api_source"]="NCBI E-utilities"
        return result
    except (requests.RequestException, ET.ParseError, KeyError) as exc:
        result=search_europepmc(domain,keywords,limit,start_year,end_year,output_dir)
        result["primary_failure"]={"service":"NCBI E-utilities","exception":type(exc).__name__,"detail":str(exc)}
        result["fallback_used"]=True
        # Save the fallback decision as well as the raw response so its provenance is explicit.
        if output_dir:
            (Path(output_dir)/result["run_id"]/"result.json").write_text(json.dumps(result,ensure_ascii=False,indent=2),encoding="utf-8")
        return result

def audit_dataframe(df: pd.DataFrame, id_column: str = "patient_id", outcome: str | None = None, time_column: str | None = None, features: list[str] | None = None):
    if not isinstance(df, pd.DataFrame) or df.empty:
        raise ValueError("Dataset is empty")
    if len(df) > 100000 or len(df.columns) > 300:
        raise ValueError("Prototype limit is 100,000 rows and 300 columns")
    if not df.columns.is_unique:
        raise ValueError("Column names must be unique")
    issues = []
    def issue(code, detail, severity="review"):
        issues.append({"code": code, "detail": detail, "severity": severity})
    if id_column not in df:
        issue("missing_id", f"Missing analysis-unit identifier: {id_column}")
    else:
        normalized, missing_identifiers = [], 0
        for value in df[id_column]:
            try:
                normalized.append(normalize_identifier(value))
            except ValueError:
                missing_identifiers += 1
        duplicates = len(normalized)-len(set(normalized))
        if duplicates:
            issue("duplicate_id", f"{duplicates} repeated nonmissing identifiers; determine clustering or duplicate records.")
        if missing_identifiers:
            issue("missing_id_value", f"{missing_identifiers} records have a missing, blank, or invalid identifier.", "error")
    missing = {str(c): int(df[c].isna().sum()) for c in df.columns}
    for c, n in missing.items():
        if n:
            issue("missing_values", f"{c}: {n}/{len(df)} missing; no imputation performed.")
    for col in df.select_dtypes(include="number"):
        if np.isinf(df[col].to_numpy(dtype=float)).any():
            issue("nonfinite", f"{col} includes infinite values", "error")
    if outcome:
        if outcome not in df:
            issue("missing_outcome", f"Outcome column {outcome} is absent", "error")
        elif df[outcome].dropna().empty:
            issue("missing_outcome_values", "No observed outcome values are available", "error")
        elif not set(df[outcome].dropna().unique()).issubset({0, 1}):
            issue("nonbinary_outcome", "Expected binary outcome values 0 or 1", "error")
        if outcome in (features or []):
            issue("outcome_leakage", "The outcome was included in the proposed predictors", "error")
    if time_column:
        if time_column not in df:
            issue("missing_follow_up", f"Follow-up column {time_column} is absent", "error")
        else:
            t = pd.to_numeric(df[time_column], errors="coerce")
            if (t.dropna() < 0).any():
                issue("negative_follow_up", "Negative follow-up is invalid", "error")
            if t.isna().any():
                issue("invalid_follow_up", "Missing or nonnumeric follow-up requires review", "error")
    for c in features or []:
        if c not in df:
            issue("missing_feature", f"Feature {c} is absent", "error")
        if any(s in c.lower() for s in ("post_", "future_", "discharge", "followup_")):
            issue("temporal_leakage_review", f"Confirm {c} is available at the prediction index time")
    return {"rows": int(len(df)), "columns": int(len(df.columns)), "missing_counts": missing,
            "issues": issues, "analysis_ready": not any(i["severity"] == "error" for i in issues),
            "limitation": "Heuristic checks do not establish de-identification, unit consistency, causal validity, or absence of all leakage."}

def describe_dataframe(df: pd.DataFrame):
    audit_dataframe(df)
    result = {}
    for col in df:
        s = df[col]
        if pd.api.types.is_numeric_dtype(s):
            numeric = s.dropna().astype(float)
            if np.isinf(numeric).any():
                raise ValueError("Infinite values must be resolved before descriptive statistics")
            result[str(col)] = {"type": "numeric", "n": int(len(numeric)), "missing": int(s.isna().sum()),
                 "mean": float(numeric.mean()) if len(numeric) else None,
                 "sd": float(numeric.std(ddof=1)) if len(numeric)>1 else None,
                 "median": float(numeric.median()) if len(numeric) else None,
                 "q1": float(numeric.quantile(.25)) if len(numeric) else None,
                 "q3": float(numeric.quantile(.75)) if len(numeric) else None}
        else:
            result[str(col)] = {"type": "categorical", "n": int(s.notna().sum()), "missing": int(s.isna().sum()),
                                "levels": {str(k): int(v) for k,v in s.value_counts().head(30).items()}}
    return result

def compare_binary(df: pd.DataFrame, group: str, outcome: str):
    if group not in df or outcome not in df or group == outcome:
        raise ValueError("Specify distinct existing group and outcome columns")
    data = df[[group, outcome]].dropna()
    if data.empty or not set(data[outcome].unique()).issubset({0, 1}):
        raise ValueError("Outcome must be binary 0/1")
    levels = sorted(data[group].unique(), key=str)
    if len(levels) != 2:
        raise ValueError("Exactly two observed groups are required")
    table = []
    summaries = []
    for level in levels:
        values = data.loc[data[group] == level, outcome]
        n, events = int(len(values)), int(values.sum())
        table.append([events, n-events])
        # Wilson score confidence interval for each group's crude event proportion.
        p, z = events/n, 1.959963984540054
        denom = 1 + z*z/n
        center = (p + z*z/(2*n))/denom
        half = z*math.sqrt(p*(1-p)/n + z*z/(4*n*n))/denom
        summaries.append({"group": str(level), "n": n, "events": events, "risk": p,
                          "wilson_95_ci": [max(0.,center-half), min(1.,center+half)]})
    odds, pvalue = fisher_exact(table, alternative="two-sided")
    return {"method": "Unadjusted 2x2 comparison; Fisher exact test, two-sided", "groups": summaries,
            "table_events_nonevents": table, "excluded_missing": int(len(df)-len(data)),
            "risk_difference_first_minus_second": summaries[0]["risk"]-summaries[1]["risk"],
            "odds_ratio": float(odds) if math.isfinite(odds) else None, "fisher_p": float(pvalue),
            "limitations": "Descriptive only; no causal interpretation, multiplicity adjustment, clustering adjustment, or censoring model."}

def validate_split(train_ids, test_ids):
    train, test = list(train_ids), list(test_ids)
    if not train or not test or any(pd.isna(x) for x in train+test):
        raise ValueError("Nonempty splits with complete patient identifiers are required")
    overlap = sorted(set(map(normalize_identifier, train)) & set(map(normalize_identifier, test)))
    return {"valid": not overlap, "overlap_count": len(overlap), "overlapping_ids": overlap}

def validate_citations(claims: list[dict], records: list[dict]):
    available = {str(r["pmid"]): r for r in records}
    checks = []
    for claim in claims:
        cited = [str(p) for p in claim.get("pmids", [])]
        missing = [p for p in cited if p not in available]
        flagged = [p for p in cited if p in available and available[p].get("retraction_flag")]
        checks.append({"claim": claim.get("text", ""), "pmids": cited, "unknown_pmids": missing,
                       "flagged_pmids": flagged, "identity_valid": bool(cited) and not missing and not flagged,
                       "entailment": "not_evaluated"})
    return checks

def save_feedback(path: str | Path, task_id: str, decision: str, correction: str, reviewer: str, source_ids: list[str] | None = None):
    if decision not in {"accept", "revise", "reject"}:
        raise ValueError("Feedback decision must be accept, revise, or reject")
    if not task_id.strip() or not reviewer.strip() or (decision in {"revise", "reject"} and not correction.strip()):
        raise ValueError("Task, reviewer, and a reason for revision/rejection are required")
    record = {"schema_version": "1.0", "feedback_id": str(uuid.uuid4()), "task_id": task_id,
              "timestamp_utc": utc_now(), "reviewer": reviewer, "decision": decision, "correction": correction,
              "source_ids": source_ids or [], "training_approved": False, "model_update_performed": False}
    record["sha256"] = digest(record)
    destination = Path(path)
    destination.parent.mkdir(parents=True, exist_ok=True)
    with destination.open("a", encoding="utf-8") as f:
        f.write(json.dumps(record, ensure_ascii=False, allow_nan=False)+"\n")
    return record
```
