## Supplementary data and reproducibility code for "PanVasc Research for AI assisted evidence analysis in panvascular intervention": README.md

# 泛血管介入科研助手

可在 Windows 本地运行的科研工作台原型，围绕文献检索、PICO 研究设计草案、CSV 描述统计和研究者审阅构建。借鉴 ScienceBuddy 的交互与证据记录思路；当前不是 ScienceBuddy 复现，不包含新训练的专科模型或递归学习实验。具体对标核查见 [ScienceBuddy 调研](docs/sciencebuddy-research.md)，领域设计见 [设计说明](docs/domain-design.md)。

## 已完成的评估与复现入口

桌面工作台与独立本地模型实验分别评估，不能互换结果：

| 评估 | 已完成内容 | 可支持的结论 |
|---|---|---|
| 历史工程演示 | 28 项离线工程检查、文献检索记录和 120 行合成数值示例 | 所检查的软件行为；不代表模型准确率或临床效果 |
| 固定本地模型 pilot | 官方 Qwen3-4B Q8_0 的 700 次测试调用：两种提示各完成 200 个注册资料受控任务、50 个干净来源对照和 100 个 NLI4CT 任务 | 给定来源上的字段一致性，以及既有乳腺癌试验语料上的语义推断表现 |

原始输出和规则检查后的输出复用同一次模型响应，形成 1,400 条评分记录，没有增加模型调用。另有 10 次开发集技术检查调用和 1 次非医学 JSON 烟测，均不计入 700 次测试调用。工程检查、烟测和模型测试不能合并为一个准确率。

协议见 [固定 pilot 协议](experiments/pilot/protocol.md)，真实模型测试汇总见 [test_summary.json](experiments/pilot/runs/test_summary.json)，本地服务与公开包重建方法见 [本地模型说明](docs/LOCAL_MODEL_SETUP.md)。公开补充材料的入口是 [复现说明](shareable_supplement/REPRODUCIBILITY.md)。本地 pilot 通过独立实验脚本执行，尚未接入桌面界面的按钮或聊天流程；桌面的可选远程模型适配器仍未经该 pilot 验证。

## 启动

需要 Python 3.11 或更高版本，安装时保留 Tcl/Tk（tkinter）。在本目录打开终端：

```powershell
python -m venv .venv
.\.venv\Scripts\python.exe -m pip install -r requirements.txt
.\.venv\Scripts\python.exe app.py
```

完成依赖安装后，也可以双击 **启动科研助手.cmd**。启动器优先使用本目录 `.venv`，否则使用系统 `python`。应用仅在手动运行时打开桌面窗口，不启动网站或后台监听服务。

## 使用与复现

1. 选择冠状动脉、外周动脉、颈动脉与脑血管、主动脉、静脉之一。不同领域采用预设 PubMed 检索范围，不能视作全面覆盖该领域。
2. 在“文献检索”填写英文短语；分号分隔的短语以 AND 组合，可选择年份和 1–25 条结果。先调用 PubMed（NCBI E-utilities），连接或解析失败时尝试 Europe PMC 官方 REST 备用接口。界面和任务 JSON 明确记录实际 `api_source`、`fallback_used`、`primary_failure`，并显示实际检索式。两者检索语义不同，备用结果不标成 PubMed 直接查询。可查看真实 PMID、原文标题、摘要与撤稿相关标记。完整查询、时间、元数据和原始 XML/JSON 均保留；无结果或错误不补造文献。
3. 在“PICO 研究草案”填写人群、干预、对照、结局和随访。生成的是可审阅的固定结构模板，需自行确定研究设计与分析决策。
4. 在“本地 CSV 分析”选择 UTF-8 或 GB18030 文件，指定 ID、分组、0/1 结局与可选随访列。可运行缺失/重复/值域等质量检查、描述统计，或两组二元结局比较。原型限制为 50 MB、100,000 行、300 列；列名应非空且唯一。
5. 在“任务、反馈与导出”选择完成的任务，填写审阅者与 accept/revise/reject。修改或拒绝时须填写理由。反馈绑定所选任务的真实 UUID；可导出该任务 JSON。

界面中的联网和分析操作由后台线程执行，主窗口可继续响应。一次只执行一个后台操作，关闭应用会终止未完成的后台工作。新任务始终生成独立记录，不覆盖旧结果。

不弹出窗口的语法检查：

```powershell
.\.venv\Scripts\python.exe -m py_compile app.py panvasc/core.py panvasc/llm.py
```

如已安装依赖，可在本目录验证核心 API（真实文献检索会联网，结果随时间和索引变化）：

```powershell
.\.venv\Scripts\python.exe -c "from panvasc.core import search_literature; r=search_literature('coronary', 'stent', limit=3, output_dir='local_workspace/literature'); print(r['run_id'], r['api_source'], r.get('fallback_used', False), r['count']); print([(x['pmid'], x['title']) for x in r['records']])"
```

程序不会在启动时运行上述联网示例，也不会自动下载模型。请保存实际检索运行的原始 XML/JSON、查询及来源标记，以便说明所使用的文献版本。

## 模式与数据去向

- **默认模板模式：** 未配置模型时，界面明确显示“模型接口未配置”。数据库元数据检索、固定研究模板和统计计算都不被标作模型推理。
- **可选模型模式：** 仅当运行环境已设置 `OPENAI_API_KEY` 和 `PANVASC_MODEL`、取得本次真实公开文献，并在界面明确勾选启用远程生成后，按钮才可用。自行选择账户可用的模型，凭据不得写入共享文件。点击按钮将公开课题问题及本次文献标题/摘要等发送至 `panvasc/llm.py` 指定的 OpenAI 接口；该调用可能产生账户费用。模型草案单独存为“未验证”任务，不能视作已完成证据核实。
- **本地数据：** CSV 不进入模型请求。CSV 分析结果、PICO 字段、模型输出及反馈记录保存在本机；导出的任务可能包含本地文件路径、汇总数据及研究问题，分享前应自行审阅。程序未实现自动去标识。
- **公开文献数据库：** 检索关键词和领域检索式发送给 NCBI；备用检索时亦发送给 Europe PMC。仅输入公开研究主题，不输入个人信息。
- **反馈：** 仅保存审阅意见，`training_approved=false`、`model_update_performed=false`。接受某个输出不等于其科学结论真实，也不触发自动训练。

研究数据与凭据应由使用者按所在机构的授权范围管理。远程问题框只用于公开课题，不应粘贴患者资料或未授权内容。

## 本地文件

```text
local_workspace/
  tasks/<task_id>.json        完成的任务、输入来源与结果
  literature/<run_id>/        原始 ESearch/EFetch 或 Europe PMC 响应与解析结果
  feedback.jsonl             审阅者、任务 ID、决定、理由与摘要校验值
```

应用再次启动时读取已有任务记录；损坏记录被跳过并提示，原文件保留。CSV 表本身不复制到任务目录；其统计结果可能含类别名称，仍应作为研究资料妥善保存。

## 范围与局限

- 领域检索由固定条件构造，返回上限 25 条，不能替代系统综述的完整数据库检索、筛选与偏倚评价。
- 原始摘要不是全文；PMID 存在不意味着特定断言受到支持。应用不自动进行研究质量评级、全文证据抽取或引文蕴含验证。
- PICO 模板要求研究者补足研究设计、估计目标、终点定义、混杂、缺失、聚类与多重比较决策；模板不会自行决定临床策略。
- 二元比较仅计算完整记录的未经调整事件比例、Wilson 95% 区间、双侧 Fisher 精确检验、组间粗风险差及优势比；不建立因果关系，不处理生存删失、竞争风险、重复观测或不等随访。
- 数据质量检查为启发式检查，不能保证去标识、无泄漏或分析有效。描述统计不执行插补或异常值修复。
- 没有专科训练权重、前瞻性临床验证或患者结局证据。软件检查和示例运行不能替代专家盲评、外部数据集评价或临床研究。
- 当前不提供账户权限、多用户协作或机构级审计系统，适用于研究原型与方法开发。
