## Supplementary data and reproducibility code for "PanVasc Research for AI assisted evidence analysis in panvascular intervention": run_alternative_retrieval.py

```
"""Separate, explicitly labeled Europe PMC run following NCBI transport failures."""
import sys,json
from pathlib import Path
ROOT=Path(__file__).resolve().parents[1]
sys.path.insert(0,str(ROOT))
from panvasc.core import DOMAINS,search_europepmc,utc_now
out=ROOT/'artifacts'
results=[]
for domain in DOMAINS:
    print('Europe PMC',domain,flush=True)
    try:
        r=search_europepmc(domain,limit=3,output_dir=out/'retrieval_raw')
        r['metadata_identity_complete']=sum(bool(x['pmid'] and x['title'] and x['url']) for x in r['records'])
    except Exception as e:
        r={'domain':domain,'status':'failed','error':str(e),'timestamp_utc':utc_now(),'records':[],'api_source':'Europe PMC REST'}
    results.append(r)
    (out/'europepmc_retrieval_retry.json').write_text(json.dumps(results,ensure_ascii=False,indent=2),encoding='utf-8')
print(json.dumps([{k:r.get(k) for k in ['domain','status','count','elapsed_seconds','error']} for r in results],indent=2),flush=True)
```
