## Supplementary data and reproducibility code for "PanVasc Research for AI assisted evidence analysis in panvascular intervention": run_demonstration.py

```
"""Run five real metadata retrieval tasks and an explicitly synthetic analysis."""
import sys
from pathlib import Path
ROOT = Path(__file__).resolve().parents[1]
sys.path.insert(0, str(ROOT))
import json
import platform
import hashlib
import importlib.metadata
import numpy as np
import pandas as pd
from panvasc.core import DOMAINS, search_pubmed, audit_dataframe, describe_dataframe, compare_binary, make_protocol, utc_now

def main():
    out = ROOT / "artifacts"
    out.mkdir(exist_ok=True)
    retrieval = []
    for domain in DOMAINS:
        print("Retrieving", domain, flush=True)
        try:
            result = search_pubmed(domain, limit=3, output_dir=out/"retrieval_raw")
            result["metadata_identity_complete"] = sum(bool(r["pmid"] and r["title"] and r["url"]) for r in result["records"])
        except Exception as e:
            result = {"domain":domain, "status":"failed", "error":str(e), "timestamp_utc":utc_now(), "records":[]}
        retrieval.append(result)
        (out/"online_retrieval.json").write_text(json.dumps(retrieval,ensure_ascii=False,indent=2),encoding="utf-8")
    rng = np.random.default_rng(20260919)
    # Fixed fictitious counts selected to illustrate descriptive calculations, not estimated clinical rates.
    data = pd.DataFrame({"patient_id":[f"SYN{i:03d}" for i in range(1,121)],
            "group":["A"]*60+["B"]*60,
            "age_years":rng.integers(40,81,120),
            "event_365d":[1]*8+[0]*52+[1]*12+[0]*48,
            "follow_up_days":[365]*120})
    data.loc[[4,44,91],"age_years"] = np.nan
    path = out/"synthetic_example.csv"
    data.to_csv(path,index=False)
    result = {"data_origin":"Fully synthetic software fixture; no real patients; not a clinical cohort", "seed":20260919,
          "data_sha256":hashlib.sha256(path.read_bytes()).hexdigest(),
          "audit":audit_dataframe(data,outcome="event_365d",time_column="follow_up_days",features=["age_years"]),
          "descriptive":describe_dataframe(data),"binary_comparison":compare_binary(data,"group","event_365d")}
    (out/"synthetic_analysis.json").write_text(json.dumps(result,indent=2,ensure_ascii=False,allow_nan=False),encoding="utf-8")
    protocol=make_protocol("coronary", "adults undergoing repeat intervention for coronary in-stent restenosis", "drug-coated balloon treatment", "repeat drug-eluting stent implantation", "target lesion failure with independently specified components", "12 months")
    (out/"protocol_example.json").write_text(json.dumps(protocol,indent=2,ensure_ascii=False),encoding="utf-8")
    runtime={"timestamp_utc":utc_now(),"python":platform.python_version(),"platform":platform.platform(),
             "packages":{p:importlib.metadata.version(p) for p in ["numpy","pandas","requests","scipy"]},
             "language_model_inference":"not_run", "model_training":"not_performed", "clinical_data":"none"}
    (out/"runtime.json").write_text(json.dumps(runtime,indent=2),encoding="utf-8")
    print(json.dumps({"retrieval":[{k:r.get(k) for k in ["domain","status","count","elapsed_seconds","error"]} for r in retrieval],"synthetic":result["binary_comparison"]},indent=2),flush=True)

if __name__=="__main__":main()
```
