## Supplementary data and reproducibility code for "PanVasc Research for AI assisted evidence analysis in panvascular intervention": run_validation.py

```
"""Run offline software checks and write real per-test JSON/CSV results."""
from __future__ import annotations

import csv
import hashlib
import importlib.metadata
import json
import platform
import sys
import time
import unittest
from datetime import datetime, timezone
from pathlib import Path

ROOT = Path(__file__).resolve().parents[1]
sys.path.insert(0, str(ROOT))

from panvasc import __version__
from tests.test_core import CoreAcceptanceTests


class RecordedResult(unittest.TextTestResult):
    def __init__(self, *args, **kwargs):
        super().__init__(*args, **kwargs)
        self.records = []
        self.started = {}

    def startTest(self, test):
        self.started[test.id()] = time.perf_counter()
        super().startTest(test)

    def record(self, test, status, detail=""):
        self.records.append({"id": test._testMethodName.split("_", 2)[1],
                             "test": test.id(), "expectation": test.shortDescription(),
                             "status": status, "elapsed_seconds": round(time.perf_counter() - self.started[test.id()], 6),
                             "detail": detail})

    def addSuccess(self, test):
        self.record(test, "passed")
        super().addSuccess(test)

    def addFailure(self, test, err):
        self.record(test, "failed", self._exc_info_to_string(err, test))
        super().addFailure(test, err)

    def addError(self, test, err):
        self.record(test, "error", self._exc_info_to_string(err, test))
        super().addError(test, err)

    def addSkip(self, test, reason):
        self.record(test, "skipped", reason)
        super().addSkip(test, reason)


def sha(path):
    return hashlib.sha256(path.read_bytes()).hexdigest()


def main():
    start = time.perf_counter()
    suite = unittest.defaultTestLoader.loadTestsFromTestCase(CoreAcceptanceTests)
    result = unittest.TextTestRunner(verbosity=2, resultclass=RecordedResult).run(suite)
    counts = {status: sum(r["status"] == status for r in result.records)
              for status in ("passed", "failed", "error", "skipped")}
    report = {
        "schema_version": "1.0",
        "evaluation_type": "offline_software_acceptance",
        "software_version": __version__,
        "timestamp_utc": datetime.now(timezone.utc).isoformat(),
        "runtime": {"python": platform.python_version(), "platform": platform.platform(),
                    "dependencies": {name: importlib.metadata.version(name) for name in ("numpy", "pandas", "requests", "scipy")}},
        "source_hashes_sha256": {name: sha(ROOT / name) for name in ("panvasc/core.py", "tests/test_core.py", "scripts/run_validation.py")},
        "summary": {"total": result.testsRun, **counts, "elapsed_seconds": round(time.perf_counter() - start, 6)},
        "interpretation": "Convenience software acceptance fixtures developed in the same implementation cycle; not independent blinded evaluation or clinical validation.",
        "data": "Artificial tabular/XML fixtures only. All network operations in this suite are mocked. No real patient data or trained-model outputs.",
        "not_evaluated": [
            "Clinical efficacy, patient outcomes, diagnosis, or treatment recommendations",
            "LLM or trained model accuracy and improvement after expert feedback",
            "Search relevance, retrieval sensitivity, source entailment, or systematic review completeness",
            "Unit recognition/conversion, complete temporal leakage detection, or de-identification",
            "Immutable external audit log, artifact-version binding, or idempotent feedback replay",
            "Adversarial robustness outside the single instruction-shaped abstract fixture",
            "Online PubMed/Europe PMC availability/performance (reported by separate experiments)",
        ],
        "notes": [
            "Cases T01-T28 are implementation-level checks, not a claim that every target D01-D20 in domain-design.md is implemented.",
            "T25-T28 are regression checks added after review identified blank identifiers, fully missing outcomes, and numeric identifier representation gaps; the implementation was corrected before this final run.",
            "Descriptive quartiles use pandas default linear interpolation; SD uses ddof=1.",
            "The timeout case documents current fail-fast behavior, not timeout retry support.",
            "A stored feedback content hash is a consistency check; it does not prevent a privileged editor rewriting the file and hash.",
        ],
        "cases": sorted(result.records, key=lambda r: r["id"]),
    }
    destination = ROOT / "artifacts"
    destination.mkdir(parents=True, exist_ok=True)
    (destination / "offline_validation.json").write_text(json.dumps(report, ensure_ascii=False, indent=2), encoding="utf-8")
    with (destination / "offline_validation.csv").open("w", encoding="utf-8-sig", newline="") as handle:
        writer = csv.DictWriter(handle, fieldnames=["id", "test", "expectation", "status", "elapsed_seconds", "detail"])
        writer.writeheader()
        writer.writerows(report["cases"])
    print(json.dumps(report["summary"]))
    return 0 if result.wasSuccessful() else 1


if __name__ == "__main__":
    raise SystemExit(main())
```
