## Supplementary data and reproducibility code for "PanVasc Research for AI assisted evidence analysis in panvascular intervention": LOCAL_MODEL_SETUP.md

# Local model setup

## 给使用者的说明

本项目原始实验电脑上配置了本地模型服务：它读取官方 Qwen3-4B 权重，并通过本机接口接收请求、返回文本。本次工作使用现成的预训练模型，没有训练、微调或修改模型权重，也没有接入外部付费推理服务。此说明不意味着另一台电脑已完成安装。

这个服务与项目中的“科研助手”界面是两个部分。本次配置尚未把模型接入该界面的按钮或聊天流程；启动模型服务不会自动增加界面功能。固定 pilot 使用单独的实验脚本调用模型，界面接入需要另行实现。

### 已配置的原始工作区：日常启动和停止

以下步骤适用于已下载并核验权重、保留本机 `runtime/local_llm/setup_manifest.json` 的原始工作区。公开补充包不包含这些运行文件；首次在其他电脑复现时，先按下方“Public-package reconstruction”准备。在 PowerShell 中进入自己的项目目录后，使用以下命令启动模型：

```powershell
# Run from the project root containing scripts/ and runtime/.
.\scripts\start_local_model.ps1
```

模型会在后台加载，随后提供 `http://127.0.0.1:18123` 本机服务。启动命令打印的进程编号、端口和日志位置也会保存在 `runtime/local_llm/active_server.json` 中。服务已运行时无需重复启动；如果端口已被占用，脚本会停止并提示，不会替换已有进程。关闭 PowerShell 窗口不等于停止后台模型。

实验结束且不再需要模型时，使用以下命令停止本项目记录的服务：

```powershell
.\scripts\stop_local_model.ps1
```

正在执行实验时应保持服务运行。模型权重已经下载并核验后，日常启动只需启动脚本，无需重新下载或再次运行配置冻结脚本。模型下载和缓存位于 `models`，运行程序及本地记录位于 `runtime/local_llm`，两者均不进入 Git。

### 技术烟测与固定 pilot 的区别

技术烟测只要求模型返回 `{"ok": true}`，用于确认本机接口能够生成并返回 JSON。它没有使用研究数据，其成功不代表模型具有医学准确性，也不计入研究结果。

固定 pilot 按 `experiments/pilot/protocol.md` 和 `execution_freeze.json` 中事先保存的模型、输入、提示词和评分规则执行。它比较两种固定提示方式，并对同一份输出分别进行原始评分和规则检查，不会在测试过程中训练模型、修改提示词或让模型反复改答案。注册资料任务用于检查字段与给定来源是否一致；NLI4CT 用于既有乳腺癌试验资料上的语义推断，不能据此证明血管领域临床能力。

本次已完成 700 次测试调用：两种提示各执行 200 个注册资料受控任务、50 个干净来源对照和 100 个 NLI4CT 任务。每份响应同时用于原始评分与规则检查后的评分，共 1,400 条评分记录，没有额外模型调用。此前的 28 项工程检查、10 次开发集技术检查调用和一次非医学 JSON 烟测分别记录，均不计入这 700 次测试调用。

研究结果见 `experiments/pilot/runs/test_summary.json` 及相应冻结记录；公开补充包中的对应路径为 `pilot/code/experiments/pilot/runs/test_summary.json`。不能把服务启动成功、工程测试通过或技术烟测输出当作模型实验结果。独立复现应遵循协议重新准备数据并建立自己的冻结记录，保留与本次历史数据和配置的差异。

## Technical configuration record

The isolated runtime is stored in `runtime/local_llm`; official model weights are in `models/Qwen3-4B-GGUF`. Both directories are excluded from Git. No global packages, GPU drivers, credentials, or paid inference services are used.

On the original machine, `scripts/setup_local_model.py` downloaded official, revision-pinned assets through its existing system proxy, validated complete SHA256 values, saved official metadata and licenses, and extracted the portable runtime. It recorded unsuccessful downloads in `runtime/local_llm/download_failures.jsonl` and supported resumable byte-range downloads for the model. This machine-specific helper is not in the public pilot package; its proxy setting is not a reproduction requirement or an inference endpoint.

- Runtime: ggml-org/llama.cpp build b10964 (recorded through stable release v0.4.1 at setup); commit `b29c606e28a01b1bc8c1351026a0fa6e616bf6c4`; Windows x64 Vulkan build; MIT license.
- Model: official Qwen/Qwen3-4B-GGUF; revision `bc640142c66e1fdd12af0bd68f40445458f3869b`; Q8_0 quantization; Apache-2.0 license.
- Model SHA256: `8c2f07f26af9747e41988551106f149b03eb9b5cb6df636027b6bf6278473300`.
- Runtime archive SHA256: `1ee3ad952f4ba71f438bd6d7bebef19e1c7af04adcaa35d08b4ddabb27d4c642`.

Start a hidden, manually stoppable background process with `scripts/start_local_model.ps1`. The default binding is **127.0.0.1:18123**; an existing listener is never replaced. Defaults are one parallel slot, 8192-token context, 99 GPU layers, eight CPU threads, the embedded Jinja chat template, and no context shifting. The context and port can be supplied as script parameters. Sampling settings belong to each experiment request and are not chosen by the startup script.

The OpenAI-compatible endpoint is `http://127.0.0.1:18123/v1/chat/completions`, with model alias `Qwen3-4B-Q8_0`. Health is available at `http://127.0.0.1:18123/health`. The exact process ID, command arguments and logs are recorded in `runtime/local_llm/active_server.json`. Stop the recorded process with `scripts/stop_local_model.ps1`; the stop script checks the executable and process start time before terminating it.

The original machine's `scripts/smoke_test_local_model.py` sent one non-medical technical prompt requesting `{"ok": true}` and saved the request, complete actual response, health reply, and outcome. It disabled thinking through `chat_template_kwargs`, used seed 20260919, temperature 0 and a 32-token limit, and did not read experiment data. This helper is not included in the public pilot package. The public provenance file retains the allowlisted smoke-test outcome. A passed smoke test demonstrates only that the local service returns JSON; it is not evidence of medical accuracy or study performance.

The original `runtime/local_llm/setup_manifest.json` is the authoritative local record of download status, verified artifacts, exact source and license URLs, startup settings and the actual smoke-test result. It is omitted from the public package because it includes machine-specific paths and network settings; `pilot/model_provenance.json` supplies selected public fields. A downloaded or verified file is not itself an executed model result.

## Public-package reconstruction

The public pilot package includes `start_local_model.ps1` and `stop_local_model.ps1`, but no automatic installer, model weights, runtime binary or ready-to-use local setup manifest. Use a separate working copy of `pilot/code/` as the project root and preserve the distributed audit records unchanged elsewhere. Install its `requirements-pilot.txt` in an isolated Python environment. The historical engineering environment has separate package versions and should not be substituted for the pilot environment recorded in its execution freeze.

1. Obtain [llama.cpp build b10964](https://github.com/ggml-org/llama.cpp/releases/tag/b10964), specifically `llama-b10964-bin-win-vulkan-x64.zip`, and the official [Qwen3-4B Q8_0 file at the frozen revision](https://huggingface.co/Qwen/Qwen3-4B-GGUF/blob/bc640142c66e1fdd12af0bd68f40445458f3869b/Qwen3-4B-Q8_0.gguf). Keep HTTPS certificate verification enabled. Compare the full downloaded files against the hashes above and `pilot/model_provenance.json`; a different asset, revision or quantization is a different reproduction configuration.
2. Put the model at `models/Qwen3-4B-GGUF/Qwen3-4B-Q8_0.gguf`, retain the ZIP under `runtime/local_llm/downloads/`, and extract all runtime files together under `runtime/local_llm/server/`. If the archive has a nested directory, adjust the executable path in the example below. Preserve its bundled libraries and license. Do not assume another computer has the original GPU or compatible Vulkan configuration; record hardware or backend changes.
3. The startup wrapper reads `runtime.server_executable`, `runtime.download.verified`, `model.download.path` and `model.download.verified` from your own `runtime/local_llm/setup_manifest.json`. Create that local file only after verification. The following PowerShell example checks both supplied file hashes before writing the minimal launch record; it does not claim a smoke test or infer scientific success:

```powershell
# Run from your separate reproduction project's root.
$pilotRuntimeZip = (Resolve-Path '.\runtime\local_llm\downloads\llama-b10964-bin-win-vulkan-x64.zip').Path
$pilotModelPath = (Resolve-Path '.\models\Qwen3-4B-GGUF\Qwen3-4B-Q8_0.gguf').Path
$pilotServerPath = (Resolve-Path '.\runtime\local_llm\server\llama-server.exe').Path
$pilotRuntimeHash = (Get-FileHash -LiteralPath $pilotRuntimeZip -Algorithm SHA256).Hash.ToLowerInvariant()
$pilotModelHash = (Get-FileHash -LiteralPath $pilotModelPath -Algorithm SHA256).Hash.ToLowerInvariant()
if ($pilotRuntimeHash -ne '1ee3ad952f4ba71f438bd6d7bebef19e1c7af04adcaa35d08b4ddabb27d4c642') { throw 'Runtime archive hash mismatch' }
if ($pilotModelHash -ne '8c2f07f26af9747e41988551106f149b03eb9b5cb6df636027b6bf6278473300') { throw 'Model hash mismatch' }
$pilotSetupRecord = [ordered]@{
    purpose = 'Independent reconstruction; verified downloads, no inference claim'
    created_utc = (Get-Date).ToUniversalTime().ToString('o')
    runtime = @{ tag = 'b10964'; server_executable = $pilotServerPath; download = @{ path = $pilotRuntimeZip; sha256 = $pilotRuntimeHash; verified = $true } }
    model = @{ revision = 'bc640142c66e1fdd12af0bd68f40445458f3869b'; download = @{ path = $pilotModelPath; sha256 = $pilotModelHash; verified = $true } }
}
$pilotSetupRecord | ConvertTo-Json -Depth 6 | Set-Content -LiteralPath '.\runtime\local_llm\setup_manifest.json' -Encoding utf8
```

4. Start the included wrapper, wait for its health endpoint to respond, and retain the actual server version, local hardware, arguments and logs. Stop it with the included stop script when finished. The startup wrapper verifies the recorded paths and verification flags; it does not itself rehash all artifacts on every launch. Keep the independently computed hashes and original public provenance with your new run.
5. Follow `experiments/pilot/protocol.md` and the registry/NLI4CT acquisition READMEs to obtain the omitted source data. Prepare inputs, run the independent verification script, and create a new execution freeze before test calls. The runner's inference and evaluation commands are documented in the public pilot README. Do not reuse the published freeze with changed paths, source hashes or local manifests. Registry updates can change the sampled records and source text, so a new live acquisition may reproduce the procedure without reproducing the historical snapshot or numerical results.

These are reconstruction instructions for a new run, not a claim that the public package alone contains all original inputs or can replay the reported outputs offline.
