## Supplementary data and reproducibility code for "PanVasc Research for AI assisted evidence analysis in panvascular intervention": metric_notes.md

# Official evidence scoring and connected clusters

The official `Task_2_Evaluation.zip` was downloaded from the same fixed upstream revision as the data. Archive SHA-256: `64f622486dfdc6b9653407c4443d59697950fe6cf20ce65c4ab167c124628db8`. Its `Task_2_Evaluation/evaluate.py` aggregates true positives, false positives and false negatives across all primary and secondary evidence arrays, then computes global precision, recall and F1. This is positive-evidence micro scoring, not a mean of per-example F1 values. Prefer the actual official evaluation script when describing the scoring convention; prose references to macro scoring should not overwrite it.

Represent evidence atoms as `(example_id, trial_id, zero_based_line_index)`, not just `(trial_id,index)` across the whole dataset, because two statements about the same trial can require different evidence. For each example, translate the original primary/secondary role to its NCT and form predicted and gold sets. Sum TP/FP/FN over examples, with every selected example retained. Compute `precision=TP/(TP+FP)`, `recall=TP/(TP+FN)`, `F1=2*TP/(2*TP+FP+FN)`; define zero denominators explicitly, normally as zero. Evidence indices refer to the original section-line positions, including headings.

The official small evaluation script has three implementation limitations: it iterates only prediction IDs, does not deduplicate repeated prediction indices, and does not guard zero denominators. A robust local implementation should retain all 100 selected IDs, score missing/invalid outputs as empty evidence, and either reject duplicate indices or deduplicate before set scoring. Report these integrity safeguards, because blindly using the historical loop can reward missing cases or repeated indices. Do not select only accepted/answered examples for the unconditional evidence metric.

A valid in-range evidence index is not necessarily relevant evidence. An input-only guard can check ID/section/index syntax and ranges but must not read gold indices. Report raw evidence performance and guarded performance separately, including coverage. Correct labels with incorrect evidence remain distinct failures. Official gold labels are not changed or supplemented by this project.

`test_nct_clusters.json` and `dev_nct_clusters.json` connect selected examples through any shared primary/secondary NCT. A Comparison example joins both of its NCT nodes; connected chains remain one component. Use component-resampling paired bootstrap with all examples in a sampled component retained, and identical component draws for both prompt arms. Recompute the metric on the concatenated examples in each draw. This preserves the original example-weighted point estimate; component-weighted performance would be a different estimand and must be named separately. These are source-sharing clusters, not newly adjudicated trial families. Report the actual component count and largest component.

The official candidate splits have no NCT overlap between dev and test in this fixed release. This is distinct from model pretraining contamination, which cannot be excluded for a long-public benchmark.

Source: [Fixed official Task 2 evaluation archive](https://github.com/ai-systems/nli4ct/blob/8e8e5fc506b916984011873403b9ea5ed6dd96c2/Task_2_Evaluation.zip).

The fixed official Task 1 script was also inspected. It maps Entailment to 1 and all other supplied strings to 0, then calls scikit-learn binary `f1_score`, `precision_score` and `recall_score` with their default positive class. Thus its original leaderboard F1 is positive-Entailment F1, not the proposed pilot's primary accuracy or secondary two-label macro F1. Those pilot metrics are reasonable if named explicitly, but should not be equated with the historical official score. In particular, malformed model outputs should not silently become correct Contradiction predictions. Keep invalid output status separate and count it as wrong for unconditional accuracy; retain all reference IDs. Any macro-F1 handling of invalid outputs must state that the macro average is over the two official labels, with invalid predictions contributing false negatives rather than a third target class.

Source: [Fixed official Task 1 evaluation archive](https://github.com/ai-systems/nli4ct/blob/8e8e5fc506b916984011873403b9ea5ed6dd96c2/Task_1_Evaluation.zip).
