## Supplementary data and reproducibility code for "PanVasc Research for AI assisted evidence analysis in panvascular intervention": prepare_data.py

```
"""Freeze an untouched official NLI4CT subset; no model calls or new labels."""
from __future__ import annotations

import collections
from datetime import datetime, timezone
import hashlib
import json
from pathlib import Path
import random
import urllib.request
from zipfile import ZipFile

ROOT = Path(__file__).resolve().parent
REVISION = '8e8e5fc506b916984011873403b9ea5ed6dd96c2'
ARCHIVE_SHA256 = 'dd972c46e69a5bc5e5fb74c2105b5444722d81879ab96b214545b943e4e7ebbe'
BASE = f'https://raw.githubusercontent.com/ai-systems/nli4ct/{REVISION}'
SEED = 20260919
LABELS = ('Contradiction', 'Entailment')
GOLD_KEYS = {'Label', 'Primary_evidence_index', 'Secondary_evidence_index'}

def digest(data: bytes) -> str:
    return hashlib.sha256(data).hexdigest()

def write_json(path: Path, obj) -> None:
    path.parent.mkdir(parents=True, exist_ok=True)
    path.write_text(json.dumps(obj, ensure_ascii=False, indent=2)+'\n', encoding='utf-8')

def write_jsonl(path: Path, rows) -> None:
    path.parent.mkdir(parents=True, exist_ok=True)
    path.write_text(''.join(json.dumps(row, ensure_ascii=False, sort_keys=True)+'\n' for row in rows), encoding='utf-8')

def source_ids(record: dict) -> set[str]:
    return {record[k] for k in ('Primary_id', 'Secondary_id') if record.get(k)}

def selected_ids(data: dict, per_label: int) -> list[str]:
    # Independent streams start from the same seed for each official split.
    rng = random.Random(SEED)
    chosen = []
    for label in LABELS:
        population = sorted(k for k, v in data.items() if v['Label'] == label)
        if len(population) < per_label:
            raise ValueError(f'Insufficient official {label} records')
        chosen.extend(rng.sample(population, per_label))
    return sorted(chosen)

def freeze() -> dict:
    stamp = datetime.now(timezone.utc).isoformat()
    source = ROOT/'source'
    source.mkdir(parents=True, exist_ok=True)
    archive = source/'Complete_dataset.zip'
    if not archive.exists():
        request = urllib.request.Request(BASE+'/Complete_dataset.zip', headers={'User-Agent':'PanVascResearch-data-preparation'})
        with urllib.request.urlopen(request, timeout=90) as response:
            archive.write_bytes(response.read())
    if digest(archive.read_bytes()) != ARCHIVE_SHA256:
        raise ValueError('The archive differs from the fixed official revision checksum')
    readme = source/'README.upstream.md'
    if not readme.exists():
        with urllib.request.urlopen(BASE+'/README.md', timeout=30) as response:
            readme.write_bytes(response.read())

    member_hashes = {}
    with ZipFile(archive) as zipped:
        def read_member(name):
            raw = zipped.read(name)
            member_hashes[name] = digest(raw)
            return json.loads(raw)
        train = read_member('Complete_dataset/train.json')
        dev = read_member('Complete_dataset/dev.json')
        test = read_member('Complete_dataset/test.json')
        test_gold = read_member('Complete_dataset/Gold_test.json')
        if set(test) != set(test_gold):
            raise ValueError('Official test and Gold_test IDs differ')
        for case_id, record in test.items():
            if record != {k:v for k,v in test_gold[case_id].items() if k not in GOLD_KEYS}:
                raise ValueError(f'Gold_test changes the official test input: {case_id}')
        if set(dev) & set(test):
            raise ValueError('Official dev/test IDs overlap')
        dev_nct = set().union(*(source_ids(v) for v in dev.values()))
        test_nct = set().union(*(source_ids(v) for v in test.values()))
        if dev_nct & test_nct:
            raise ValueError('Official dev/test NCT IDs overlap; freeze needs an explicit revised selection rule')

        selections = {'dev': selected_ids(dev, 10), 'test': selected_ids(test_gold, 50)}
        rows_by_split = {}
        expected_by_split = {}
        source_cache = {}
        for split, gold_data in [('dev', dev), ('test', test_gold)]:
            input_rows, expected_rows = [], []
            for case_id in selections[split]:
                record = gold_data[case_id]
                input_record = dev[case_id] if split == 'dev' else test[case_id]
                sources = []
                evidence = {}
                for official_role, role in [('Primary','primary'), ('Secondary','secondary')]:
                    nct = record.get(official_role+'_id')
                    if not nct:
                        continue
                    if nct not in source_cache:
                        source_cache[nct] = read_member('Complete_dataset/CT json/'+nct+'.json')
                    ctr = source_cache[nct]
                    if ctr['Clinical Trial ID'] != nct:
                        raise ValueError('Trial ID disagrees with source filename')
                    section = record['Section_id']
                    lines = ctr[section]
                    if not isinstance(lines, list) or not all(isinstance(x, str) for x in lines):
                        raise ValueError('Source section is not a list of unmodified strings')
                    indices = record[official_role+'_evidence_index']
                    if not all(isinstance(i,int) and not isinstance(i,bool) and 0 <= i < len(lines) for i in indices):
                        raise ValueError(f'Out-of-range official evidence index in {case_id}')
                    sources.append({'role':role, 'trial_id':nct, 'section':section,
                        'lines':[{'index':i,'text':text} for i,text in enumerate(lines)]})
                    evidence[role] = list(indices)
                row = {'id':case_id, 'benchmark':'NLI4CT SemEval 2023', 'official_split':split,
                    'type':input_record['Type'], 'section':input_record['Section_id'],
                    'statement':input_record['Statement'], 'sources':sources}
                if set(row) != {'id','benchmark','official_split','type','section','statement','sources'}:
                    raise ValueError('Unexpected input schema')
                input_rows.append(row)
                expected_rows.append({'id':case_id,'official_split':split,'label':record['Label'],
                    'evidence_indices':evidence,
                    'source_trial_ids':{s['role']:s['trial_id'] for s in sources},
                    'section':record['Section_id'],
                    'annotation_origin':'Unmodified official dev.json' if split=='dev' else 'Unmodified official Gold_test.json'})
            rows_by_split[split] = input_rows
            expected_by_split[split] = expected_rows
            write_jsonl(ROOT/f'{split}_inputs.jsonl',input_rows)
            write_jsonl(ROOT/'reference'/f'{split}_expected.jsonl',expected_rows)

        inventory = []
        all_reference_index = []
        for split, dataset in [('dev',dev),('test',test_gold)]:
            selected = set(selections[split])
            for case_id, r in sorted(dataset.items()):
                inventory.append({'id':case_id,'official_split':split,'type':r['Type'],'section':r['Section_id'],
                    'source_trial_ids':sorted(source_ids(r)),'selected':case_id in selected,
                    'selection_status':'selected_by_fixed_seed_label_stratum' if case_id in selected else 'not_selected_by_fixed_seed'})
                all_reference_index.append({'id':case_id,'official_split':split,'label':r['Label'],
                    'Primary_evidence_index':r['Primary_evidence_index'],
                    **({'Secondary_evidence_index':r['Secondary_evidence_index']} if 'Secondary_evidence_index' in r else {})})
        write_jsonl(ROOT/'candidate_inventory.jsonl', inventory)
        write_jsonl(ROOT/'reference'/'all_official_reference_index.jsonl',all_reference_index)
        write_json(ROOT/'selection.json',{'seed':SEED,'labels_in_sampling_order':list(LABELS),
            'algorithm':'For each official split independently: Python random.Random(20260919); for Contradiction then Entailment, sample from lexicographically sorted UUIDs; concatenate and sort selected UUIDs. No outcome, length, section or model-output filtering.',
            'requested_counts':{'dev':20,'test':100},'per_label_counts':{'dev':10,'test':50},
            'selected_ids':selections,'excluded_as_invalid':[]})

        cluster_summaries = {}
        for split, inputs in rows_by_split.items():
            parent = {s['trial_id']:s['trial_id'] for r in inputs for s in r['sources']}
            def find(nct):
                while parent[nct] != nct:
                    parent[nct] = parent[parent[nct]]
                    nct = parent[nct]
                return nct
            for row in inputs:
                trials = [s['trial_id'] for s in row['sources']]
                for trial in trials[1:]:
                    a,b = find(trials[0]),find(trial)
                    parent[max(a,b)] = min(a,b)
            components = {}
            for trial in sorted(parent):
                components.setdefault(find(trial), {'trial_ids':[], 'example_ids':[]})['trial_ids'].append(trial)
            assignments = []
            for row in inputs:
                representative = find(row['sources'][0]['trial_id'])
                components[representative]['example_ids'].append(row['id'])
                assignments.append({'id':row['id'],'cluster_id':split+'-'+representative})
            output_components = [{'cluster_id':split+'-'+key,**value,
                'n_trials':len(value['trial_ids']),'n_examples':len(value['example_ids'])}
                for key,value in sorted(components.items())]
            cluster_summaries[split] = {'connected_components':len(output_components),
                'largest_component_examples':max(c['n_examples'] for c in output_components),
                'largest_component_trials':max(c['n_trials'] for c in output_components)}
            write_json(ROOT/f'{split}_nct_clusters.json', {'rule':'Connected components of selected examples sharing a primary or secondary NCT ID; all trial IDs in a Comparison example are joined. No gold label or evidence is used.',
                'scope':'Shared-source dependence only; not adjudicated biological trial families.',
                'assignments':assignments,'components':output_components,'summary':cluster_summaries[split]})

        archive_license_members = [name for name in zipped.namelist() if 'license' in name.lower() or 'licence' in name.lower() or 'copyright' in name.lower()]

    license_review = {
        'checked_at_utc':stamp,
        'repository':'https://github.com/ai-systems/nli4ct',
        'revision':REVISION,
        'repository_api_license':None,
        'repository_license_file_present':False,
        'archive_license_members':archive_license_members,
        'named_license_status':'not_explicitly_specified_in_official_repository_or_archive',
        'official_release_basis':'Official task website explicitly offers Complete_dataset.zip for the task; repository README explicitly announces train, development, test and Gold Test availability.',
        'official_download_page':'https://sites.google.com/view/nli4ct/semeval-2023/get-data-and-starting-kit',
        'official_format_page':'https://sites.google.com/view/nli4ct/semeval-2023/data-formatting',
        'competition_terms_page':'https://codalab.lisn.upsaclay.fr/competitions/8937#learn_the_details-terms_and_conditions',
        'competition_terms_observation':'Terms require use and redistribution according to the data licence but the inspected terms do not identify a specific licence. No competition submission or acceptance of terms was performed.',
        'use_scope':'Local preparation and scholarly benchmark evaluation from the official public task release; no public redistribution or commercial-use grant is asserted.',
        'redistribution_status':'not_cleared_for_repackaging; distribute preparation code, record IDs, hashes and original download instructions instead until a primary-source licence is confirmed',
        'third_party_mirror_license_adopted':False,
        'paper_license_is_dataset_license':False,
    }
    write_json(ROOT/'license_review.json',license_review)

    summary = {}
    for split in ['dev','test']:
        inputs=rows_by_split[split]
        expected=expected_by_split[split]
        summary[split]={'count':len(inputs),
            'labels':dict(collections.Counter(r['label'] for r in expected)),
            'types':dict(collections.Counter(r['type'] for r in inputs)),
            'sections':dict(collections.Counter(r['section'] for r in inputs)),
            'unique_nct_ids':len({s['trial_id'] for r in inputs for s in r['sources']})}
    official_counts={'train':len(train),'dev':len(dev),'test':len(test),'Gold_test':len(test_gold)}
    checks={'official_test_gold_ids_identical':set(test)==set(test_gold),
        'official_test_inputs_identical_to_gold_without_answers':True,
        'official_dev_test_example_disjoint':not bool(set(dev)&set(test)),
        'official_dev_test_nct_disjoint':not bool(dev_nct&test_nct),
        'selected_counts_correct':len(rows_by_split['dev'])==20 and len(rows_by_split['test'])==100,
        'selected_labels_balanced':all(set(s['labels'].values())=={10 if split=='dev' else 50} for split,s in summary.items()),
        'official_evidence_indices_in_range':True,
        'input_fields_do_not_include_gold':True,
        'statements_and_source_lines_unmodified':True,
        'no_model_calls':True,
        'no_new_annotations':True}
    write_json(ROOT/'preparation_checks.json',{'checked_at_utc':stamp,'status':'passed','checks':checks,
        'official_counts':official_counts,'selected_summary':summary,
        'scope':'Data integrity and separation only; not model performance or new annotation validation.'})
    manifest={'schema_version':'1.0','frozen_at_utc':stamp,
        'source_repository':'https://github.com/ai-systems/nli4ct','source_revision':REVISION,
        'source_revision_commit_time_utc':'2024-04-21T13:54:00Z',
        'archive_url':BASE+'/Complete_dataset.zip','archive_bytes':archive.stat().st_size,
        'archive_sha256':ARCHIVE_SHA256,'seed':SEED,'official_counts':official_counts,
        'selected_summary':summary,'checks':checks,
        'cluster_summary':cluster_summaries,
        'source_archive_member_sha256':member_hashes,
        'inference_input_files':['dev_inputs.jsonl','test_inputs.jsonl'],
        'reference_only_files':['reference/dev_expected.jsonl','reference/test_expected.jsonl','reference/all_official_reference_index.jsonl'],
        'inference_boundary':'Model runners may read only *_inputs.jsonl; never load reference/, Gold_test, the source archive, or all_official_reference_index into prompts.',
        'annotation_origin':'Original official NLI4CT annotations only; no new expert labels or adjudication were created.',
        'domain':'Breast cancer clinical trial reports; not panvascular validation.',
        'public_benchmark_contamination':'Long-public benchmark; inclusion in model pretraining cannot be excluded. Do not claim previously unseen evaluation.',
        'subsample_scope':'Fixed, label-stratified official-test subset; results will not equal a full official leaderboard evaluation.',
        'license_status':license_review['named_license_status'],
        'redistribution_status':license_review['redistribution_status'],
        'model_inference':'not_run','prompt_tuning':'not_performed_by_preparation_script',
        'files_sha256':{str(p.relative_to(ROOT)).replace('\\','/'):digest(p.read_bytes()) for p in sorted(ROOT.rglob('*')) if p.is_file() and p.name!='frozen_manifest.json' and '__pycache__' not in p.parts}}
    write_json(ROOT/'frozen_manifest.json',manifest)
    return {'status':'prepared','source_revision':REVISION,'archive_sha256':ARCHIVE_SHA256,
        'selected_summary':summary,'all_checks_passed':all(checks.values()),'license_status':license_review['named_license_status']}

if __name__=='__main__':
    print(json.dumps(freeze(),ensure_ascii=False,indent=2))
```
