## Supplementary data and reproducibility code for "PanVasc Research for AI assisted evidence analysis in panvascular intervention": README.md

# Frozen NLI4CT evaluation inputs

This directory prepares an untouched subset of the official SemEval 2023 NLI4CT release. It contains no model outputs, no tuned prompts, and no new annotations. The source domain is breast cancer clinical trial reports, not panvascular research.

The official repository revision is `8e8e5fc506b916984011873403b9ea5ed6dd96c2` (2024-04-21). The original `Complete_dataset.zip` is retained locally. Its SHA-256 is `dd972c46e69a5bc5e5fb74c2105b5444722d81879ab96b214545b943e4e7ebbe`.

## Frozen selection

- Development: 20 examples drawn only from official `dev.json`, 10 per label.
- Test: 100 examples drawn only from official `test.json`, with the matching original `Gold_test.json` used solely for stratification and reference answers, 50 per label.
- Seed: `20260919`. For each split independently, start `random.Random(20260919)`, sample sorted UUIDs for `Contradiction`, then `Entailment`, and sort the selected UUIDs. No filtering uses model behavior, text length, section, or apparent difficulty.
- Full candidate inventories for all 200 official dev and 500 official test examples are retained. Official train is inspected only for the release count and remains unused. The complete dev/test example IDs and NCT source IDs are disjoint.
- The published release has 1,700 train, 200 dev and 500 test examples. Record the actual release counts rather than replacing them with an approximate split ratio from a paper.

## Inference schema

Read only `dev_inputs.jsonl` or `test_inputs.jsonl` in a model runner. Each line is an object with these fields:

```json
{
  "id": "official UUID",
  "benchmark": "NLI4CT SemEval 2023",
  "official_split": "test",
  "type": "Single or Comparison",
  "section": "official section name",
  "statement": "unchanged official statement",
  "sources": [
    {
      "role": "primary or secondary",
      "trial_id": "official NCT ID",
      "section": "official section name",
      "lines": [{"index": 0, "text": "unchanged source line"}]
    }
  ]
}
```

`sources` has one item for a Single example and two for a Comparison example. `index` is the original zero-based position within the selected section. Leading spaces and source text are preserved. Do not renumber after removing headings, blank lines, or other entries; such removal would invalidate the official evidence indices. All section lines are included, not just gold evidence. The original statement's primary/secondary trial roles remain intact.

Labels and evidence are **absent from input objects**. Do not load `reference/`, the source ZIP, or `all_official_reference_index.jsonl` into prompts. Do not select evidence lines before inference using the gold index.

## Reference schema

`reference/dev_expected.jsonl` and `reference/test_expected.jsonl` contain:

```json
{
  "id": "same official UUID",
  "official_split": "test",
  "label": "Entailment or Contradiction",
  "evidence_indices": {"primary": [0], "secondary": [1]},
  "source_trial_ids": {"primary": "NCT ID", "secondary": "NCT ID"},
  "section": "official section name",
  "annotation_origin": "Unmodified official Gold_test.json"
}
```

The `secondary` entry is absent for Single examples. Labels and evidence index arrays are copied from official files without reinterpretation, relabeling or adjudication. A compatible model response could use `label` and `evidence_indices`, but this preparation does not choose or tune the prompt.

## Files and checks

- `candidate_inventory.jsonl`: all official dev/test UUIDs, type, section, source IDs, selection flags; no gold labels.
- `reference/all_official_reference_index.jsonl`: separate full label/evidence index inventory used for stratification audit.
- `selection.json`: exact IDs, seed and sampling procedure.
- `preparation_checks.json`: integrity, source-ID separation, evidence range, balance and counts.
- `frozen_manifest.json`: source revision, ZIP/member hashes, file hashes, time and limitations.
- `license_review.json`: primary-source license inspection and unresolved redistribution status.
- `prepare_data.py`: deterministic data preparation only. It does not invoke models or alter the source archive.

## Interpretation and licensing boundaries

The task organizers published this corpus for the NLI4CT benchmark; its annotations are credited to the original dataset authors. This preparation adds no expert labeling. The benchmark has been public for years and may have been in model pretraining; previously unseen evaluation cannot be claimed. A 100-example subset is not a full official leaderboard evaluation. Group related examples by trial when assessing uncertainty; two statements about the same trial are not independent trials.

The inspected official GitHub repository reports no named license, has no LICENSE file, and the ZIP includes no license document. The official download page expressly supplies the benchmark, but the inspected competition terms refer to its licence without naming one. Do not infer a dataset license from the paper's license or from a third-party mirror. This copy is retained for local preparation/evaluation from the official task release; a permission to redistribute or commercially reuse the corpus is not asserted. Do not include the ZIP, input text or gold files in the public manuscript supplement until the upstream dataset license is established; publish preparation code, IDs, hashes and upstream download instructions instead.

Official sources:

- [Fixed repository revision](https://github.com/ai-systems/nli4ct/tree/8e8e5fc506b916984011873403b9ea5ed6dd96c2)
- [Official download page](https://sites.google.com/view/nli4ct/semeval-2023/get-data-and-starting-kit)
- [Official data formatting](https://sites.google.com/view/nli4ct/semeval-2023/data-formatting)
- [Task paper](https://aclanthology.org/2023.semeval-1.307/)
- [Dataset paper](https://aclanthology.org/2023.emnlp-main.1041/)
- [Competition terms](https://codalab.lisn.upsaclay.fr/competitions/8937#learn_the_details-terms_and_conditions)
