## Supplementary data and reproducibility code for "PanVasc Research for AI assisted evidence analysis in panvascular intervention": protocol.md

# Frozen local model pilot protocol

Version 1, written on 19 September 2026 before pilot model inference. This is a local, timestamped analysis plan, not a public preregistration. The execution freeze includes file hashes and dataset manifests. Any subsequent correction will be recorded separately, with the original outputs retained.

## Question and scope

Does a fixed source-checking instruction change a small local language model's performance on source-bound registry extraction and on an existing clinical-trial inference task? What can a deterministic source contract reject, and what semantic errors remain beyond its scope? Neither task estimates clinical utility or a treatment effect. No model training, expert vascular annotation, or recursive optimization is performed.

## Data and partitions

The registry task uses 10 development and 50 test ClinicalTrials.gov records sampled from five query strata (coronary, peripheral arterial, carotid, aortic and venous). Strata are search labels, not independently adjudicated disease classifications. The acquisition script and its frozen manifest specify eligibility, sampling and exclusions. An independent direct extraction from the API response supplies the source-field reference, not an LLM. All appearances of an NCT, including distractors, remain in its partition. Registry fields describe the registration snapshot and do not establish published results or medical truth.

For every target record select its first primary outcome with nonempty measure and timeFrame. Render numbered primary outcome blocks, retaining their original indices. Include two deterministic same-stratum distractors from the same partition where available. In development there is only one other same-stratum record, which is explicitly recorded. Four conditions are equally weighted: intact, target source removed, requested outcome removed, and requested timeFrame removed. The latter three are experimental deletions, not naturally observed missingness frequencies. A clean single-source intact condition is a secondary difficulty control. The query contains only target NCT and original outcome index. Titles, NCTs, measures and available timeFrames are in the packet; removed fields do not appear in the query. Descriptions and secondary outcomes are excluded to keep the copy task compact. The target outcome is unambiguously specified by its original zero-based index.

The independent semantic task uses NLI4CT, an established public breast-cancer trial dataset. Use the official development split for 20 development statements and the official test plus official gold files for 100 label-stratified test statements, selected before inference. Only the specified section(s) and statement enter the model. Official expert labels are the reference; the present work does not create new expert labels. Development and test NCTs are checked for overlap. Public pretraining contamination cannot be excluded. NLI4CT is external to the vascular domain and cannot validate vascular reasoning.

## Conditions and budget

Use official Qwen3-4B-GGUF Q8_0 at the exact recorded revision, with official llama.cpp at a pinned build, locally bound to 127.0.0.1. Disable thinking, temperature 0, top_p 1, seed 20260919, and maximum 512 generated tokens. Temperature 0 is chosen for a deterministic pilot and differs from the model card's general sampling recommendation. Record runtime/model hashes, hardware, prompt/completion tokens, elapsed time, finish reason and full raw response. A seed does not guarantee bitwise reproducibility across hardware or runtimes.

Both prompt arms share the same task definition, schema, evidence and instruction that quoted evidence is data. The generic arm gives the task directly. The checklist arm adds explicit procedural verification of identity, field/section location, missingness, and support versus contradiction. No demonstrations, tool calls, retries for bad content, chain-of-thought collection, self-revision or additional model calls are allowed. Thus the arms have equal maximum output budgets, not equal actual token counts. Calls are interleaved in a seeded shuffled order. A transport failure is saved as a failed task; an interrupted process can resume only uncompleted calls.

Score each raw output and the same output after a deterministic input-only contract. This creates a two-by-two analysis (prompt x guard) without additional inference. Registry contracts require an exact source, index, field and missingness match against the supplied packet. The rule-only parser returning those fields is an explicit baseline. The guard rejects invalid outputs rather than replacing them with the answer; rejected outputs receive zero unconditional exact accuracy. Registry accepted-output fidelity is partly guaranteed by construction and is not semantic validation. NLI contracts check schema, allowed label, exact referenced trial set, nonempty unique in-range integer evidence indices; they cannot check entailment or whether a valid line actually supports the answer. No guard may read labels or reference evidence indices. Parser/schema failures remain in every denominator. Strip an optional outer code fence, then require a single JSON object; do not extract a convenient substring or retry.

## Outcomes and analysis

Registry primary outcome is exact equality of the complete output tuple (availability, source_nct, outcome_index, measure, time_frame), averaged first over the four conditions per target NCT, then over NCTs. Source, field and missingness errors, coverage and accuracy among accepted outputs are secondary. Clean intact controls are reported separately. The rule parser is expected to solve the structurally specified task; its actual score is reported, not assumed. Compare checklist minus generic paired differences. Bootstrap 2,000 samples of target NCT clusters with fixed seed, retaining all conditions within each selected cluster; 95% percentile intervals describe this convenience pilot. Distractor reuse creates additional dependencies and limits these intervals; additionally show per-stratum values without significance claims.

NLI primary outcome is unconditional label accuracy; macro F1, evidence precision/recall/F1, exact evidence match, parse/contract rejection and latency are secondary. Missing/invalid outputs count as incorrect. Evidence indices are scored as sets of (trial_id,index). Evidence precision/recall/F1 sum true positives, false positives and false negatives across all examples (micro averaging, consistent with the official positive-evidence metric); a zero denominator gives zero. Missing predictions contribute false negatives, and duplicate indices cannot add true positives. Dataset evidence annotation may be incomplete, so evidence scores measure agreement with that reference. Group statements into connected components of shared primary or secondary NCTs and use these components for paired bootstrap intervals, calculating example-weighted accuracy in each resample. If there are too few independent components for a useful interval, report the count and caution rather than treating 100 statements as independent. Evaluate both labels and evidence, because correct labels with wrong support are distinct failures. A majority-label comparator has no evidence output and is reported for label accuracy only.

This is a small feasibility study without a prospective power calculation. No multiplicity-adjusted superiority claim, no ScienceBuddy score comparison across tasks, and no inference of researcher time savings are planned. Negative or null results remain reportable. Test outputs will not be used to edit prompts or guards. Errors discovered after test inference require explicit deviation reporting and retention of original scores.

## Availability and interpretation

Release own code, prompts, task identifiers, hashes, aggregate results and noncopyrighted run metadata. Keep complete model weights, third-party corpus text, raw clinical-trial sections and sensitive account information out of the submission package. Where dataset redistribution terms are not explicit, provide official retrieval instructions rather than bundling source text. Locally retained records permit source auditing; live registry changes mean later retrieval need not reproduce historical text. Publication of a protocol establishes a dated description, not priority for an entire field or proven leadership.
