## Supplementary data and reproducibility code for "PanVasc Research for AI assisted evidence analysis in panvascular intervention": verify_pilot.py

````
"""Independent software preflight; never connects to the inference server."""
from __future__ import annotations
import collections
import contextlib
import copy
from datetime import datetime, timezone
import hashlib
import importlib.util
import io
import json
from pathlib import Path
import tempfile
import unittest
from unittest.mock import patch

HERE=Path(__file__).resolve().parent
ROOT=HERE.parents[1]
spec=importlib.util.spec_from_file_location('pilot_under_review',HERE/'run_pilot.py')
pilot=importlib.util.module_from_spec(spec)
spec.loader.exec_module(pilot)

def readj(path): return json.loads(Path(path).read_text(encoding='utf-8-sig'))
def rows(path): return [json.loads(x) for x in Path(path).read_text(encoding='utf-8-sig').splitlines() if x.strip()]
def reg_task():
    return {'id':'registry-fixture','benchmark':'registry','cluster':'NCT00000001','domain':'coronary','condition':'intact',
            'query':{'target_nct':'NCT00000001','outcome_index':0},
            'packet':[{'nct_id':'NCT00000001','title':'Fixture only','primary_outcomes':[
                {'index':0,'measure':'Fixture outcome','time_frame':'30 days'},
                {'index':3,'measure':'Other outcome','time_frame':'90 days'}]}]}
def reg_answer():
    return {'availability':'complete','source_nct':'NCT00000001','outcome_index':0,'measure':'Fixture outcome','time_frame':'30 days'}
def nli_task(case='nli-1',nct='NCT00000011'):
    return {'id':case,'benchmark':'nli4ct','statement':'Fixture statement','section':'Results',
            'sources':[{'role':'primary','trial_id':nct,'section':'Results','lines':[
                {'index':0,'text':'Fixture context'},{'index':1,'text':'Fixture answer line'}]}]}

class SoftwarePreflight(unittest.TestCase):
    def test_parser_single_object_and_fence(self):
        obj={'label':'Entailment','evidence':[]}
        self.assertEqual(pilot.parse_output('```json\n'+json.dumps(obj)+'\n```'),(obj,None))
        for text in ['[]','null','before '+json.dumps(obj),json.dumps(obj)+' trailing',json.dumps(obj)+json.dumps(obj)]:
            self.assertIsNone(pilot.parse_output(text)[0])

    def test_registry_exact_types(self):
        task=reg_task(); answer=reg_answer()
        self.assertTrue(pilot.schema_valid(task,answer))
        self.assertTrue(pilot.contract(task,answer)[0])
        for invalid in [False,0.0,'0']:
            wrong={**answer,'outcome_index':invalid}
            self.assertFalse(pilot.schema_valid(task,wrong))
            self.assertFalse(pilot.contract(task,wrong)[0])

    def test_registry_deletions_do_not_infer(self):
        original=reg_task(); task=copy.deepcopy(original)
        task['packet']=[]
        self.assertEqual(pilot.reference_from_packet(task),{'availability':'missing_source','source_nct':None,'outcome_index':None,'measure':None,'time_frame':None})
        task=copy.deepcopy(original); task['packet'][0]['primary_outcomes'].pop(0)
        self.assertEqual(pilot.reference_from_packet(task),{'availability':'missing_endpoint','source_nct':'NCT00000001','outcome_index':None,'measure':None,'time_frame':None})
        task=copy.deepcopy(original); task['packet'][0]['primary_outcomes'][0].pop('time_frame')
        task['packet'][0]['title']='A title suggesting 30 days must not fill the missing field'
        self.assertEqual(pilot.reference_from_packet(task),{'availability':'missing_time','source_nct':'NCT00000001','outcome_index':0,'measure':'Fixture outcome','time_frame':None})
        self.assertFalse(pilot.contract(task,reg_answer())[0])
        self.assertEqual(original,reg_task())

    def test_nli_schema_and_input_only_guard(self):
        task=nli_task(); answer={'label':'Entailment','evidence':[{'trial_id':'NCT00000011','indices':[0]}]}
        # Line zero is only context in this fixture; the guard intentionally cannot assess support.
        self.assertTrue(pilot.contract(task,answer)[0])
        self.assertFalse(pilot.schema_valid(task,{'label':'Entailment'}))
        for indices in [[0,0],[2],[-1],[True],['0'],[]]:
            invalid={'label':'Entailment','evidence':[{'trial_id':'NCT00000011','indices':indices}]}
            self.assertFalse(pilot.contract(task,invalid)[0])
        self.assertFalse(pilot.contract(task,{'label':'Contradiction','evidence':[{'trial_id':'NCT99999999','indices':[0]}]})[0])

    def test_nct_connections_and_row_weighted_estimate(self):
        a=nli_task('a','NCT1');b=nli_task('b','NCT2');b['sources'].append(copy.deepcopy(a['sources'][0]))
        c=nli_task('c','NCT3');d=nli_task('d','NCT3')
        groups=pilot.components([a,b,c,d])
        self.assertEqual(groups['a'],groups['b']);self.assertEqual(groups['c'],groups['d'])
        self.assertNotEqual(groups['a'],groups['c'])
        mixed={'one':[1],'three':[0,0,0]}
        self.assertEqual(pilot.interval(mixed,row_weighted=True)['estimate'],0.25)
        self.assertEqual(pilot.interval(mixed,row_weighted=False)['estimate'],0.5)

    def test_metrics_count_invalids_and_evidence_per_example(self):
        tasks=[reg_task(),nli_task('nli-1','NCT00000011'),nli_task('nli-2','NCT00000011')]
        truth=[{'id':'registry-fixture','answer':reg_answer()},
               {'id':'nli-1','label':'Entailment','evidence':[['NCT00000011',0]]},
               {'id':'nli-2','label':'Contradiction','evidence':[['NCT00000011',1]]}]
        with tempfile.TemporaryDirectory(prefix='panvasc_preflight_') as folder:
            root=Path(folder);(root/'data').mkdir();(root/'runs').mkdir()
            pilot.save_rows(root/'data/test_inputs.jsonl',tasks)
            pilot.save_rows(root/'data/test_expected.jsonl',truth)
            pilot.save(root/'execution_freeze.json',{'fixture_only':True})
            freeze_hash=pilot.sha(root/'execution_freeze.json');outputs=[]
            for arm in pilot.ARMS:
                for task in tasks:
                    if task['benchmark']=='registry':content={**reg_answer(),'outcome_index':False} if arm=='generic' else reg_answer()
                    elif task['id']=='nli-1':content={'label':'Entailment'}  # schema-invalid, must be wrong
                    else:content={'label':'Contradiction','evidence':[{'trial_id':'NCT00000011','indices':[0,1]}]}
                    outputs.append({'id':task['id'],'arm':arm,'transport_ok':True,'elapsed_seconds':0.01,
                        'execution_freeze_sha256':freeze_hash,
                        'response':{'choices':[{'message':{'content':json.dumps(content)},'finish_reason':'stop'}],
                            'usage':{'completion_tokens':10,'prompt_tokens':30}}})
            pilot.save_rows(root/'runs/test_responses.jsonl',outputs)
            with patch.object(pilot,'HERE',root),contextlib.redirect_stdout(io.StringIO()):pilot.evaluate('test')
            result=readj(root/'runs/test_summary.json')
            self.assertEqual(result['arms']['registry/generic/raw']['correct'],0)
            self.assertEqual(result['arms']['registry/checklist/raw']['correct'],1)
            nli=result['arms']['nli4ct/generic/raw']
            self.assertEqual((nli['n'],nli['correct']),(2,1))
            self.assertEqual((nli['evidence']['tp'],nli['evidence']['fp'],nli['evidence']['fn']),(1,1,1))
            self.assertEqual(nli['evidence']['f1'],0.5)
            self.assertEqual(nli['cluster_accuracy']['estimate'],0.5)
            # A missing arm cannot be silently dropped from a denominator.
            pilot.save_rows(root/'runs/test_responses.jsonl',outputs[:-1])
            with patch.object(pilot,'HERE',root),self.assertRaises(AssertionError):pilot.evaluate('test')

    def test_actual_partitions_and_unmodified_nli_reference(self):
        tasks={s:rows(HERE/'data'/f'{s}_inputs.jsonl') for s in ['development','test']}
        self.assertEqual(collections.Counter(t['benchmark'] for t in tasks['development']),{'registry':50,'nli4ct':20})
        self.assertEqual(collections.Counter(t['benchmark'] for t in tasks['test']),{'registry':250,'nli4ct':100})
        for benchmark in ['registry','nli4ct']:
            seen={}
            for split,values in tasks.items():
                subset=[t for t in values if t['benchmark']==benchmark]
                seen[split]={r['nct_id'] for t in subset for r in t['packet']} if benchmark=='registry' else {s['trial_id'] for t in subset for s in t['sources']}
                self.assertEqual(len({t['id'] for t in subset}),len(subset))
            self.assertFalse(seen['development']&seen['test'])
        for split,tag in [('development','dev'),('test','test')]:
            expected={r['id']:r for r in rows(HERE/'data'/f'{split}_expected.jsonl')}
            reference={r['id']:r for r in rows(ROOT/'experiments/nli4ct/reference'/f'{tag}_expected.jsonl')}
            source_input={r['id']:r for r in rows(ROOT/'experiments/nli4ct'/f'{tag}_inputs.jsonl')}
            for t in tasks[split]:
                if t['benchmark']!='nli4ct':continue
                original=source_input[t['id']];gold=reference[t['id']]
                for key in ['statement','sources','section','type']:self.assertEqual(t[key],original[key])
                self.assertNotIn('label',t);self.assertNotIn('evidence_indices',t)
                self.assertEqual(expected[t['id']]['label'],gold['label'])
                pairs={(gold['source_trial_ids'][role],i) for role,ii in gold['evidence_indices'].items() for i in ii}
                self.assertEqual({tuple(x) for x in expected[t['id']]['evidence']},pairs)

    def test_actual_registry_perturbations_against_source_truth(self):
        source=Path(readj(ROOT/'experiments/registry_grounding/latest_source_run.json')['path'])
        original={(r['nct_id'],r['outcome_index']):r['source_fields'] for r in rows(source/'source_truth.jsonl') if r['outcome_type']=='PRIMARY'}
        for split in ['development','test']:
            expected={r['id']:r for r in rows(HERE/'data'/f'{split}_expected.jsonl')}
            registry=[t for t in rows(HERE/'data'/f'{split}_inputs.jsonl') if t['benchmark']=='registry']
            bytarget=collections.defaultdict(list)
            for t in registry:
                q=t['query'];self.assertEqual(set(q),{'target_nct','outcome_index'})
                bytarget[q['target_nct']].append(t)
                fields=original[(q['target_nct'],q['outcome_index'])]
                answer={'availability':'complete','source_nct':q['target_nct'],'outcome_index':q['outcome_index'],'measure':fields['measure'],'time_frame':fields['timeFrame']}
                if t['condition']=='missing_source':answer={'availability':'missing_source','source_nct':None,'outcome_index':None,'measure':None,'time_frame':None}
                elif t['condition']=='missing_endpoint':answer={'availability':'missing_endpoint','source_nct':q['target_nct'],'outcome_index':None,'measure':None,'time_frame':None}
                elif t['condition']=='missing_time':answer.update(availability='missing_time',time_frame=None)
                self.assertEqual(expected[t['id']]['answer'],answer)
                self.assertEqual(pilot.reference_from_packet(t),answer)
                if t['condition']=='clean':self.assertEqual(len(t['packet']),1)
            for group in bytarget.values():
                self.assertEqual({t['condition'] for t in group},{'intact','clean','missing_source','missing_endpoint','missing_time'})

if __name__=='__main__':
    # Block accidental network/inference calls by the imported implementation.
    suite=unittest.defaultTestLoader.loadTestsFromTestCase(SoftwarePreflight)
    buffer=io.StringIO()
    with patch.object(pilot.urllib.request,'build_opener',side_effect=AssertionError('Network forbidden in preflight')):
        result=unittest.TextTestRunner(stream=buffer,verbosity=2).run(suite)
    report={'checked_at_utc':datetime.now(timezone.utc).isoformat(),
        'status':'passed' if result.wasSuccessful() else 'failed','tests':result.testsRun,
        'failures':len(result.failures),'errors':len(result.errors),
        'run_pilot_sha256':hashlib.sha256((HERE/'run_pilot.py').read_bytes()).hexdigest(),
        'verify_pilot_sha256':hashlib.sha256(Path(__file__).read_bytes()).hexdigest(),
        'scope':'Independent software fixtures and frozen data integrity; no model inference, no scientific performance results.',
        'details':buffer.getvalue()}
    (HERE/'preflight_verification.json').write_text(json.dumps(report,ensure_ascii=False,indent=2)+'\n',encoding='utf-8')
    print(buffer.getvalue());print(json.dumps({k:v for k,v in report.items() if k!='details'},indent=2))
    raise SystemExit(0 if result.wasSuccessful() else 1)
````
