## Supplementary data and reproducibility code for "PanVasc Research for AI assisted evidence analysis in panvascular intervention": prepare_records.py

```
"""Acquire a bounded, query-stratified ClinicalTrials.gov source-grounding set.

Only standard-library dependencies. TLS certificate/hostname verification stays on.
No contacts, locations, eligibility, participant results, or raw responses are saved.
The independent source_truth file is copied directly from structured source fields,
not generated by a model and not an expert clinical-semantic reference standard.
"""
from __future__ import annotations

import argparse
from collections import Counter
from datetime import datetime, timezone
import hashlib
import json
from pathlib import Path
import random
import ssl
import time
import urllib.error
import urllib.parse
import urllib.request

BASE_URL = "https://clinicaltrials.gov/api/v2/studies"
SEED = 20260919
EXCLUDED_PILOT_NCTS = {"NCT00589927", "NCT02004483", "NCT00790335"}
QUERIES = [
    ("coronary", {"query.cond": "coronary", "query.intr": "stent"}),
    ("peripheral", {"query.cond": "peripheral arterial disease", "query.intr": "endovascular"}),
    ("carotid", {"query.cond": "carotid stenosis", "query.intr": "stent"}),
    ("aortic", {"query.cond": "aortic aneurysm", "query.intr": "endovascular"}),
    ("venous", {"query.cond": "venous thrombosis", "query.intr": "stent OR thrombectomy OR catheter"}),
]
OUTCOME_FIELDS = ("measure", "description", "timeFrame")
RETRY_HTTP = {408, 429, 500, 502, 503, 504}


def utc_now():
    return datetime.now(timezone.utc).isoformat()


def digest(data: bytes):
    return hashlib.sha256(data).hexdigest()


def canonical_bytes(obj):
    return json.dumps(obj, sort_keys=True, ensure_ascii=False, separators=(",", ":")).encode("utf-8")


def write_json(path, obj):
    path.write_text(json.dumps(obj, ensure_ascii=False, indent=2) + "\n", encoding="utf-8")


def write_jsonl(path, rows):
    path.write_text("".join(json.dumps(row, ensure_ascii=False, sort_keys=True) + "\n" for row in rows), encoding="utf-8")


def field_state(outcome, field):
    if field not in outcome:
        return "absent"
    value = outcome[field]
    if value is None:
        return "null"
    if isinstance(value, str) and not value.strip():
        return "empty"
    return "reported"


def source_truth_from_response(study):
    """Read truth independently from the transient original API object."""
    section = study.get("protocolSection", {})
    nct = section.get("identificationModule", {}).get("nctId")
    outcomes = section.get("outcomesModule", {})
    rows = []
    for kind, key in (("PRIMARY", "primaryOutcomes"), ("SECONDARY", "secondaryOutcomes")):
        for index, outcome in enumerate(outcomes.get(key, [])):
            values = {field: outcome.get(field) for field in OUTCOME_FIELDS}
            rows.append({
                "nct_id": nct,
                "outcome_type": kind,
                "outcome_index": index,
                "source_id": nct,
                "source_url": f"https://clinicaltrials.gov/study/{nct}",
                "source_object_path": f"protocolSection.outcomesModule.{key}[{index}]",
                "source_fields": values,
                "source_field_states": {field: field_state(outcome, field) for field in OUTCOME_FIELDS},
                "source_field_sha256": {field: digest(canonical_bytes(outcome.get(field))) for field in OUTCOME_FIELDS},
                "source_outcome_sha256": digest(canonical_bytes(values)),
                "reference_type": "registry_structured_source_truth_not_expert_gold",
                "controlled_perturbations_applied": False,
            })
    return rows


def sanitize(study, query_domain, query_rank):
    section = study.get("protocolSection", {})
    identification = section.get("identificationModule", {})
    arms = section.get("armsInterventionsModule", {})
    outcomes = section.get("outcomesModule", {})
    nct = identification.get("nctId")
    core = {
        "nct_id": nct,
        "brief_title": identification.get("briefTitle"),
        "official_title": identification.get("officialTitle"),
        "study_type": section.get("designModule", {}).get("studyType"),
        "conditions": section.get("conditionsModule", {}).get("conditions", []),
        "interventions": [
            {key: intervention[key] for key in ("type", "name", "description", "otherNames") if key in intervention}
            for intervention in arms.get("interventions", [])
        ],
        "primary_outcomes": [{field: outcome[field] for field in OUTCOME_FIELDS if field in outcome}
                             for outcome in outcomes.get("primaryOutcomes", [])],
        "secondary_outcomes": [{field: outcome[field] for field in OUTCOME_FIELDS if field in outcome}
                               for outcome in outcomes.get("secondaryOutcomes", [])],
    }
    valid = [index for index, outcome in enumerate(core["primary_outcomes"])
             if isinstance(outcome.get("measure"), str) and outcome["measure"].strip()
             and isinstance(outcome.get("timeFrame"), str) and outcome["timeFrame"].strip()]
    return {
        **core,
        "source_url": f"https://clinicaltrials.gov/study/{nct}",
        "source_system": "ClinicalTrials.gov API v2",
        "first_query_domain": query_domain,
        "query_memberships": [{"domain": query_domain, "rank": query_rank}],
        "eligible_primary_indices": valid,
        "registry_fields_sha256": digest(canonical_bytes(core)),
        "domain_assignment_type": "query_stratum_not_expert_clinical_label",
    }


def eligibility_reasons(record):
    reasons = []
    if not record["nct_id"]:
        reasons.append("missing_nct_id")
    if record["nct_id"] in EXCLUDED_PILOT_NCTS:
        reasons.append("previously_viewed_pilot")
    if record["study_type"] != "INTERVENTIONAL":
        reasons.append("not_interventional")
    if not any(intervention.get("type") in {"DEVICE", "PROCEDURE"} for intervention in record["interventions"]):
        reasons.append("no_device_or_procedure")
    if not record["eligible_primary_indices"]:
        reasons.append("no_primary_with_nonempty_measure_and_timeframe")
    return reasons


def fetch_query(domain, query, *, timeout, max_attempts, context, output_dir, query_log):
    params = {**query, "pageSize": 100, "format": "json", "countTotal": "true"}
    url = BASE_URL + "?" + urllib.parse.urlencode(params)
    entry = {"domain": domain, "parameters": params, "url": url,
             "one_page_only": True, "ordering": "API default; no explicit sort",
             "tls_verification": True, "attempts": [], "status": "pending"}
    query_log.append(entry)
    write_json(output_dir / "query_log.json", query_log)
    for attempt in range(1, max_attempts + 1):
        attempt_log = {"attempt": attempt, "started_utc": utc_now()}
        started = time.perf_counter()
        body = None
        try:
            request = urllib.request.Request(url, headers={
                "Accept": "application/json",
                "User-Agent": "PanVascResearch-registry-grounding/0.1 (bounded research metadata retrieval)",
            })
            with urllib.request.urlopen(request, timeout=timeout, context=context) as response:
                body = response.read()
                attempt_log["http_status"] = response.status
                attempt_log["final_url"] = response.url
            attempt_log["response_sha256"] = digest(body)
            attempt_log["response_bytes"] = len(body)
            payload = json.loads(body)
            studies = payload.get("studies")
            if not isinstance(studies, list):
                raise ValueError("JSON response lacks a studies list")
            if len(studies) > 100:
                raise ValueError("Response exceeds the predetermined 100-record page limit")
            attempt_log["success"] = True
            entry.update(status="success", total_count=payload.get("totalCount"),
                         returned_count=len(studies), has_next_page=bool(payload.get("nextPageToken")))
            entry["returned_nct_ids"] = [study.get("protocolSection", {}).get("identificationModule", {}).get("nctId")
                                         for study in studies]
            return studies
        except urllib.error.HTTPError as error:
            attempt_log.update(success=False, error_type=type(error).__name__, http_status=error.code,
                               error_message=str(error))
            body = error.read()
            attempt_log["response_sha256"] = digest(body)
            attempt_log["response_bytes"] = len(body)
            if error.code not in RETRY_HTTP:
                break
        except (urllib.error.URLError, TimeoutError, OSError, ValueError, json.JSONDecodeError) as error:
            attempt_log.update(success=False, error_type=type(error).__name__, error_message=str(error))
        finally:
            if body is not None and "response_sha256" not in attempt_log:
                attempt_log["response_sha256"] = digest(body)
                attempt_log["response_bytes"] = len(body)
            attempt_log["elapsed_seconds"] = round(time.perf_counter() - started, 6)
            attempt_log["ended_utc"] = utc_now()
            entry["attempts"].append(attempt_log)
            write_json(output_dir / "query_log.json", query_log)
        if attempt < max_attempts:
            time.sleep(min(2 ** (attempt - 1), 4))
    entry["status"] = "failed"
    entry["returned_count"] = 0
    write_json(output_dir / "query_log.json", query_log)
    return []


def validate_source_truth(selected, truths):
    by_id = {row["nct_id"]: row for row in selected}
    for truth in truths:
        source_key = "primary_outcomes" if truth["outcome_type"] == "PRIMARY" else "secondary_outcomes"
        outcome = by_id[truth["nct_id"]][source_key][truth["outcome_index"]]
        for field in OUTCOME_FIELDS:
            if outcome.get(field) != truth["source_fields"][field]:
                raise AssertionError("Sanitized record disagrees with independent source copy")
    if len(by_id) != len(selected):
        raise AssertionError("Selected NCT appears more than once")
    if set(by_id) & EXCLUDED_PILOT_NCTS:
        raise AssertionError("Previously viewed pilot record leaked into frozen split")


def main():
    parser = argparse.ArgumentParser(description=__doc__)
    parser.add_argument("--output-root", type=Path, default=Path(__file__).resolve().parent / "data")
    parser.add_argument("--timeout", type=float, default=30)
    parser.add_argument("--attempts", type=int, default=3, choices=(1, 2, 3))
    args = parser.parse_args()
    run_id = datetime.now(timezone.utc).strftime("%Y%m%dT%H%M%SZ")
    output_dir = args.output_root / run_id
    output_dir.mkdir(parents=True, exist_ok=False)
    query_log, candidates, truths, selection_log = [], {}, {}, []
    started_utc = utc_now()
    context = ssl.create_default_context()
    if not context.check_hostname or context.verify_mode != ssl.CERT_REQUIRED:
        raise AssertionError("TLS verification must be enabled")

    for domain, query in QUERIES:
        studies = fetch_query(domain, query, timeout=args.timeout, max_attempts=args.attempts,
                              context=context, output_dir=output_dir, query_log=query_log)
        for rank, study in enumerate(studies, start=1):
            record = sanitize(study, domain, rank)
            nct = record["nct_id"]
            if not nct:
                selection_log.append({"domain": domain, "rank": rank, "reason": "missing_nct_id"})
                continue
            if nct in candidates:
                candidates[nct]["query_memberships"].append({"domain": domain, "rank": rank})
                selection_log.append({"nct_id": nct, "domain": domain, "reason": "duplicate_assigned_to_first_query_domain",
                                      "assigned_domain": candidates[nct]["first_query_domain"],
                                      "snapshot_fields_match": candidates[nct]["registry_fields_sha256"] == record["registry_fields_sha256"]})
                continue
            candidates[nct] = record
            truths[nct] = source_truth_from_response(study)
        # Only sanitized data are persisted. Original response bodies remain transient.
        write_jsonl(output_dir / "candidates.jsonl", list(candidates.values()))
        print(json.dumps({"domain": domain, "returned": len(studies), "unique_candidates_so_far": len(candidates)}, ensure_ascii=False), flush=True)

    development, test = [], []
    counts = {}
    for domain_index, (domain, _) in enumerate(QUERIES):
        domain_candidates = [row for row in candidates.values() if row["first_query_domain"] == domain]
        eligible = []
        for record in domain_candidates:
            reasons = eligibility_reasons(record)
            record["eligibility_status"] = "eligible" if not reasons else "ineligible"
            record["eligibility_reasons"] = reasons
            if not reasons:
                eligible.append(record)
        eligible.sort(key=lambda row: row["nct_id"])
        # Separate deterministic streams make other-domain request failures unable
        # to change selection within this domain. Seed and index are frozen.
        random.Random(SEED + domain_index).shuffle(eligible)
        dev_rows, test_rows = eligible[:2], eligible[2:12]
        for split, rows in (("development", dev_rows), ("test", test_rows)):
            for row in rows:
                selected = {**row, "split": split, "domain": domain, "retrieval_run_id": run_id}
                (development if split == "development" else test).append(selected)
                selection_log.append({"nct_id": row["nct_id"], "domain": domain, "split": split,
                                      "selection": "frozen_seeded_sample_before_model_inference"})
        counts[domain] = {"assigned_unique_candidates": len(domain_candidates), "eligible": len(eligible),
                          "development": len(dev_rows), "test": len(test_rows),
                          "requested_development": 2, "requested_test": 10,
                          "shortfall": max(0, 12 - len(eligible)), "rng_seed": SEED + domain_index}

    selected = development + test
    truth_rows = []
    for record in selected:
        for truth in truths[record["nct_id"]]:
            truth_rows.append({**truth, "domain": record["domain"], "split": record["split"],
                               "retrieval_run_id": run_id,
                               "registry_fields_sha256": record["registry_fields_sha256"]})
    validate_source_truth(selected, truth_rows)
    write_jsonl(output_dir / "candidates.jsonl", list(candidates.values()))
    write_jsonl(output_dir / "development_records.jsonl", development)
    write_jsonl(output_dir / "test_records.jsonl", test)
    write_jsonl(output_dir / "source_truth.jsonl", truth_rows)
    write_json(output_dir / "selection_log.json", selection_log)
    summary = {
        "run_id": run_id, "started_utc": started_utc, "frozen_utc": utc_now(), "counts_by_domain": counts,
        "unique_candidates": len(candidates), "development_records": len(development), "test_records": len(test),
        "source_truth_rows": len(truth_rows), "source_truth_rows_by_kind": dict(Counter(row["outcome_type"] for row in truth_rows)),
        "successful_queries": sum(row["status"] == "success" for row in query_log),
        "total_attempts": sum(len(row["attempts"]) for row in query_log),
        "failed_attempts": sum(not attempt["success"] for row in query_log for attempt in row["attempts"]),
        "nct_overlap_between_splits": 0, "duplicate_selected_ncts": 0,
        "excluded_previously_viewed_ncts": sorted(EXCLUDED_PILOT_NCTS),
        "model_inference_performed_by_preparation": False, "expert_semantic_annotations": 0,
        "raw_responses_retained": False, "contacts_or_locations_retained": False,
        "reference_scope": "source-field fidelity, not medical validity or treatment efficacy",
    }
    write_json(output_dir / "summary.json", summary)
    hashed_names = ["candidates.jsonl", "development_records.jsonl", "test_records.jsonl", "source_truth.jsonl",
                    "query_log.json", "selection_log.json", "summary.json"]
    manifest = {
        "schema_version": "registry-grounding-source-v1", "run_id": run_id,
        "frozen_utc": summary["frozen_utc"], "random_seed": SEED,
        "domain_order": [domain for domain, _ in QUERIES],
        "selection_algorithm": "first-query domain assignment; eligible NCT ascending; Random(seed + domain_index).shuffle; first 2 development, next 10 test",
        "eligibility": ["studyType == INTERVENTIONAL", "at least one intervention type DEVICE or PROCEDURE",
                        "at least one primary outcome with nonempty string measure and timeFrame", "not a previously viewed pilot NCT"],
        "excluded_previously_viewed_ncts": sorted(EXCLUDED_PILOT_NCTS),
        "counts_by_domain": counts,
        "split_ids": {"development": [row["nct_id"] for row in development], "test": [row["nct_id"] for row in test]},
        "files_sha256": {name: digest((output_dir / name).read_bytes()) for name in hashed_names},
        "preparation_script_sha256": digest(Path(__file__).read_bytes()),
        "terms_url": "https://clinicaltrials.gov/about-site/terms-conditions",
        "api_documentation_url": "https://clinicaltrials.gov/data-api/api",
        "field_documentation_url": "https://clinicaltrials.gov/data-api/about-api/study-data-structure",
        "domain_labels_are_query_strata": True,
        "registry_text_public_redistribution_planned": False,
        "public_release_plan": "script, NCT identifiers, field hashes and experiment outputs; no long registry text in manuscript supplement",
        "not_systematic_review": True, "not_expert_gold_standard": True,
        "controlled_missingness_not_applied_here": True,
    }
    write_json(output_dir / "frozen_manifest.json", manifest)
    write_json(Path(__file__).resolve().parent / "latest_source_run.json", {
        "run_id": run_id, "path": str(output_dir.resolve()),
        "manifest_sha256": digest((output_dir / "frozen_manifest.json").read_bytes()),
        "summary": summary,
    })
    print(json.dumps(summary, ensure_ascii=False, indent=2), flush=True)


if __name__ == "__main__":
    main()
```
