## Supplementary data and reproducibility code for "PanVasc Research for AI assisted evidence analysis in panvascular intervention": README.md

# Registry source-grounding preparation

This directory contains a bounded ClinicalTrials.gov API v2 preparation run for a source-field fidelity pilot. It is **not** a systematic review, an expert clinical gold standard, an endpoint harmonization validation, or evidence of treatment efficacy. Domain names are query strata, not manually certified medical classifications. Preparation performed no model inference.

## Completed source run

Frozen run: `data/20260919T133142Z/`; acquired 2026-09-19 13:31:42–13:31:54 UTC. `latest_source_run.json` records its absolute path and manifest hash. Each query requested one page, at most 100 records, with the API's default ordering. No pagination, adaptive query replacement, or model-result-based selection occurred. All five requests succeeded on their first attempt; TLS certificate and hostname verification remained enabled.

| Query stratum | Condition query | Intervention query | Returned | Unique assigned | Eligible | Development | Test |
|---|---|---|---:|---:|---:|---:|---:|
| coronary | coronary | stent | 100 | 100 | 57 | 2 | 10 |
| peripheral | peripheral arterial disease | endovascular | 100 | 100 | 53 | 2 | 10 |
| carotid | carotid stenosis | stent | 100 | 100 | 54 | 2 | 10 |
| aortic | aortic aneurysm | endovascular | 100 | 99 | 52 | 2 | 10 |
| venous | venous thrombosis | stent OR thrombectomy OR catheter | 100 | 99 | 30 | 2 | 10 |

There are 498 unique candidates and 60 selected records: 10 development and 50 test. NCT identifiers do not repeat across selected domains or splits. Candidate-level publication/registry relationships beyond identical NCT identifiers have not been adjudicated. API total hits exceed the retained first page in every domain; these are convenience frames, not complete domain inventories.

Eligibility was mechanical and specified before model inference: `studyType == INTERVENTIONAL`; at least one `DEVICE` or `PROCEDURE` intervention; at least one primary outcome with nonempty string `measure` and `timeFrame`; exclusion of previously viewed pilot records. Duplicate NCTs were assigned to the first query domain in the fixed order above, while all query memberships were retained.

Sampling used base seed `20260919`. Within each domain, eligible records were sorted by NCT ID, shuffled using `random.Random(base_seed + zero_based_domain_index)`, and assigned the first 2 to development and the next 10 to test. The exact derived seeds and selected IDs are in the manifest. Separate deterministic streams prevent a failed query in another domain from changing this domain's sample.

## Development exposure

`NCT00589927` and `NCT02004483` had been viewed as API pilot examples and were excluded from selection. After acquisition, the prior novelty review's exposure to ATTRACT (`NCT00790335`) was explicitly documented. ATTRACT is absent from all 498 candidates and both frozen splits. The updated preparation script excludes it on future runs. A replay using all three exclusions reproduced the frozen split exactly; no source record, selected ID, or frozen data hash changed.

`development_exposure_addendum.json` records reasons and the original manifest hash. The run folder's `preparation_script_executed.py` preserves the exact acquisition script corresponding to the manifest's script hash. The root `prepare_records.py` additionally contains the ATTRACT exclusion. A TAMBE original paper was discussed during the novelty review, but its specific NCT identifier had not been viewed or established in that review; no guessed ID was added.

## Files and schemas

| File | Content |
|---|---|
| `prepare_records.py` | Standard-library acquisition/sanitization/filtering/sampling script; future runs write new timestamped folders |
| `data/<run>/candidates.jsonl` | Sanitized unique candidates, query memberships, mechanical eligibility and reasons |
| `data/<run>/development_records.jsonl` | 10 complete sanitized development records |
| `data/<run>/test_records.jsonl` | 50 complete sanitized test records |
| `data/<run>/source_truth.jsonl` | Independent copies of original structured outcome fields: 464 rows, 96 PRIMARY and 368 SECONDARY |
| `data/<run>/frozen_manifest.json` | Split IDs, exact rules, seed, seven file hashes, executed script hash and limitations |
| `data/<run>/query_log.json` | Exact query URLs/parameters, timestamps, API total/returned counts, every attempt, response hashes and status |
| `data/<run>/selection_log.json` | Duplicate assignment and actual selected split assignments |
| `data/<run>/summary.json` | Actual counts and explicit non-evaluation statements |
| `public_source_index.jsonl` | IDs, source paths and field hashes for the 464 outcome rows; no outcome names/descriptions/timeframes |

Each selected record contains `nct_id`, `domain`, `split`, `brief_title`, `official_title`, `study_type`, `conditions`, `interventions`, `primary_outcomes`, `secondary_outcomes`, `source_url`, `eligible_primary_indices`, `registry_fields_sha256`, and acquisition provenance. Original primary and secondary outcome order is preserved. Outcome objects retain exactly the source-provided `measure`, `description`, and `timeFrame` keys; absent keys are not invented. Intervention fields are limited to type, name, description and other names. No contacts, locations, eligibility criteria, participant result rows, or raw API response bodies are retained.

Each `source_truth.jsonl` row contains an `outcome_type`, zero-based `outcome_index`, `source_fields` with the three original field values, their reported/absent/null/empty states, official source URL, JSON source path, per-field hashes and an outcome-object hash. This reference file was copied directly from transient source API objects, separately from the sanitized record path, and then checked for exact agreement. It is an automatic **source-field reference**, not an expert interpretation of the endpoint. The reference file includes all primary/secondary outcomes for selected records; the experiment runner can select `eligible_primary_indices[0]` without making semantic judgments.

Seven frozen file hashes were recomputed successfully. The 60 selected NCT IDs are unique, development and test IDs are disjoint, all selected records contain at least one qualifying primary endpoint, and the independent source copies match the sanitized fields. These are data-integrity checks, not model-performance results.

## Intended controlled task

The separate runner prepares evidence packets containing records from the same query stratum and asks for a specified NCT's primary endpoint name, timeframe and source. Four planned conditions are complete target evidence, absent target source, absent target outcome entry and removed target timeframe. The reference answer must follow what is actually supplied in each packet and preserve missingness with null/abstention. The root runner determines packet construction, prompts, model settings, guard behavior, scoring and controlled perturbation labels.

These deletions are experimentally introduced missingness, not estimates of natural registry missingness. Unchanged source records and independent truth stay intact in this folder. Successful copying of registry fields does not demonstrate that a model understands endpoint definitions, estimands, treatment effects or clinical care.

## Re-running and use of source material

From the project directory:

```powershell
& 'C:/Users/Administrator/.cache/codex-runtimes/codex-primary-runtime/dependencies/python/python.exe' experiments/registry_grounding/prepare_records.py
```

Each execution performs new public API requests and can return a different first page as the registry changes. It does not overwrite the completed frozen run. Default timeout is 30 seconds and there are at most 3 attempts per query; failures are retained, TLS is never disabled, and missing domains are not filled with invented records. Archived raw-body hashes identify response bytes but cannot reconstruct omitted raw bodies.

ClinicalTrials.gov provides the official [API](https://clinicaltrials.gov/data-api/api), [data structure](https://clinicaltrials.gov/data-api/about-api/study-data-structure), [outcome field definitions](https://clinicaltrials.gov/policy/protocol-definitions) and [terms](https://clinicaltrials.gov/about-site/terms-conditions). The terms page did not expose its body in the web retrieval used during this preparation; no universal CC license or unrestricted redistribution authorization is claimed. Registry statements are submitted by study organizations, and matching them is not independent clinical validation. The planned public release consists of acquisition code, NCT IDs, source/field hashes and experiment outputs, with ClinicalTrials.gov attribution; long registry descriptions are kept out of the manuscript supplement. No external model service received these records during preparation.
