## Supplementary data and reproducibility code for "PanVasc Research for AI assisted evidence analysis in panvascular intervention": build_concept_figures.py

```
"""Build two vector scientific figures, with explicit evaluation boundaries.

Uses ReportLab for vector PDF and fitz for PNG rendering. This file does not call
models, change the manuscript, or assert that any model evaluation has finished.
"""
from __future__ import annotations

import math
from pathlib import Path

import fitz
from reportlab.lib.colors import HexColor, white
from reportlab.pdfbase.pdfmetrics import stringWidth
from reportlab.pdfgen import canvas

ROOT = Path(__file__).resolve().parents[1]
OUT = ROOT / "manuscript"
NAVY = HexColor("#19384B")
TEAL = HexColor("#147A80")
INK = HexColor("#243C48")
GRAY = HexColor("#6D7B85")
LIGHT = HexColor("#A7B2B9")
PALE = HexColor("#EEF6F6")
PALE_GRAY = HexColor("#F4F6F7")


class Figure:
    def __init__(self, name, height=462):
        self.path = OUT / f"{name}.pdf"
        self.width, self.height = 720, height
        self.c = canvas.Canvas(str(self.path), pagesize=(self.width, height))
        self.c.setTitle(name.replace("_", " "))
        self.c.setAuthor("PanVasc Research")
        self.c.setSubject("Evaluation design and proposed endpoint representation; no performance claim")
        self.text_boxes = []

    def text(self, x, y, text, *, size=12, bold=False, color=INK, max_width=None, centered=False):
        assert text.isascii(), f"Use ASCII figure text: {text!r}"
        font = "Helvetica-Bold" if bold else "Helvetica"
        width = stringWidth(text, font, size)
        if max_width is not None:
            assert width <= max_width, f"Text wider than panel: {text!r} ({width:.1f}>{max_width})"
        left = x - width / 2 if centered else x
        assert left >= 12 and left + width <= self.width - 12, f"Text crosses page edge: {text}"
        assert y - size * .25 > 6 and y + size < self.height - 6, f"Vertical text bounds: {text}"
        self.c.setFillColor(color)
        self.c.setFont(font, size)
        self.c.drawString(left, y, text)
        self.text_boxes.append((left, y - size * .25, left + width, y + size))

    def box(self, x, y, width, height, *, fill=PALE, stroke=TEAL, dashed=False, radius=7):
        self.c.saveState()
        self.c.setFillColor(fill)
        self.c.setStrokeColor(stroke)
        self.c.setLineWidth(1.25)
        if dashed:
            self.c.setDash(4, 3)
        self.c.roundRect(x, y, width, height, radius, stroke=1, fill=1)
        self.c.restoreState()

    def arrow(self, points, *, color=TEAL, dashed=False, width=1.5, head=5):
        self.c.saveState()
        self.c.setStrokeColor(color)
        self.c.setFillColor(color)
        self.c.setLineWidth(width)
        if dashed:
            self.c.setDash(4, 3)
        path = self.c.beginPath()
        path.moveTo(*points[0])
        for point in points[1:]:
            path.lineTo(*point)
        self.c.drawPath(path)
        self.c.setDash()
        x0, y0 = points[-2]
        x1, y1 = points[-1]
        angle = math.atan2(y1 - y0, x1 - x0)
        p = self.c.beginPath()
        p.moveTo(x1, y1)
        for offset in (.52, -.52):
            p.lineTo(x1 - head * math.cos(angle + offset), y1 - head * math.sin(angle + offset))
        p.close()
        self.c.drawPath(p, fill=1, stroke=0)
        self.c.restoreState()

    def line(self, x1, y1, x2, y2, *, color=LIGHT, dashed=False):
        self.c.saveState()
        self.c.setStrokeColor(color)
        self.c.setLineWidth(1)
        if dashed:
            self.c.setDash(4, 3)
        self.c.line(x1, y1, x2, y2)
        self.c.restoreState()

    def finish(self):
        self.c.showPage()
        self.c.save()
        document = fitz.open(self.path)
        assert len(document) == 1
        page = document[0]
        words = page.get_text("words")
        assert words, "Figure contains no extractable vector text"
        for x0, y0, x1, y1, text, *_ in words:
            assert x0 >= 10 and y0 >= 7 and x1 <= self.width - 10 and y1 <= self.height - 7, text
        png_path = self.path.with_suffix(".png")
        page.get_pixmap(matrix=fitz.Matrix(3, 3), alpha=False).save(png_path)
        document.close()
        print(f"Created {self.path.name} + {png_path.name}: {self.width} x {self.height} pt; {len(words)} words; bounds checked")


def architecture():
    f = Figure("Figure_1_architecture", 465)
    f.text(24, 433, "PanVasc Research: evaluation design", size=21, bold=True, color=NAVY)
    f.text(24, 410, "Source fidelity and external semantic inference are evaluated separately", size=13.2, color=GRAY)
    f.text(360, 383, "Coronary  |  Peripheral  |  Carotid  |  Aortic  |  Venous", size=13.5, color=TEAL, centered=True)

    panels = [
        (24, 150, "Question + sources", ["Specified target", "Fixed source snapshot", "Queries, IDs, hashes"]),
        (195, 159, "Local Qwen3-4B", ["Fixed model weights", "Generic / checklist", "Same supplied evidence"]),
        (375, 151, "Input-only contract", ["Source and field checks", "Preserve missingness", "Accept or abstain"]),
        (547, 149, "Saved outputs", ["Raw + checked answers", "Provenance + failures", "Human review"]),
    ]
    y, height = 267, 98
    for x, width, title, lines in panels:
        f.box(x, y, width, height)
        f.text(x + width / 2, 342, title, size=13.6, bold=True, color=NAVY, max_width=width-12, centered=True)
        for line_index, line in enumerate(lines):
            f.text(x + width / 2, 320 - 18 * line_index, line, size=11.8, max_width=width-14, centered=True)
    for x1, x2 in ((174, 195), (354, 375), (526, 547)):
        f.arrow([(x1, 316), (x2-3, 316)])

    # Both branches describe planned evaluation, not a learned capability ladder.
    f.text(360, 237, "Two complementary pilot tasks", size=13.2, bold=True, color=NAVY, centered=True)
    f.line(360, 231, 360, 221)
    f.line(184, 221, 536, 221)
    f.arrow([(184, 221), (184, 211)], color=GRAY)
    f.arrow([(536, 221), (536, 211)], color=GRAY)
    f.box(24, 109, 320, 100, fill=white)
    f.box(376, 109, 320, 100, fill=white)
    f.text(184, 185, "Registry source fidelity", size=15, bold=True, color=TEAL, centered=True)
    for y, line in ((163, "Endpoint name, timeframe and target source"),
                    (145, "Complete evidence / controlled omissions"),
                    (127, "Reference: structured registry fields")):
        f.text(184, y, line, size=12.4, max_width=296, centered=True)
    f.text(536, 185, "NLI4CT external semantics", size=15, bold=True, color=TEAL, centered=True)
    for y, line in ((163, "Trial statements and entailment labels"),
                    (145, "External breast-cancer trial benchmark"),
                    (127, "Does not validate vascular expertise")):
        f.text(536, y, line, size=12.4, max_width=296, centered=True)

    f.box(24, 54, 672, 38, fill=PALE_GRAY, stroke=LIGHT)
    f.text(38, 68, "Supporting tools", size=12.7, bold=True, color=NAVY)
    f.text(157, 68, "Literature retrieval, source caches and local statistics", size=12.7, color=GRAY)
    f.text(24, 26, "No model training or recursive optimization.", size=11.5, color=GRAY)
    f.text(371, 26, "Design shown; no performance claim.", size=11.5, color=GRAY)
    f.finish()


def endpoint_scope():
    f = Figure("Figure_2_endpoint_scope", 474)
    f.text(24, 442, "Endpoint semantics: proposed research scope", size=20.5, bold=True, color=NAVY)
    f.text(24, 419, "Four-layer representation - proposed and not validated", size=13.5, bold=True, color=GRAY)

    f.box(24, 228, 173, 165, fill=PALE)
    f.text(110.5, 368, "Registry input fields", size=14, bold=True, color=TEAL, centered=True, max_width=151)
    for y, label in ((340, "measure"), (311, "timeFrame"), (282, "NCT source ID")):
        f.text(110.5, y, label, size=14, bold=True, color=NAVY, centered=True)
    f.text(110.5, 248, "Exact field / source binding", size=11.5, centered=True, max_width=151)

    layers = [
        (334, "1  Endpoint definition", "Name; components; thresholds; target anatomy"),
        (267, "2  Observation rule", "Time anchor; window; visit schedule; repetition"),
        (200, "3  Analysis context", "Patient / lesion unit; denominator; event handling"),
        (133, "4  Estimand evidence", "Population; treatment; outcome; summary; event strategy"),
    ]
    for y, title, body in layers:
        f.box(234, y, 462, 59, fill=PALE_GRAY, stroke=LIGHT, dashed=True)
        f.text(248, y+35, title, size=14.2, bold=True, color=NAVY)
        f.text(248, y+14, body, size=12, color=GRAY, max_width=434)
    # Solid arrows indicate only fields genuinely available from the registry;
    # they do not assert that the richer grey semantic layer has been validated.
    f.arrow([(197, 341), (213, 341), (213, 364), (231, 364)])
    f.arrow([(197, 311), (220, 311), (220, 297), (231, 297)])
    f.box(234, 83, 462, 33, fill=PALE, stroke=TEAL)
    f.text(465, 94, "Field-level provenance: NCT identifier + source pointer", size=12.2, color=TEAL, centered=True, max_width=434)
    f.arrow([(197, 283), (207, 283), (207, 100), (231, 100)])

    f.line(24, 198, 54, 198, color=TEAL)
    f.text(64, 194, "Available input", size=12.3, color=TEAL)
    f.line(24, 175, 54, 175, dashed=True)
    f.text(64, 171, "Untested semantics", size=12.3, color=GRAY)
    f.text(24, 143, "Names and time strings", size=11.8, color=GRAY)
    f.text(24, 127, "are only partial evidence.", size=11.8, color=GRAY)

    f.text(24, 51, "Illustrations only:  patient vs lesion   |   30 days vs 1 year", size=13, bold=True, color=NAVY)
    f.text(24, 29, "Abstract distinctions, not trial labels or observed study results.", size=12, color=GRAY)
    f.finish()


if __name__ == "__main__":
    OUT.mkdir(exist_ok=True)
    architecture()
    endpoint_scope()
```
