## Supplementary data and reproducibility code for "PanVasc Research for AI assisted evidence analysis in panvascular intervention": build_pilot_figures.py

```
"""Render pilot data flow and, only when complete actual results exist, Figures 4-6.

No model outputs, accuracies, intervals or placeholders are invented. Missing
result files skip Figures 4-6 by default; --require-results raises instead.
All figure values come from saved source manifests or the completed test run.
"""
from __future__ import annotations

import argparse
from collections import Counter
import hashlib
import json
from pathlib import Path

from reportlab.lib.colors import HexColor, white
from reportlab.pdfbase.pdfmetrics import stringWidth

from build_concept_figures import Figure, GRAY, INK, LIGHT, NAVY, PALE, PALE_GRAY, TEAL

ROOT = Path(__file__).resolve().parents[1]
PILOT = ROOT / "experiments" / "pilot"
RUNS = PILOT / "runs"
ARM_KEYS = (("generic", "raw"), ("generic", "guard"), ("checklist", "raw"), ("checklist", "guard"))
ARM_LABELS = ("Generic raw", "Generic + guard", "Checklist raw", "Checklist + guard")
ARM_COLORS = (NAVY, HexColor("#8198A8"), TEAL, HexColor("#81BABC"))
CONDITIONS = ("intact", "missing_source", "missing_endpoint", "missing_time")


def read_json(path):
    return json.loads(Path(path).read_text(encoding="utf-8"))


def read_rows(path):
    return [json.loads(line) for line in Path(path).read_text(encoding="utf-8").splitlines() if line.strip()]


def sha(path):
    return hashlib.sha256(Path(path).read_bytes()).hexdigest()


def require(condition, message):
    if not condition:
        raise ValueError(message)


def rate(value, context):
    require(isinstance(value, (float, int)) and not isinstance(value, bool), f"Non-numeric {context}")
    value = float(value)
    require(0 <= value <= 1, f"Out-of-range {context}: {value}")
    return value


def arm(summary, benchmark, prompt, guard):
    key = f"{benchmark}/{prompt}/{guard}"
    require(key in summary["arms"], f"Missing observed arm: {key}")
    data = summary["arms"][key]
    require(data["n"] > 0 and 0 <= data["correct"] <= data["n"], f"Invalid arm denominator: {key}")
    require(abs(rate(data["accuracy"], key) - data["correct"] / data["n"]) < 1e-9, f"Accuracy/count mismatch: {key}")
    ci = data["cluster_accuracy"]
    estimate = rate(ci["estimate"], key + " bootstrap estimate")
    require(abs(estimate-data["accuracy"]) < 1e-9, f"Bootstrap estimate differs from plotted accuracy: {key}")
    require(len(ci["ci95"]) == 2 and ci["clusters"] > 0, f"Missing actual cluster CI: {key}")
    low, high = [rate(v, key + " interval") for v in ci["ci95"]]
    require(low <= high, f"Reversed interval: {key}")
    return data


def rect(f, x, y, width, height, color, *, stroke=None):
    f.c.saveState()
    f.c.setFillColor(color)
    f.c.setStrokeColor(stroke or color)
    f.c.setLineWidth(.5)
    f.c.rect(x, y, width, height, fill=1, stroke=int(stroke is not None))
    f.c.restoreState()


def legend_item(f, x, y, label, color, size=12):
    rect(f, x, y-1, 11, 11, color)
    f.text(x+18, y, label, size=size)


def y_axis(f, left, bottom, width, height, *, percent=True):
    for value in (0, .25, .5, .75, 1):
        y = bottom + value * height
        f.line(left, y, left+width, y, color=HexColor("#DDE3E6"))
        label = f"{value*100:g}" if percent else f"{value:g}"
        f.text(left-8-stringWidth(label, "Helvetica", 11.5), y-4, label, size=11.5, color=GRAY)
    f.line(left, bottom, left, bottom+height, color=LIGHT)


def error_bar(f, x, ylow, yhigh, *, color=INK, cap=4):
    f.line(x, ylow, x, yhigh, color=color)
    f.line(x-cap, ylow, x+cap, ylow, color=color)
    f.line(x-cap, yhigh, x+cap, yhigh, color=color)


def load_data_flow():
    registry_latest = read_json(ROOT / "experiments/registry_grounding/latest_source_run.json")
    source_dir = Path(registry_latest["path"])
    summary = read_json(source_dir / "summary.json")
    manifest = read_json(source_dir / "frozen_manifest.json")
    require(sha(source_dir / "frozen_manifest.json") == registry_latest["manifest_sha256"], "Registry source manifest hash changed")
    for name, expected in manifest["files_sha256"].items():
        require(sha(source_dir/name) == expected, f"Changed frozen registry file: {name}")
    nli = read_json(ROOT / "experiments/nli4ct/frozen_manifest.json")
    tasks = read_rows(PILOT / "data/test_inputs.jsonl")
    development = read_rows(PILOT / "data/development_inputs.jsonl")
    prep = read_json(PILOT / "data/preparation_manifest.json")
    for name, expected in prep["hashes"].items():
        require(sha(ROOT / Path(name)) == expected, f"Changed frozen pilot input: {name}")
    counts = Counter(t["condition"] for t in tasks if t["benchmark"] == "registry")
    require(set(counts) == set(CONDITIONS) | {"clean"}, "Unexpected registry task condition set")
    primary = sum(counts[c] for c in CONDITIONS)
    require(all(counts[c] == summary["test_records"] for c in CONDITIONS), "Registry condition denominators differ")
    selected_nli = nli["selected_summary"]
    require(len(summary["counts_by_domain"]) == 5, "Registry design does not have five query strata")
    per_stratum = {v["test"] for v in summary["counts_by_domain"].values()}
    require(len(per_stratum) == 1, "Registry strata have unequal test counts; update the flow label")
    label_counts = selected_nli["test"]["labels"]
    require(set(label_counts) == {"Entailment", "Contradiction"} and len(set(label_counts.values())) == 1,
            "NLI reference labels are not balanced; update the flow label")
    require(sum(t["benchmark"] == "nli4ct" for t in tasks) == selected_nli["test"]["count"], "NLI test count mismatch")
    require(sum(t["benchmark"] == "nli4ct" for t in development) == selected_nli["dev"]["count"], "NLI development count mismatch")
    return {"registry": summary, "eligible": sum(v["eligible"] for v in summary["counts_by_domain"].values()),
            "primary_tasks": primary, "clean_tasks": counts["clean"], "nli": nli,
            "per_stratum": next(iter(per_stratum)), "per_label": label_counts["Entailment"]}


def figure3():
    data = load_data_flow()
    registry, nli = data["registry"], data["nli"]
    ns = nli["selected_summary"]
    clusters = nli["cluster_summary"]["test"]["connected_components"]
    f = Figure("Figure_3_data_flow", 474)
    f.c.setSubject("Actual frozen pilot data preparation counts; no model performance")
    f.text(24, 442, "Pilot data and evaluation units", size=21, bold=True, color=NAVY)
    f.text(24, 419, "Frozen source preparation; controlled deletions are task variants", size=13, color=GRAY)
    f.box(24, 92, 324, 304, fill=PALE_GRAY, stroke=LIGHT)
    f.box(372, 92, 324, 304, fill=PALE_GRAY, stroke=LIGHT)
    f.text(186, 372, "Registry source fidelity", size=15.5, bold=True, color=TEAL, centered=True)
    f.text(534, 372, "NLI4CT external semantics", size=15.5, bold=True, color=TEAL, centered=True)

    f.box(44, 316, 284, 39, fill=white)
    f.text(186, 330, f"{registry['unique_candidates']} unique candidates", size=14.5, bold=True, centered=True)
    f.arrow([(186, 316), (186, 305)])
    f.box(44, 262, 284, 40, fill=white)
    f.text(186, 276, f"{data['eligible']} mechanically eligible", size=14.5, bold=True, centered=True)
    f.line(186, 262, 186, 251, color=TEAL)
    f.line(108, 251, 264, 251, color=TEAL)
    f.arrow([(108, 251), (108, 239)])
    f.arrow([(264, 251), (264, 239)])
    f.box(44, 189, 128, 47, fill=white)
    f.box(192, 189, 136, 47, fill=PALE)
    f.text(108, 216, f"{registry['development_records']} development", size=12.5, bold=True, centered=True)
    f.text(108, 200, "records", size=12.2, centered=True)
    f.text(260, 216, f"{registry['test_records']} test records", size=13.5, bold=True, centered=True)
    f.text(260, 200, f"{data['per_stratum']} per query stratum", size=11.5, centered=True)
    f.arrow([(260, 189), (260, 176)])
    f.box(44, 110, 284, 63, fill=white)
    f.text(186, 151, f"{data['primary_tasks']} primary tasks", size=14, bold=True, centered=True)
    f.text(186, 133, f"{registry['test_records']} targets x {len(CONDITIONS)} controlled conditions", size=12, centered=True)
    f.text(186, 116, f"+ {data['clean_tasks']} clean single-source controls", size=12, centered=True)

    f.box(392, 316, 284, 39, fill=white)
    f.text(534, 330, "Official source and reference labels", size=13.4, bold=True, centered=True)
    f.line(534, 316, 534, 302, color=TEAL)
    f.line(456, 302, 612, 302, color=TEAL)
    f.arrow([(456, 302), (456, 290)])
    f.arrow([(612, 302), (612, 290)])
    f.box(392, 240, 128, 47, fill=white)
    f.box(540, 240, 136, 47, fill=PALE)
    f.text(456, 267, f"{ns['dev']['count']} development", size=12.5, bold=True, centered=True)
    f.text(456, 251, "statements", size=12.2, centered=True)
    f.text(608, 267, f"{ns['test']['count']} test statements", size=13.1, bold=True, centered=True)
    f.text(608, 251, f"{data['per_label']} per reference label", size=11.5, centered=True)
    f.arrow([(608, 240), (608, 222)])
    f.box(392, 110, 284, 109, fill=white)
    f.text(534, 195, f"{ns['test']['unique_nct_ids']} unique NCT identifiers", size=14, bold=True, centered=True)
    f.text(534, 172, f"{clusters} shared-NCT clusters", size=14, bold=True, centered=True)
    f.text(534, 146, "Connected components of shared sources", size=12, centered=True)
    f.text(534, 126, "Existing labels; no new expert annotation", size=12, centered=True)

    f.text(24, 63, "Registry: five query strata, not independently certified vascular diagnoses.", size=12, color=GRAY)
    f.text(24, 41, "Variants sharing sources are not independent observations.", size=12, color=GRAY)
    f.text(24, 19, "Sampling and data integrity do not establish model accuracy or clinical validity.", size=11.5, color=GRAY)
    f.finish()


def load_results():
    needed = [RUNS/"test_summary.json", RUNS/"test_scored.jsonl", RUNS/"test_responses.jsonl", PILOT/"execution_freeze.json"]
    missing = [str(p.relative_to(ROOT)) for p in needed if not p.is_file()]
    if missing:
        raise FileNotFoundError("Actual completed test artifacts unavailable: " + ", ".join(missing))
    summary = read_json(needed[0])
    require(summary["split"] == "test", "Results do not describe the test split")
    require(summary["response_sha256"] == sha(needed[2]), "Test response hash differs from summary")
    require(summary["execution_freeze_sha256"] == sha(needed[3]), "Execution freeze hash differs from summary")
    inputs = read_rows(PILOT/"data/test_inputs.jsonl")
    require(summary["inference_calls"] == 2 * len(inputs), "Summary does not represent a complete two-arm test run")
    scored = read_rows(needed[1])
    require(len(scored) == 4 * len(inputs), "Scored output count is incomplete")
    seen = {(s["id"], s["arm"], s["guard"]) for s in scored}
    expected = {(t["id"], a, g) for t in inputs for a in ("generic", "checklist") for g in (False, True)}
    require(len(seen) == len(scored) and seen == expected, "Duplicate, missing or unexpected scored output")
    for benchmark in ("registry", "nli4ct"):
        for prompt, guard in ARM_KEYS:
            d = arm(summary, benchmark, prompt, guard)
            subset = [s for s in scored if s["benchmark"] == benchmark and s["arm"] == prompt
                      and s["guard"] == (guard == "guard") and s.get("condition") != "clean"]
            require(len(subset) == d["n"] and sum(s["correct"] for s in subset) == d["correct"], "Scored output/summary disagreement")
    return summary, scored


def figure4(summary):
    f = Figure("Figure_4_registry_results", 538)
    f.c.setSubject("Observed registry task accuracy from complete saved test run")
    f.text(24, 506, "Registry source fidelity", size=21, bold=True, color=NAVY)
    f.text(24, 483, "Exact output tuple accuracy; four-condition primary task", size=13, color=GRAY)
    f.text(24, 450, "A  Raw accuracy by condition", size=14, bold=True, color=NAVY)
    legend_item(f, 427, 450, "Generic", NAVY)
    legend_item(f, 556, 450, "Checklist", TEAL)
    left, bottom, width, height = 58, 276, 630, 152
    y_axis(f, left, bottom, width, height)
    raw = [arm(summary, "registry", a, "raw") for a in ("generic", "checklist")]
    names = (("Intact", ""), ("Source", "absent"), ("Endpoint", "absent"), ("Timeframe", "absent"))
    for index, condition in enumerate(CONDITIONS):
        center = left + (index+.5) * width/4
        for j, data in enumerate(raw):
            d = data["conditions"][condition]
            value = rate(d["accuracy"], condition)
            require(d["n"] > 0 and abs(value-d["correct"]/d["n"]) < 1e-9, "Condition score/count mismatch")
            x = center-34+j*36
            rect(f, x, bottom, 30, value*height, (NAVY, TEAL)[j])
            f.text(x+15, bottom+value*height+6, f"{100*value:.0f}", size=11.5, centered=True)
        f.text(center, 254, names[index][0], size=12, centered=True)
        if names[index][1]:
            f.text(center, 238, names[index][1], size=12, centered=True)
    f.text(24, 214, "Percent correct; each condition retains its full test denominator.", size=11.5, color=GRAY)
    f.text(24, 190, "B  Overall accuracy and cluster bootstrap 95% CI", size=14, bold=True, color=NAVY)
    forest_left, forest_width = 190, 326
    for v in (0, .5, 1):
        x = forest_left + forest_width*v
        f.line(x, 61, x, 173, color=HexColor("#E0E5E8"))
        f.text(x, 44, f"{100*v:g}", size=11.5, color=GRAY, centered=True)
    for i, ((prompt, guarded), label, color) in enumerate(zip(ARM_KEYS, ARM_LABELS, ARM_COLORS)):
        d = arm(summary, "registry", prompt, guarded)
        ci = d["cluster_accuracy"]["ci95"]
        y = 166-i*29
        f.text(24, y-4, label, size=12)
        lo, hi = [forest_left+forest_width*v for v in ci]
        f.line(lo, y, hi, y, color=color)
        f.line(lo, y-4, lo, y+4, color=color)
        f.line(hi, y-4, hi, y+4, color=color)
        f.c.setFillColor(color)
        f.c.circle(forest_left+forest_width*d["accuracy"], y, 4, fill=1, stroke=0)
        f.text(537, y-4, f"{100*d['accuracy']:.1f} [{100*ci[0]:.1f}, {100*ci[1]:.1f}]", size=11.5)
    baseline = summary["baseline"]["registry_rule_parser"]
    require(baseline["n"] == raw[0]["n"], "Rule baseline denominator differs from primary task")
    rate_value = rate(baseline["correct"]/baseline["n"], "rule-only baseline")
    clusters = summary["counts"]["registry"]["clusters"]
    f.text(24, 18, f"Rule parser: {baseline['correct']}/{baseline['n']} ({100*rate_value:.1f}%). CI clusters: {clusters} target NCTs.", size=11.5, color=GRAY)
    f.finish()


def figure5(summary):
    f = Figure("Figure_5_nli_results", 501)
    f.c.setSubject("Observed NLI4CT label accuracy and evidence micro F1")
    baseline = summary["baseline"]["nli_majority"]
    labels = baseline["label_counts"]
    f.text(24, 469, "External clinical-trial inference", size=21, bold=True, color=NAVY)
    f.text(24, 446, f"NLI4CT: {labels['Entailment']} Entailment / {labels['Contradiction']} Contradiction reference labels", size=13, color=GRAY)
    f.text(24, 416, "A  Label accuracy (%)", size=14.5, bold=True, color=NAVY)
    f.text(385, 416, "B  Evidence micro-F1", size=14.5, bold=True, color=NAVY)
    bottom, height, width = 174, 217, 270
    for left, percent in ((56, True), (415, False)):
        y_axis(f, left, bottom, width, height, percent=percent)
    for i, ((prompt, guard), color) in enumerate(zip(ARM_KEYS, ARM_COLORS)):
        data = arm(summary, "nli4ct", prompt, guard)
        accuracy = data["accuracy"]
        evidence = rate(data["evidence"]["f1"], "evidence micro F1")
        tp, fp, fn = [data["evidence"][k] for k in ("tp", "fp", "fn")]
        expected = 2*tp/(2*tp+fp+fn) if 2*tp+fp+fn else 0
        require(abs(evidence-expected) < 1e-9, "Evidence F1/count mismatch")
        ci = data["cluster_accuracy"]["ci95"]
        for left, value in ((56, accuracy), (415, evidence)):
            center = left+(i+.5)*width/4
            rect(f, center-18, bottom, 36, value*height, color)
            if left == 56:
                error_bar(f, center, bottom+ci[0]*height, bottom+ci[1]*height)
                top = max(value, ci[1])
                label = f"{100*value:.0f}"
            else:
                top, label = value, f"{value:.2f}"
            f.text(center, bottom+top*height+6, label, size=11.5, centered=True)
            f.text(center, 153, "G" if prompt == "generic" else "C", size=12.5, bold=True, centered=True)
            f.text(center, 136, "raw" if guard == "raw" else "guard", size=11.5, centered=True)
    f.text(24, 107, "G: generic prompt   |   C: checklist prompt", size=12, color=GRAY)
    f.text(24, 84, "Guarded answers reuse the same model responses; rejected answers count as incorrect.", size=11.5, color=GRAY)
    clusters = summary["counts"]["nli4ct"]["clusters"]
    f.text(24, 61, f"Accuracy CIs: {clusters} shared-NCT clusters. Evidence F1 is shown without a CI.", size=11.5, color=GRAY)
    majority = baseline["correct"]/baseline["n"]
    f.text(24, 32, f"Label-only majority baseline: {100*majority:.1f}% ({baseline['correct']}/{baseline['n']}); no evidence score.", size=12, color=GRAY)
    f.finish()


def guard_breakdown(summary, scored):
    primary = [s for s in scored if s.get("condition") != "clean"]
    keyed = {(s["benchmark"], s["arm"], s["id"], s["guard"]): s for s in primary}
    counts = {}
    for benchmark in ("registry", "nli4ct"):
        for prompt in ("generic", "checklist"):
            raw = [s for s in primary if s["benchmark"] == benchmark and s["arm"] == prompt and s["guard"] is False]
            result = [0, 0, 0, 0]  # accepted-correct, accepted-wrong, rejected-raw-correct, rejected-raw-wrong
            for original in raw:
                guarded = keyed[(benchmark, prompt, original["id"], True)]
                accepted = original["contract_accepted"]
                require(type(accepted) is bool and type(original["correct"]) is bool, "Guard category lacks boolean state")
                require(accepted == guarded["contract_accepted"], "Contract acceptance changed across raw/guard records")
                require(guarded["covered"] == accepted, "Guard coverage is not contract acceptance")
                if accepted:
                    require(guarded["correct"] == original["correct"], "Guard changed an accepted answer's correctness")
                else:
                    require(guarded["correct"] is False, "Rejected answer unexpectedly scored correct")
                result[(0 if accepted else 2) + (0 if original["correct"] else 1)] += 1
            guard_arm = arm(summary, benchmark, prompt, "guard")
            raw_arm = arm(summary, benchmark, prompt, "raw")
            require(sum(result) == raw_arm["n"] == guard_arm["n"], "Guard category denominator mismatch")
            require(result[0] == guard_arm["correct"] and result[0]+result[2] == raw_arm["correct"], "Guard category accuracy mismatch")
            require(abs((result[0]+result[1])/sum(result)-guard_arm["coverage"]) < 1e-9, "Guard coverage mismatch")
            counts[(benchmark, prompt)] = result
    return counts


def figure6(summary, scored):
    groups = guard_breakdown(summary, scored)
    f = Figure("Figure_6_guard_boundary", 511)
    f.c.setSubject("Observed paired guard acceptance/rejection and original-answer correctness")
    colors = (TEAL, HexColor("#B66765"), HexColor("#D6A561"), HexColor("#B7C1C8"))
    f.text(24, 479, "Guard acceptance and rejection trade-offs", size=21, bold=True, color=NAVY)
    f.text(24, 456, "Paired raw and guarded answers; every task remains in the denominator", size=12.8, color=GRAY)
    legend_item(f, 24, 425, "Accepted + correct (AC)", colors[0], 12.2)
    legend_item(f, 378, 425, "Accepted + wrong (AW)", colors[1], 12.2)
    legend_item(f, 24, 402, "Rejected + raw correct (RC)", colors[2], 12.2)
    legend_item(f, 378, 402, "Rejected + raw wrong (RW)", colors[3], 12.2)
    left, width, bar_height = 194, 492, 25
    for value in (0, .25, .5, .75, 1):
        x = left+width*value
        f.line(x, 125, x, 355, color=HexColor("#E5E9EC"))
        f.text(x, 83, f"{100*value:g}%", size=11.5, color=GRAY, centered=True)
    positions = (("registry", "generic", 330), ("registry", "checklist", 274),
                 ("nli4ct", "generic", 181), ("nli4ct", "checklist", 125))
    f.text(24, 371, "Registry primary tasks", size=14, bold=True, color=NAVY)
    f.text(24, 222, "NLI4CT statements", size=14, bold=True, color=NAVY)
    for benchmark, prompt, y in positions:
        values = groups[(benchmark, prompt)]
        n = sum(values)
        f.text(24, y+8, f"{prompt.capitalize()} (n={n})", size=12.5)
        x = left
        for value, color in zip(values, colors):
            segment = value/n*width
            if value:
                rect(f, x, y, segment, bar_height, color)
                if segment >= 34:
                    f.text(x+segment/2, y+8, str(value), size=11.5, color=white if color == TEAL else NAVY, centered=True)
            x += segment
        f.text(left, y-17, f"AC {values[0]}  |  AW {values[1]}  |  RC {values[2]}  |  RW {values[3]}", size=11.5, color=GRAY)
    f.text(24, 53, "Correctness means the complete registry tuple or the NLI reference label.", size=11.5, color=GRAY)
    f.text(24, 31, "Registry clean controls are excluded. Source checks cannot establish semantic support.", size=11.5, color=GRAY)
    f.finish()


def main():
    parser = argparse.ArgumentParser(description=__doc__)
    parser.add_argument("--figures", nargs="+", type=int, choices=(3, 4, 5, 6), default=[3, 4, 5, 6])
    parser.add_argument("--require-results", action="store_true")
    args = parser.parse_args()
    if 3 in args.figures:
        figure3()
    requested = [number for number in args.figures if number in (4, 5, 6)]
    if not requested:
        return
    try:
        summary, scored = load_results()
    except FileNotFoundError as error:
        if args.require_results:
            raise
        print(f"SKIP Figures {','.join(map(str, requested))}: {error}. No placeholder figures are generated.")
        return
    for number in requested:
        if number == 4:
            figure4(summary)
        elif number == 5:
            figure5(summary)
        elif number == 6:
            figure6(summary, scored)


if __name__ == "__main__":
    main()
```
