## Supplementary data and reproducibility code for "PanVasc Research for AI assisted evidence analysis in panvascular intervention": start_local_model.ps1

```
[CmdletBinding()]
param(
    [ValidateRange(1024, 65535)][int]$Port = 18123,
    [ValidateRange(512, 32768)][int]$ContextSize = 8192,
    [ValidateRange(0, 999)][int]$GpuLayers = 99,
    [ValidateRange(1, 128)][int]$Threads = 8
)
$ErrorActionPreference = 'Stop'
$workspaceRoot = Split-Path -Parent $PSScriptRoot
$runtimeDir = Join-Path $workspaceRoot 'runtime\local_llm'
$manifest = Get-Content -LiteralPath (Join-Path $runtimeDir 'setup_manifest.json') -Raw | ConvertFrom-Json
$serverPath = $manifest.runtime.server_executable
$modelPath = $manifest.model.download.path
if (-not $serverPath -or -not (Test-Path -LiteralPath $serverPath)) { throw 'Verified llama-server executable is unavailable.' }
if (-not $modelPath -or -not (Test-Path -LiteralPath $modelPath)) { throw 'Verified model file is unavailable.' }
if (-not $manifest.model.download.verified -or -not $manifest.runtime.download.verified) { throw 'Setup SHA256 verification is incomplete.' }
if (@(Get-NetTCPConnection -LocalPort $Port -State Listen -ErrorAction SilentlyContinue).Count -gt 0) { throw "Port $Port already has a listener." }
$timestamp = (Get-Date).ToUniversalTime().ToString('yyyyMMddTHHmmssfffZ')
$stdout = Join-Path $runtimeDir "server-$timestamp.stdout.log"
$stderr = Join-Path $runtimeDir "server-$timestamp.stderr.log"
$serverArguments = @('--model', ('"' + $modelPath + '"'), '--alias', 'Qwen3-4B-Q8_0', '--host', '127.0.0.1', '--port', $Port.ToString(), '--ctx-size', $ContextSize.ToString(), '--parallel', '1', '--gpu-layers', $GpuLayers.ToString(), '--threads', $Threads.ToString(), '--jinja', '--no-context-shift')
$serverProcess = Start-Process -FilePath $serverPath -ArgumentList $serverArguments -WorkingDirectory (Split-Path -Parent $serverPath) -WindowStyle Hidden -RedirectStandardOutput $stdout -RedirectStandardError $stderr -PassThru
$record = [ordered]@{
    started_utc = (Get-Date).ToUniversalTime().ToString('o')
    pid = $serverProcess.Id
    executable = $serverPath
    model = $modelPath
    arguments = $serverArguments
    host = '127.0.0.1'
    port = $Port
    base_url = "http://127.0.0.1:$Port/v1"
    health_url = "http://127.0.0.1:$Port/health"
    context_size = $ContextSize
    parallel = 1
    stdout_log = $stdout
    stderr_log = $stderr
    status = 'starting'
}
$recordPath = Join-Path $runtimeDir 'active_server.json'
$record | ConvertTo-Json -Depth 6 | Set-Content -LiteralPath $recordPath -Encoding utf8
$record | ConvertTo-Json -Depth 6
```
