## Supplementary data and reproducibility code for "PanVasc Research for AI assisted evidence analysis in panvascular intervention": stop_local_model.ps1

```
[CmdletBinding()]
param()
$ErrorActionPreference = 'Stop'
$workspaceRoot = Split-Path -Parent $PSScriptRoot
$recordPath = Join-Path $workspaceRoot 'runtime\local_llm\active_server.json'
if (-not (Test-Path -LiteralPath $recordPath)) { throw 'No recorded local model process.' }
$record = Get-Content -LiteralPath $recordPath -Raw | ConvertFrom-Json
$serverProcess = Get-Process -Id $record.pid -ErrorAction SilentlyContinue
if ($serverProcess) {
    if ($serverProcess.Path -ne $record.executable) { throw 'Recorded PID now belongs to another executable; refusing to stop it.' }
    $expectedStart = [datetime]::Parse($record.started_utc).ToUniversalTime()
    if ([math]::Abs(($serverProcess.StartTime.ToUniversalTime() - $expectedStart).TotalSeconds) -gt 15) { throw 'Recorded PID start time differs; refusing to stop it.' }
    Stop-Process -Id $record.pid
}
$record.status = 'stopped'
$record | Add-Member -NotePropertyName stopped_utc -NotePropertyValue ((Get-Date).ToUniversalTime().ToString('o')) -Force
$record | ConvertTo-Json -Depth 6 | Set-Content -LiteralPath $recordPath -Encoding utf8
Write-Output "Local model process $($record.pid) is stopped."
```
