## Supplementary data and reproducibility code for "PanVasc Research for AI assisted evidence analysis in panvascular intervention": README.md

# Local model pilot reproducibility

The manuscript's new local-model study is distinct from the earlier 28-test engineering demonstration. The fixed model, inputs, prompts, guards and analysis plan are documented in the execution freeze and manifests. Test numerical scores retain every task and both prompt arms with and without the same-output guard. Malformed, rejected and transport-failed cases are not removed.

## Materials deliberately omitted

This package does not contain clinical-trial text, NLI4CT's source archive, complete model responses that may reproduce source text, article abstracts, model weights, authentication material or signed download URLs. Obtain NLI4CT from its official pinned revision and Qwen weights from the official model publisher. Registry identifiers and hashes are supplied. Live registry records are mutable; re-fetching them may not reproduce the historical local snapshot. A hash is not a replacement for unavailable source text.

## Reproduction sequence

Use the original project root layout under code. Review the acquisition READMEs and frozen protocol before running. Obtain the official source material, prepare the fixed splits, then run experiments/pilot/run_pilot.py prepare, verify_pilot.py, and run_pilot.py freeze. Start the specified local model server on 127.0.0.1:18123 before run_pilot.py run --split test, then run_pilot.py evaluate --split test. The release freeze is an audit record of the reported run; an independent reproduction creates its own timestamp and records any changed source hashes. Do not silently overwrite or blend freezes.

The rule baseline verifies only source-field fidelity. NLI4CT is a breast-cancer benchmark with historical expert annotations, not new vascular annotation. Public-data pretraining exposure cannot be excluded. No model weights were trained in this study.
