## Supplementary data and reproducibility code for "PanVasc Research for AI assisted evidence analysis in panvascular intervention": REPRODUCIBILITY.md

# PanVasc Research reproducibility supplement

This supplement documents two distinct evaluations conducted on 19 September 2026: the historical software feasibility demonstration described below, and the completed local-model pilot documented separately under `pilot/`. The historical 28 engineering tests are not model-inference tests. The later pilot made 700 test inference calls using fixed official Qwen3-4B Q8_0 weights. Neither evaluation involved model training or patient-level clinical prediction.

The supplement contains no real patient records, article abstracts, raw PubMed XML, raw Europe PMC response bodies, third-party trial text, complete model responses, local research tasks, credentials, or private author correspondence. Public article titles and bibliographic identifiers are supplied for source discovery. No manuscript approval, publication license, or public repository accession is implied by the existence of this local directory.

## Files

| File | Contents |
|---|---|
| `retrieval_runs_without_abstracts.json` | Three distinct live experiments, preserving successful and failed cases, actual or explicitly reconstructed queries, source identity, timestamps, and recorded transport attempts |
| `retrieval_queries_all_runs.csv` | Fifteen domain-level query cases: five per experiment |
| `bibliographic_records_all_runs.csv` | Twenty-four returned-record rows across the successful cases of the two Europe PMC experiments; repeated records across experiments are retained |
| `supplementary_table_S1_final_retrieval.csv` | Fifteen rows from the final Europe PMC experiment, with exact query, execution time, PMID, title, available DOI, and source link |
| `synthetic_example.csv` | The 120 wholly fictitious records used in the numerical demonstration |
| `synthetic_analysis.json` | Original computed summaries and the source CSV hash |
| `independent_numeric_verification.json` | Independent CSV count verification, integer-combination Fisher probability calculation, and Wilson formula comparison |
| `offline_validation.json` / `.csv` | Final 28-case offline acceptance results, test expectations, source hashes, runtime versions, and scope limitations |
| `runtime.json` | Historical engineering-demonstration environment; its `language_model_inference: not_run` applies to that earlier run, not the later pilot |
| `manifest_sha256.json` | Hashes of the accompanying data and documentation files at assembly time |
| `pilot/README.md` | Separate local-model experiment scope, acquisition limits and reproduction sequence |
| `pilot/code/experiments/pilot/runs/test_summary.json` | Actual aggregate results from 700 local-model test calls |
| `pilot/test_numerical_scores.jsonl` | 1,400 numerical scoring rows: raw and guarded scoring of each of the same 700 responses, with third-party text omitted |
| `pilot/model_provenance.json` | Public model/runtime revisions, verified hashes and setup metadata without original machine paths or network settings |

The desktop source distribution is included under `code/`, including the interface, optional remote model adapter, test suite, and reproduction scripts. Run the historical engineering commands below from `code/`. The separate local-model runner and protocol are under `pilot/code/`; do not mix the two code roots or their recorded environments. File hashes in the final offline report identify `panvasc/core.py`, `tests/test_core.py`, and `scripts/run_validation.py`; release manifests additionally identify packaged source and data files. No article abstracts, local workspace, credentials, or raw response files are included.

## Completed local-model pilot

The 700 test calls comprise two fixed prompt arms applied to 350 inputs per arm: 200 registry tasks across four controlled conditions, 50 clean registry controls, and 100 NLI4CT statements. Applying the input-only guard to the same saved responses requires no additional inference. The 10 development-set technical calls and one non-medical JSON smoke test are recorded separately and excluded from these 700 test calls. They do not add test examples or clinical validation.

See [the pilot protocol](pilot/code/experiments/pilot/protocol.md), [actual test summary](pilot/code/experiments/pilot/runs/test_summary.json), and [pilot reproduction guide](pilot/README.md). Model setup and public-package reconstruction are described in [LOCAL_MODEL_SETUP.md](pilot/code/docs/LOCAL_MODEL_SETUP.md). The original machine's model-acquisition helper, smoke-test helper, downloaded binaries, model weights and private setup paths are not packaged. Reproduction requires official downloads, hash verification, a new local setup record, source acquisition and a new execution freeze; the historical freeze must remain an audit record. Live registry records may have changed, so this release does not promise exact recovery of the omitted historical source text.

## What was run in the historical engineering demonstration

1. **Initial NCBI experiment:** five calls to the primary `search_pubmed` API, each with one domain, no extra keywords, years 2015–2026, relevance sorting, and a limit of three records. All five failed to complete because of recorded TLS errors. The peripheral case reached EFetch, showing that its preceding ESearch had completed; this is still a failed end-to-end task.
2. **Initial Europe PMC experiment:** a distinct series of five direct Europe PMC calls, with equivalent domain intent but different title/abstract query semantics and default provider ordering. Three completed: carotid, aortic, and venous. Two failed because of TLS errors.
3. **Retry-enabled Europe PMC experiment:** another distinct five-call series after bounded retry handling was added. All five completed and each returned three records. The recorded attempt arrays contain exactly one HTTP 200 attempt in every case. These observations do not demonstrate that retrying caused the improvement.

The final hit counts were 49,016; 2,659; 8,420; 15,697; and 4,976 for coronary, peripheral, carotid, aortic, and venous queries, respectively. Elapsed seconds were 3.441, 5.291, 5.129, 5.152, and 5.178. Counts are service snapshot counts, not screened eligible studies. The five intents were convenience-selected and adapted during development. Fifteen complete identifiers/titles do not measure relevance or claim support.

The two earlier runs did not record attempt arrays. A missing `transport_attempts` field remains null and must not be interpreted as zero attempts or as proof of one attempt. Failed records did not retain structured query/timing fields. Where a query is recoverable from a recorded error URL, `query_origin` says so. The peripheral failed EFetch URL lacks the preceding search term; its query is explicitly reconstructed from runner defaults and the audited domain configuration. Original initial source snapshots were not frozen, so exact historical implementation replay is unavailable.

The historical adapter labeled returned records `Europe PMC MEDLINE`. This is retained as a recorded software label, not a verified indexing classification. The actual source restriction was `SRC:MED` (PubMed-source records); the experiment did not establish each record's MEDLINE indexing status.

## Offline reproducibility

The reported environment was Python 3.11.9, NumPy 2.4.6, pandas 2.3.3, requests 2.34.2, and SciPy 1.17.1 on Windows. These are the recorded key package versions, not a complete transitive environment lock. A source release containing the original package layout can be checked from its root with:

```text
python -m unittest tests.test_core -v
python scripts/run_validation.py
```

The second command writes new validation results under `artifacts/`; run it in a separate copy to preserve this historical supplement. The tests use mocked network responses and temporary files. The 28 final tests were developed and revised in the same development cycle as the software. Four regression checks were added after review exposed blank identifier, fully missing outcome, and numeric identifier representation gaps; the final passing result followed fixes. This is not a blinded or held-out benchmark.

Numeric reference cases include a two-by-two table `[[1,9],[11,3]]` with two-sided Fisher P = 0.0027594561852200836 and a zero-event, n = 10 Wilson interval with upper bound 0.2775327998628892. Numeric tests use declared tolerances. Sample standard deviation uses n − 1; reported quartiles use the installed pandas default linear interpolation.

## Synthetic demonstration

The CSV contains SYN001–SYN120. A and B each have 60 observations, with event counts 8 and 12 and a fixed 365-day follow-up. Age values were generated with NumPy `default_rng(20260919).integers(40,81,120)`, then zero-based rows 4, 44, and 91 were set to missing. The CSV SHA-256 is `c8182789ac919813e632a5b8cce2ccf345472ee3b24f1bd5fea5bf57e4ac1d3b`.

Reading the CSV independently gives table `[[8,52],[12,48]]`. A two-sided Fisher calculation by summing hypergeometric probabilities no greater than the observed table probability gives P = 0.46315569016131075; the original SciPy output is 0.46315569016131064. Their difference is floating-point rounding. The odds ratio is 0.6153846153846154 and the A-minus-B risk difference is −0.06666666666666668. These numbers describe a constructed fixture and have no clinical interpretation.

## Repeating network requests

From a frozen source release, `python scripts/run_demonstration.py` runs the primary NCBI cases and recreates the synthetic example; `python scripts/run_alternative_retrieval.py` runs the current Europe PMC implementation. Both perform live network requests and write outputs, so use a separate copy. The current alternate-service script writes a retry-run result file; it does not reproduce the historical pre-retry implementation. Requests may return different records, counts, delays, or errors as indexes and networks change.

Use the queries in this supplement to identify the recorded search intent. Raw abstracts and responses remain excluded from this distributable supplement. A hash of an omitted raw response records its identity, but cannot allow a reader to reconstruct or independently hash that omitted content. The manuscript should not claim stronger reproducibility than these retained inputs and outputs permit.

## Boundaries

No language-model inference was performed in the historical retrieval, 28-test engineering and synthetic-data demonstration documented in the preceding sections. That scope-qualified statement and the historical `runtime.json` do not describe the later 700-call local-model pilot. The latter measures registry source-field fidelity and NLI4CT reference-label/evidence agreement; NLI4CT is an external breast-cancer benchmark, not vascular clinical validation.

Neither evaluation performed fine-tuning, weight updates, a real clinician feedback panel, clinical endpoint prediction, or biological validation. Neither establishes systematic-review completeness, literature relevance, exhaustive prompt-injection resistance, clinical benefit, or a panvascular advantage. Citation identity remains distinct from source support. The desktop's optional remote model adapter remains untested by these experiments and requires separate activation and evaluation.
